# Ultra-stable insulin-glucagon fusion protein exploits an endogenous hepatic switch to mitigate hypoglycemic risk

**DOI:** 10.1101/2024.05.20.594997

**Authors:** Nicolas Varas, Rachel Grabowski, Mark A. Jarosinski, Ningwen Tai, Raimund I. Herzog, Faramarz Ismail-Beigi, Yanwu Yang, Alan D. Cherrington, Michael A. Weiss

## Abstract

The risk of hypoglycemia and its serious medical sequelae restrict insulin replacement therapy for diabetes mellitus. Such adverse clinical impact has motivated development of diverse glucose-responsive technologies, including algorithm-controlled insulin pumps linked to continuous glucose monitors (“closed-loop systems”) and glucose-sensing (“smart”) insulins. These technologies seek to optimize glycemic control while minimizing hypoglycemic risk. Here, we describe an alternative approach that exploits an endogenous glucose-dependent switch in hepatic physiology: preferential insulin signaling (under hyperglycemic conditions) *versus* preferential counter-regulatory glucagon signaling (during hypoglycemia). Motivated by prior reports of glucagon-insulin co-infusion, we designed and tested an ultra-stable glucagon-insulin fusion protein whose relative hormonal activities were calibrated by respective modifications; physical stability was concurrently augmented to facilitate formulation, enhance shelf life and expand access. An N-terminal glucagon moiety was stabilized by an α-helix-compatible Lys^13^-Glu^17^ lactam bridge; A C-terminal insulin moiety was stabilized as a single chain with foreshortened C domain. Studies *in vitro* demonstrated (a) resistance to fibrillation on prolonged agitation at 37 °C and (b) dual hormonal signaling activities with appropriate balance. Glucodynamic responses were monitored in rats relative to control fusion proteins lacking one or the other hormonal activity, and continuous intravenous infusion emulated basal subcutaneous therapy. Whereas efficacy in mitigating hyperglycemia was unaffected by the glucagon moiety, the fusion protein enhanced endogenous glucose production under hypoglycemic conditions. Together, these findings provide proof of principle toward a basal glucose-responsive insulin biotechnology of striking simplicity. The fusion protein’s augmented stability promises to circumvent the costly cold chain presently constraining global insulin access.

**Significance Statement:** The therapeutic goal of insulin replacement therapy in diabetes is normalization of blood-glucose concentration, which prevents or delays long-term complications. A critical barrier is posed by recurrent hypoglycemic events that results in short- and long-term morbidities. An innovative approach envisions co-injection of glucagon (a counter-regulatory hormone) to exploit a glycemia-dependent hepatic switch in relative hormone responsiveness. To provide an enabling technology, we describe an ultra-stable fusion protein containing insulin- and glucagon moieties. Proof of principle was obtained in rats. A single-chain insulin moiety provides glycemic control whereas a lactam-stabilized glucagon extension mitigates hypoglycemia. This dual-hormone fusion protein promises to provide a basal formulation with reduced risk of hypoglycemia. Resistance to fibrillation may circumvent the cold chain required for global access.

## Introduction

The goal of insulin replacement therapy (IRT) for diabetes mellitus (DM) is regulation of blood-glucose concentration (BGC) with minimal excursions above or below the normal range (80-100 mg/dL fasting and 120 mg/dL at 2 h postprandial) (1). The advent of continuous glucose monitoring (CGM) has motivated *time in range* (TIR; 70-180 mg/dL) and *time below range* (TBR; <70 mg/dL) as key metrics (1). Although tight glycemic control prevents or delays microvascular complications (and likely macrovascular disease) in Type 1 DM (T1D) (2, 3), the fear of hypoglycemia poses a most significant barrier to its implementation (4). Acute IRT-related hypoglycemia induces progressive adrenergic and neuroglycopenic symptoms (tremulousness, confusion, loss of consciousness, convulsions) (5, 6) whereas chronic repeated episodes can lead to cognitive decline with decreased life span (7, 8). Clinical approaches have traditionally relied on education of patients (and their families) to recognize signs and symptoms of hypoglycemia, enabling prompt intervention: ingestion of glucose-rich substances in mild cases to subcutaneous (SQ) injection of glucagon when severe (7, 9, 10). Education alone is often ineffective, especially in patients in whom hypoglycemia-induced autonomic failure (11) attenuates counterregulatory responses, in turn leading to hypoglycemia unawareness (12). The present study envisions a physiology-based biotechnology to mitigate hypoglycemic risk without compromising overall glycemic control (13).

Diverse diabetes technologies have been explored to reduce hypoglycemic risk and enhance TIR (14, 15). These include (i) mechanical closed-loop systems with CGM-based control of insulin- or bihormonal insulin-glucagon pumps (16, 17) (Fig. 1A); (ii) glucose-responsive polymers for hormone encapsulation to effect glucose-dependent release from a SQ depot or microneedles (18–20) (Fig. 1B); (iii) glucose-dependent bioavailability or clearance of an insulin analog modified to bind to an endogenous macromolecule (Fig. 1C) (21–23); and (iv) intrinsic glucose-responsive insulin (GRI) analogs (Fig. 1D) (24). These approaches share a requirement for measuring or sensing glucose concentrations in interstitial fluid or blood. *We sought to circumvent this requirement by exploiting a potential physiological switch in hepatocytes:* glycemia-dependent regulation of opposing insulin-*versus*-glucagon signaling, due to cross-sensitization between the hormones (25–27). This strategy exploited “yin-yang” endocrine mechanisms: under hyperglycemic conditions insulin inhibits glucagon action (28), whereas glucagon predominates under hypoglycemia (25) (Fig. 2A)—even in the face of a 35-fold increase in insulin concentration (29). Indeed, glucagon-stimulated hepatic glucose output has been observed to mitigate hypoglycemia following concurrent infusion of the two hormones at fixed molar ratios (30–32). Despite the elegance of this idea, its application is presently limited by the hormones’ discordant pharmacokinetic/pharmacodynamic (PK/PD) properties (33, 34) and incompatible formulation chemistries (31, 35), necessitating separate SQ depots. In addition, formulation of glucagon is complicated by its marked physical instability (36, 37). In the case of homologous peptide hormones and cognate receptors (as exemplified by glucagon and related incretins; for review, see (38)), these challenges have been overcome via chimeric multi-agonist peptides (39, 40).

**Figure 1.**
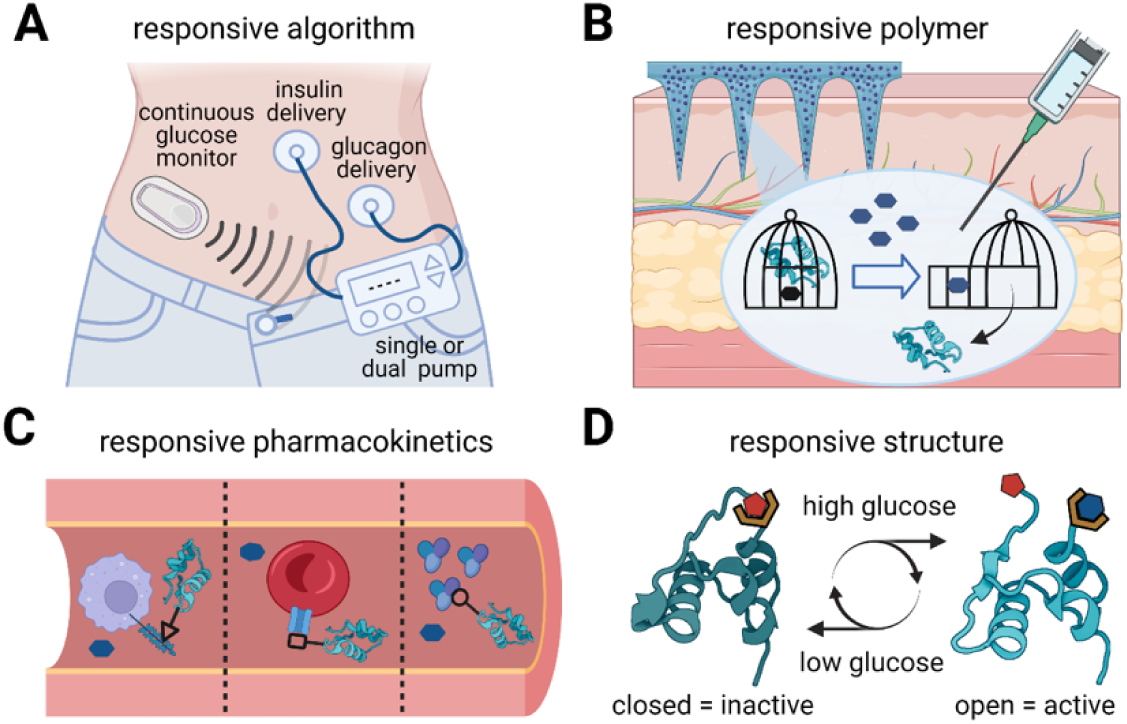
Glucose-responsive systems. (A) Mechanical closed-loop systems consist of a CGM that sends glucose data to a single-(insulin) or dual hormone (insulin-glucagon) pump, which in turn delivers the hormones based on a responsive algorithm. (B) Glucose-responsive polymers for hormone encapsulation to effect glucose-dependent release from a SQ depot or microneedles. Closed and open cages provide a representation of different strategies used to trap insulin (or glucagon). (C) Glucose-dependent binding to a macromolecule that in turn regulates bioavailability or clearance. From left to right, binding to mannose receptor, GLUT1, or albumin. (D) Intrinsic glucose-responsive switch within an insulin analog that opens (activates) on binding glucose, creating a conformational switch. Images were created with Biorender.com.

**Figure 2.**
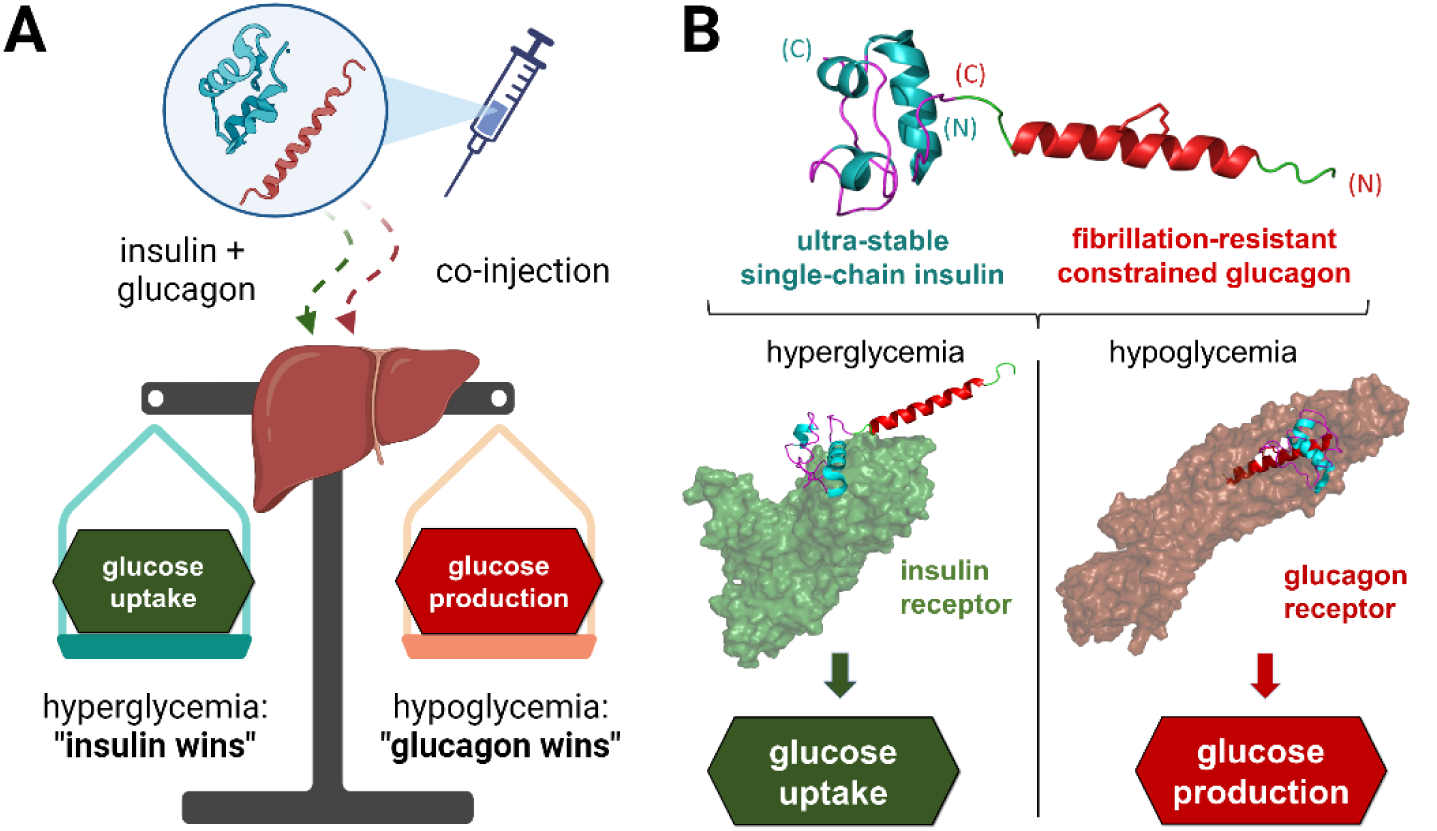
Physiological glucose-responsive “switch.ℍ. (A) Co-injection of insulin and glucagon exploits a physiological switch in hepatic hormone responsiveness. Under hyperglycemic conditions, insulin activity prevails, whereas glucagon overpowers insulin under hypoglycemic conditions. (B) Fibrillation-resistant FP exploits the same physiological switch via an ultra-stable single polypeptide. In respective hormone-receptor complexes insulin exhibit a free N terminus (Phe^B1^), and glucagon a free C terminus (H^1^). SCI in FP is depicted based on NMR structure of homologous analog (PDB ID: 5WBT). Glucagon analog in FP is depicted based on crystal structure of native glucagon (as an α-helical trimer; PDB ID: 1GCN). Images were created with Biorender.com.

Can the dual-hormone approach be generalized to non-homologous signaling systems? To investigate this possibility, we undertook the design, synthesis and characterization of a novel glucagon-insulin fusion protein (FP). The envisioned single-chain FP would provide respective insulin- and glucagon activities in appropriate balance with similar bioavailability and time-action profiles. To enable dual hormonal activities, the proposed scheme exploited the distinct structures of the glucagon receptor (a G-protein-coupled receptor; GPCR) (41) and insulin receptor (IR; a transmembrane tyrosine kinase) (42, 43), whose respective mechanisms of hormone binding motivated an *N-terminal* glucagon domain flexibly linked to a *C-terminal* insulin domain (41–43) (Fig. 2B). To prevent fibrillation, the glucagon moiety was protected by a central [i, i+4] side-chain lactam bridge (44, 45), and the insulin moiety by an engineered connecting (C) peptide between B- and A chains (46, 47). Given the intrinsic 1:1 stoichiometry of the FP, relative hormonal activities were calibrated by amino acid substitutions in each domain (30–32). Following biophysical and cell-based studies, the FP was evaluated in rats by continuous intravenous (IV) infusion to emulate the PK/PD profile of a basal insulin analog. Strikingly, hyperinsulinemic clamps demonstrated predominance of insulin action under hyperglycemic conditions, whereas glucagon signaling enhanced endogenous glucose production under hypoglycemic conditions thereby mitigating insulin action. Structural independence of the dual hormonal domains was verified by heteronuclear NMR spectroscopy.

Together, our results provide proof of concept for a fibrillation-resistant GRI biotechnology remarkable for its molecular simplicity. An ultra-stable glucagon-insulin FP, compatible with the structures of respective cognate receptors (41–43), promises to enhance the safety of insulin therapy in both affluent societies and the developing world.

## Results

Our study had three parts. We first designed a glucagon-insulin FP based on enzymatic ligation of ultra-stable hormone analogs. Functional properties were next evaluated *in vitro*, in human cell culture and in rats. Hyperinsulinemic clamps with defined glycemic set-points were employed to demonstrate endogenous glucose-dependent metabolic control on continuous IV infusion. Finally, biophysical studies were undertaken to validate structural design assumptions.

### Overview of peptide design

To mitigate the shared propensities of glucagon and insulin to form inactive fibrils, we sought molecular tactics to stabilize the respective hormones with an appropriate balance of hormone-specific activities. Control FPs were constructed by mixing and matching active and inactive hormone segments. For clarity, unless otherwise indicated, glucagon residues are designated in single-letter code whereas insulin residues are designated in three-letter code. To facilitate comparison between respective component hormones and FPs (89 residues), polypeptides positions (superscripts) are designated in relation to the glucagon moiety (residues 1-29 in FPs), tripeptide linker (residues 30-32), and successive insulin segments (B1-B30, C1-C6 and A1-A21). Aside from C-terminal linker residue Lys^32^ (K^32^), glucagon- and insulin analogs each contained ornithine (Orn; O) in place of Lys- and Arg residues to prevent internal trypsin cleavage.

#### (i) Glucagon

An [i, i+4] lactam bridge between the side chains of Lys^13^ (K^13^) and Glu^17^ (E^17^) was inserted to prevent nascent β-sheet formation while favoring a bioactive α-helical structure (44); this lactam-constrained (lc) analog is designated *lc-glucagon*. A C-terminal -EEK extension was added to increase solubility, allow trypsin-mediated ligation to the N terminus of a single-chain insulin analog (SCI; Fig. 3B and Table S1), and decrease potency to better match insulin. The latter emulates preferred molar ratios in prior co-administration studies, which favored reduced relative glucagon activity (31).

**Figure 3.**
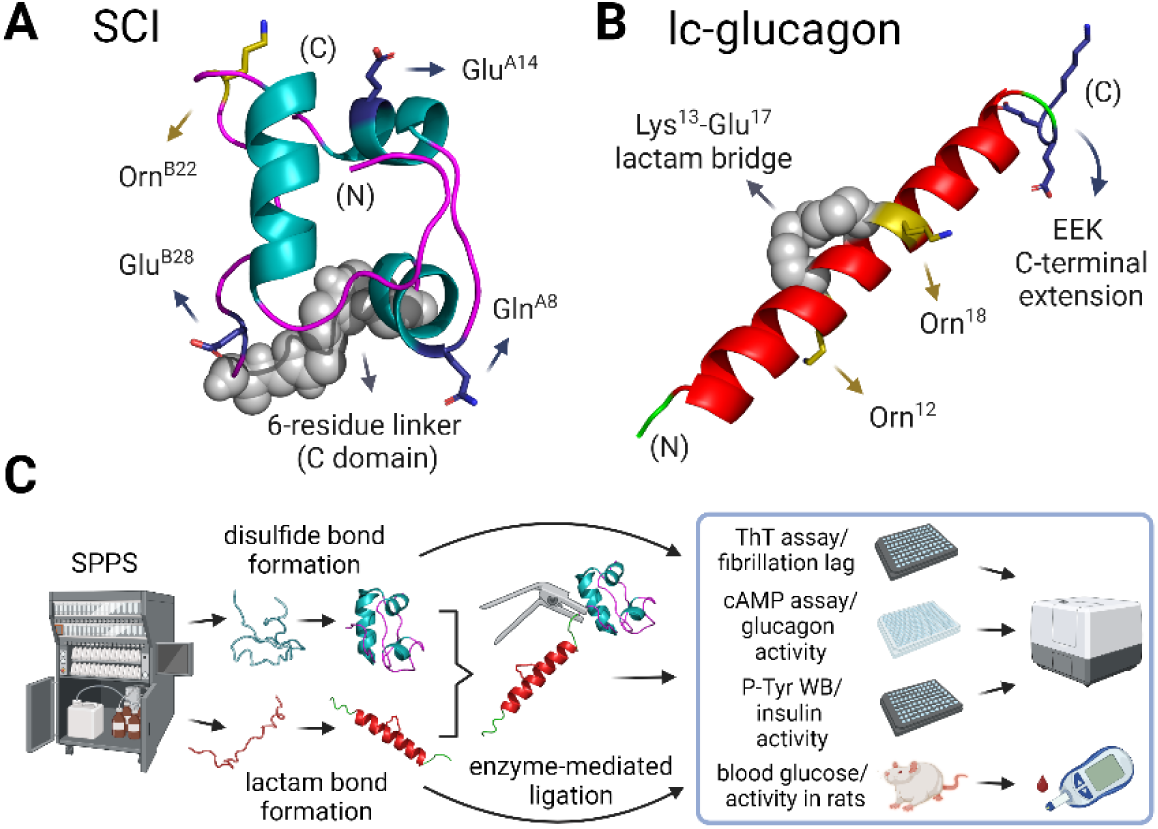
Analog design and workflow. (A) Model of SCI and its modifications: single chain with a 6-residue linker (foreshortened C domain) between B- and A segments. Additional modifications were introduced to enhance solubility (Glu^A14^, Glu^B28^), stability (Glu^A14^), and activity (Gln^A8^); substitution by Arg^B22^ by Orn removed a tryptic site to permit trypsin-mediated ligation. SCI structure is shown based on NMR structure of homologous analog (PDB ID: 5WBT). (B) Model of lc-glucagon and its modifications. The analog was designed to contain (a) a lactam bond between the side chains of K^13^ and E^17^, (b) an -EEK C-terminal extension to increase solubility and (c) Orn^12^, to prevent trypsin cleavage. Structure is illustrated based on crystal structure of a glucagon analog bound to glucagon receptor (PDB ID: 5YQZ). (C) Workflow from SPPS to rat assays. SCI has a folding step (via air oxidation); lc-glucagon exploits on-resin formation of the lactam bond following differential side-chain deprotection. Purified analogs were used for trypsin-mediated ligation. *Box at right,* Isolated hormones and FPs were studied to determine fibrillation lag times, signaling in mammalian cell culture and glucodynamic activities in normal and STZ-diabetic rats. Images were created with Biorender.com.

#### (ii) Insulin

The SCI moiety in the FPs contains design elements conferring thermal stability and fibrillation resistance (46, 47). A central feature is a 6-residue linker between B- and A segments: this tether permits receptor binding but prevents fibril formation (48, 49). Additional modifications enhanced solubility (Glu^A14^, Glu^B28^), activity (Gln^A8^ (50)), and thermodynamic stability (Glu^A14^ (47)) (Fig. 3A and Table S1).

The overall workflow employed solid-phase peptide synthesis (SPPS) followed by oxidative folding (SCI) or lactam-bond formation (lc-glucagon), with respective purification effected by reversed-phase high-performance liquid chromatography (rp-HPLC). The analogs were then fused through trypsin-mediated ligation, connecting the C-terminal Lys of lc-glucagon (E^32^) to the N-terminal Phe in the SCI (Phe^B1^). The resulting FPs were evaluated via fibrillation-, cell-based, and rat assays (Fig. 3C). In addition, structural studies were undertaken using circular dichroism (CD) and solution NMR spectroscopy (Fig. S3-9).

### Fibrillation-resistant glucagon analog

Protection from fibrillation was evaluated in glucagon analogs constrained by a lactam bridge spaced [i, i+4] in the peptide at positions 13-17 or 17-21. Lys-Glu constraints at these positions were previously observed not to impair bio-activity, presumably due to preservation of an active α-helical conformation (44, 45); effects on fibrillation were not reported. The K^17^-E^21^ constraint delayed fibrillation by fivefold relative to lactam-free control glucagon-EEK (lag time 159(±15) hours (hr)). (Fig. 4A). Whereas pairwise K^13^-E^17^ substitutions without a lactam bond exhibited fivefold protection (lag time 160(±17) hr; *ca*. 1 week), insertion of the 13-17 constraint conferred protection for at least 2 weeks (the length of the experiment; Fig. 4A). Based on these findings, the K^13^-E^17^- lactam-constrained glucagon analog (lc-glucagon) was selected for further evaluation. In cell culture assays the analog exhibited full agonism with desired fivefold reduction in potency (EC_50_ of 813 nM [95% confidence interval (CI_95%_) 671-988 nM] *vs*. 163 nM [CI_95%_ 134-198 nM] for wildtype (WT) glucagon; Fig. 4B; Table 1). A similar reduction in potency was observed in studies of glucagon-EEK (EC_50_ of 389 nM [CI_95%_ 342-442 nM]), suggesting that the C-terminal -EEK extension modestly impairs activity (Fig. 4B; Table 1). Fortuitously, this degree of impairment would predict a favorable balance of respective glucagon- and insulin signaling activities in a FP, as extrapolated from the prior co-infusion studies (30–32) and assuming functional independence of the constituent domains (below). To provide a broader context for assessment of the selected lc-glucagon, alternative positions for the lactam-bond formation [residues 9-13, 12-16, 16-20, 20-24, or 24-28] were evaluated without achieving the desired functional properties (Table S6).

**Figure 4.**
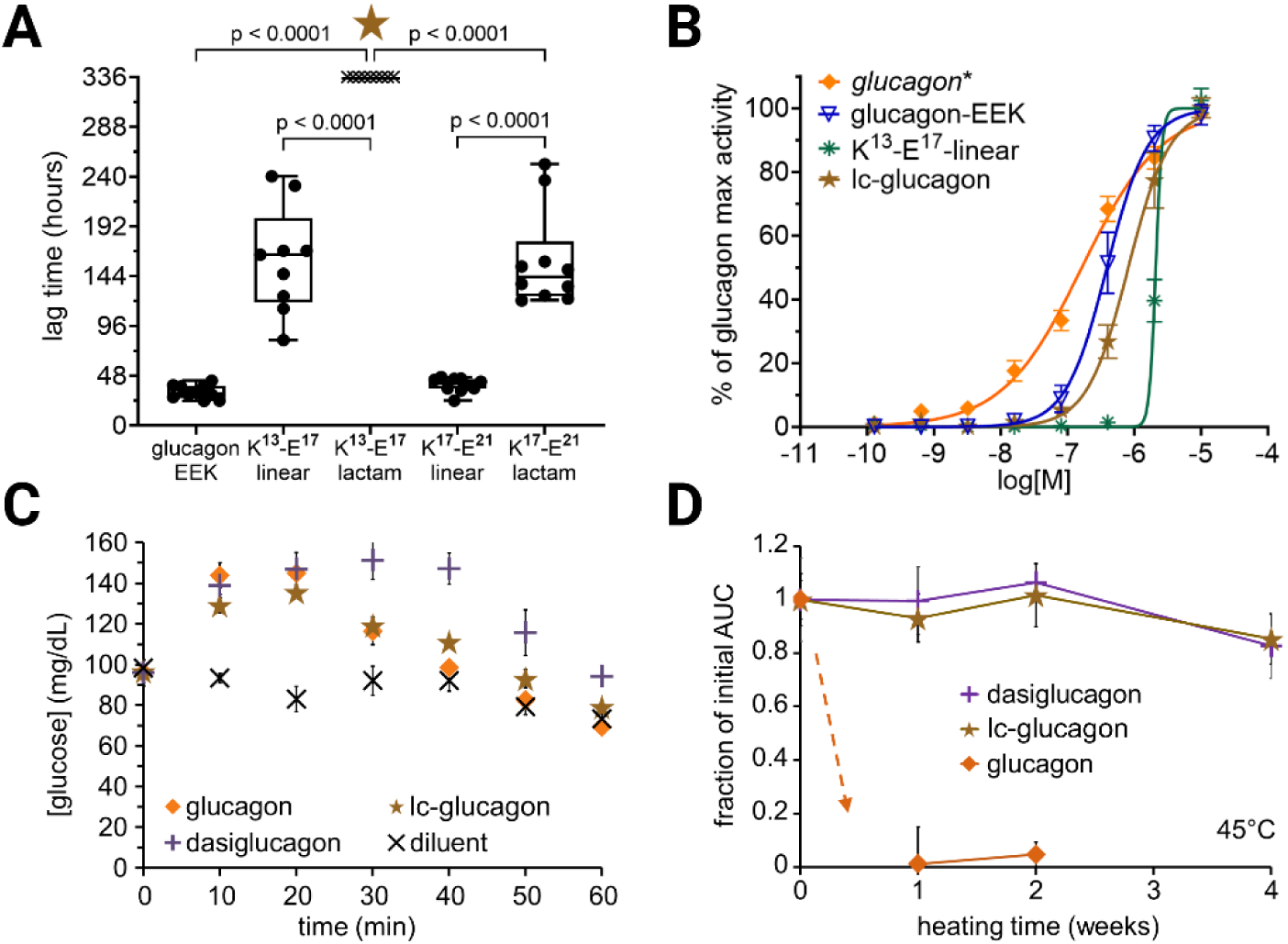
Glucagon analog characterization. (A) ThT assay to measure lag time to onset of fibrillation. Each point represents one well in a 96-well plate set, subjected to continuous agitation at 37 °C. As an unconstrained control, glucagon-EEK carries the same set of modifications except the lactam bond (Table S1): it provided a more workable control than native glucagon, which starts to form fibrils at time t=0 under these conditions. Upper- and lower boundaries of boxes respectively indicate first and third quartiles; upper and lower whiskers demarcate max and min values; and horizontal lines correspond to median. Points at the last recorded time (t = 336 hr) are marked “X” (top) if no fibrils were present. K^13^-E^17^-lactam analog (lc-glucagon) was selected for further studies. (B) Dose-response relationship of [cAMP] production after incubation with glucagon (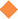) vs glucagon-EEK (∇), K^13^-E^17^-linear (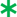) and lc-glucagon (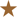); The assay exploited a human cell line engineered to stably overexpress the glucagon receptor (Fig. S12). (C) *In vivo* activities via SQ injection of glucagon (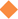) vs lc-glucagon (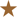), dasiglucagon (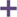) and diluent (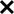) as control; AUC (0-60 min) was calculated and used as the initial time point for a heat stability assay (n=5 per group). (D) Stability assay employing mild agitation at 45 °C. Bio-activities of glucagon (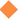), lc-glucagon (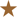) and dasiglucagon (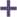) were measured every week using normal rats. Five rats were used for each time point and analog, except for WT glucagon in weeks 1 and 2 when three rats were used. Glucagon and dasiglucagon were injected at a dose of 12 nmol/kg, whereas 23 nmol/kg was used for the less potent lc-glucagon analog. AUC was calculated and graphed as a fraction of the obtained at time 0. Error bars represent SEM.

**Table 1.**
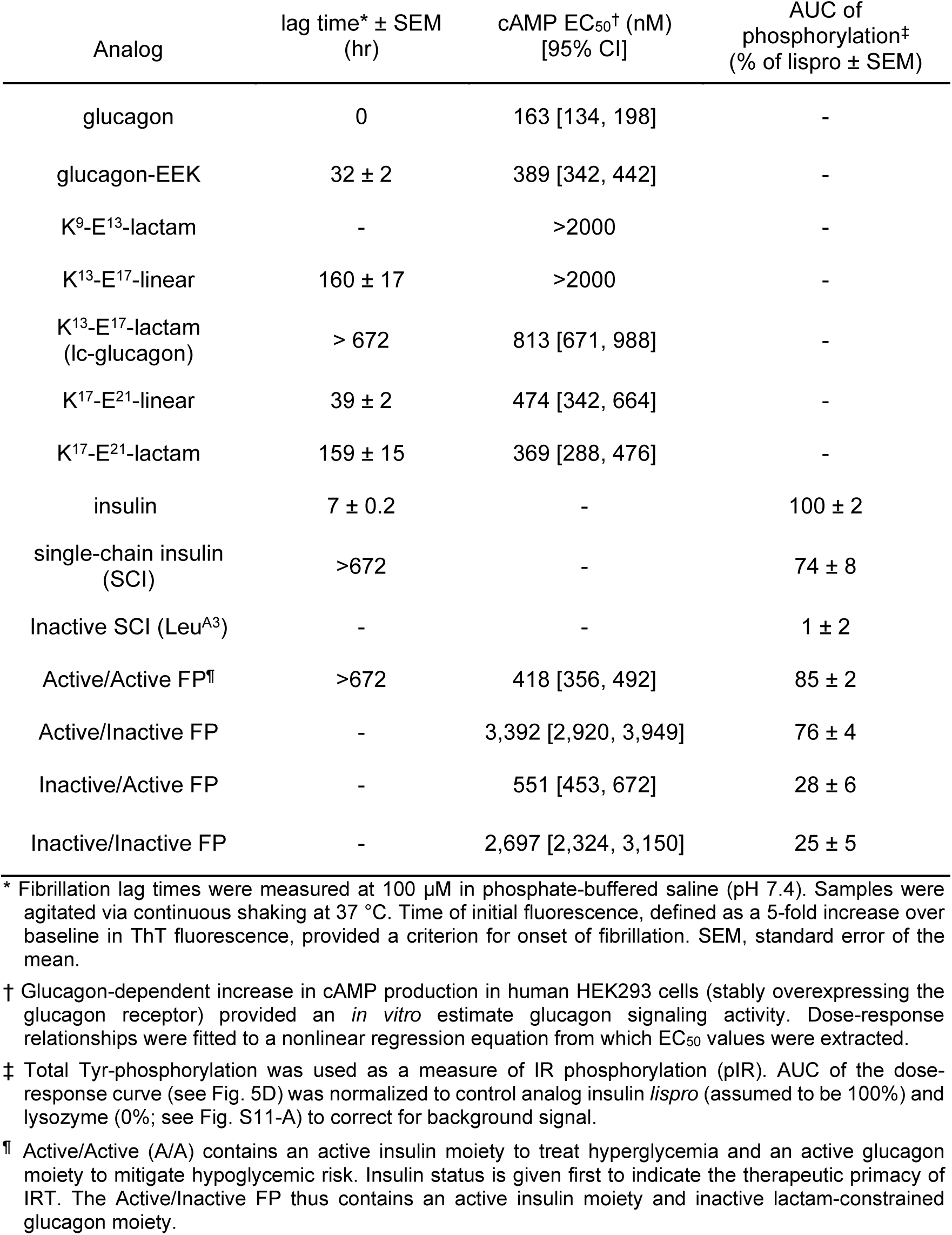
Characterization of Hormone Analogs.

To extend fibrillation studies to a higher temperature, lc-glucagon was kept at 45 °C under mild agitation for 4 weeks, and successive aliquots were tested in rats at weeks 0, 1, 2 and 4 (Fig. 4D). In Figure 4C are shown *in vivo* activities at time t=0 of native glucagon, lc-glucagon and dasiglucagon (a fibrillation-resistant control analog in clinical use as a hypoglycemic rescue agent (51)). Whereas native glucagon was completely inactive within one week, lc-glucagon maintained full activity for 2 weeks and 85(±9.3)% activity at 4 weeks; dasiglucagon exhibited similar robustness.

### Functional studies in vitro

Inactive variants of the SCI and the lc-glucagon were made (Fig. S11) to generate three additional FPs where at least one moiety was inactive. The inactive SCI contained substitution Val^A3^ →Leu (insulin Wakayama), a clinical mutation known to markedly impair receptor binding (52); the glucagon moiety was inactivated by a K^9^-E^13^ lactam bridge (44). These variant FPs provided complementary controls to evaluate respective hormonal activities. For brevity, respective designations are: “insulin analog/glucagon analog” FP; A/A FP (active/active FP), A/I FP (active/inactive FP), I/A FP (inactive/active FP), and I/I FP (inactive/inactive FP) (Fig. 5A). In this nomenclature the first letter (A or I) pertains to the SCI moiety (reflecting the clinical primacy of insulin therapy), and the second letter (A or I) to the lc-glucagon moiety (reflecting its ancillary role).

**Figure 5.**
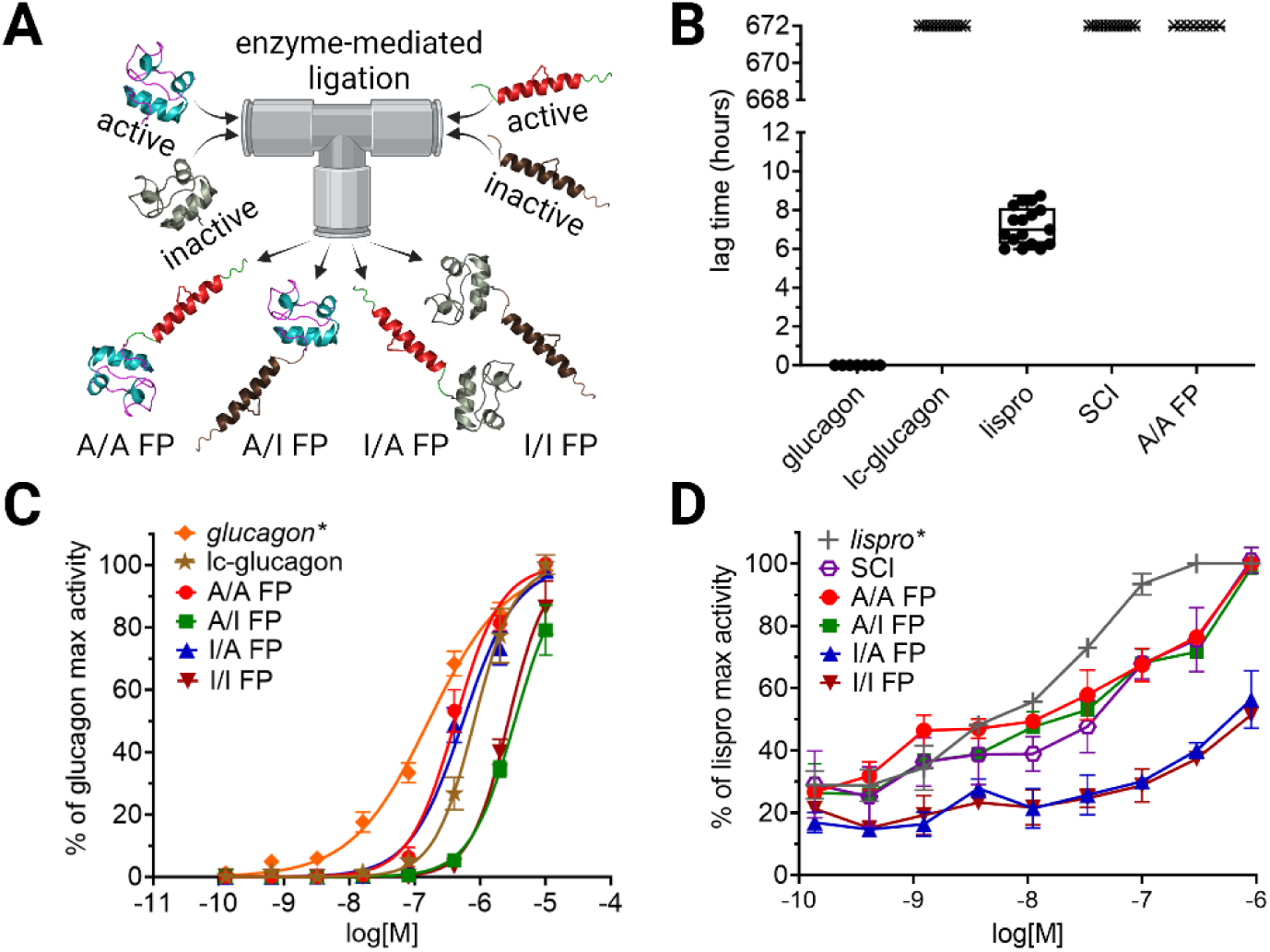
Fusion protein in vitro characterization. (A) “Mixing and matching” inactive versions of SCI and lc-glucagon (Fig. S11) yielded three different control FPs (A/I, I/A and I/I; see main text). (B) Results of ThT fibrillation assays (agitation at 37 °C). Each point represents one well in a 96-well plate; points at the last recorded time (t = 672 hr) are marked “X” (top) if no fibrils were present. (C) *In vitro* glucagon activities of A/A FP (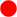) *vs*. lc-glucagon (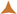), WT glucagon (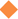) and control FPs (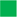 [A/I], (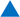 [I/A] and 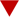 [I/I]; see inset legend); 100% is defined as maximal activity of WT glucagon (*) (D) *In vitro* insulin activity assay where total Tyr phosphorylation was measured through an in-cell western blot, as an indirect way of measuring IR autophosphorylation. Control analogs were provided by insulin *lispro* (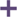) and the isolated SCI (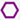); the color code is otherwise as in panel C (inset). In this assay 100% is defined as the maximal activity of insulin *lispro* activity (*). AUC (137 pm to 3 µM) was calculated for all analogs and presented in Table 1. A mammalian cell line overexpressing IR was employed; error bars represent SEM.

In accordance with fibrillation resistance of its component domains, A/A FP exhibited no fibril formation for 4 weeks under continuous agitation at 37 °C (Fig. 5B). In control assays native glucagon formed fibrils within the dead time of the protocol (time t=0) while insulin *lispro* exhibited a lag time of 7.2(±0.2) hr (Fig. 5B). The respective hormonal activities of A/A FP *in vitro*, as evaluated through cell-based assays, were in accordance with the isolated domains. Its EC_50_ for glucagon signaling in cell culture was 418 nM [CI_95%_ 356-492 nM], slightly more active than the K^13^- E^17^-lactam (EC_50_ 813 nM [CI_95%_ 671-988] nM; Fig. 5C and Table 1). Insulin signaling was likewise evaluated in cell culture by measurement of hormone dose-dependent receptor autophosphorylation (pIR/IR ratio as assessed by Western blotting; see Methods). The areas under the curve (AUC, from 137 pM to 3 µM) for A/A FP and the isolated SCI were statistically indistinguishable relative to insulin *lispro* (85(±8)% and 74(±8)%, respectively; Fig. 5D and Table 1). The FPs carrying domain-selective activities (A/I- and I/A FPs) likewise exhibited potencies similar to those their respective parent analogs (Fig. 5C and D). Together, these findings validated the assumption of functional independence implicit in our FP strategy.

### Glucodynamic activities of A/A FP and control analogs

The constituent SCI and K^13^-E^17^-lactam each exhibited full agonism as tested on SQ injection either in rats rendered diabetic by streptozotocin (SCI; Fig. 6A) or in normal rats (glucagon analogs; Fig. 6B) in accordance with the above findings *in vitro*. Comparative *in vivo* assessment of the four FPs (A/A, A/I, I/A and I/I FPs) was then undertaken.

**Figure 6.**
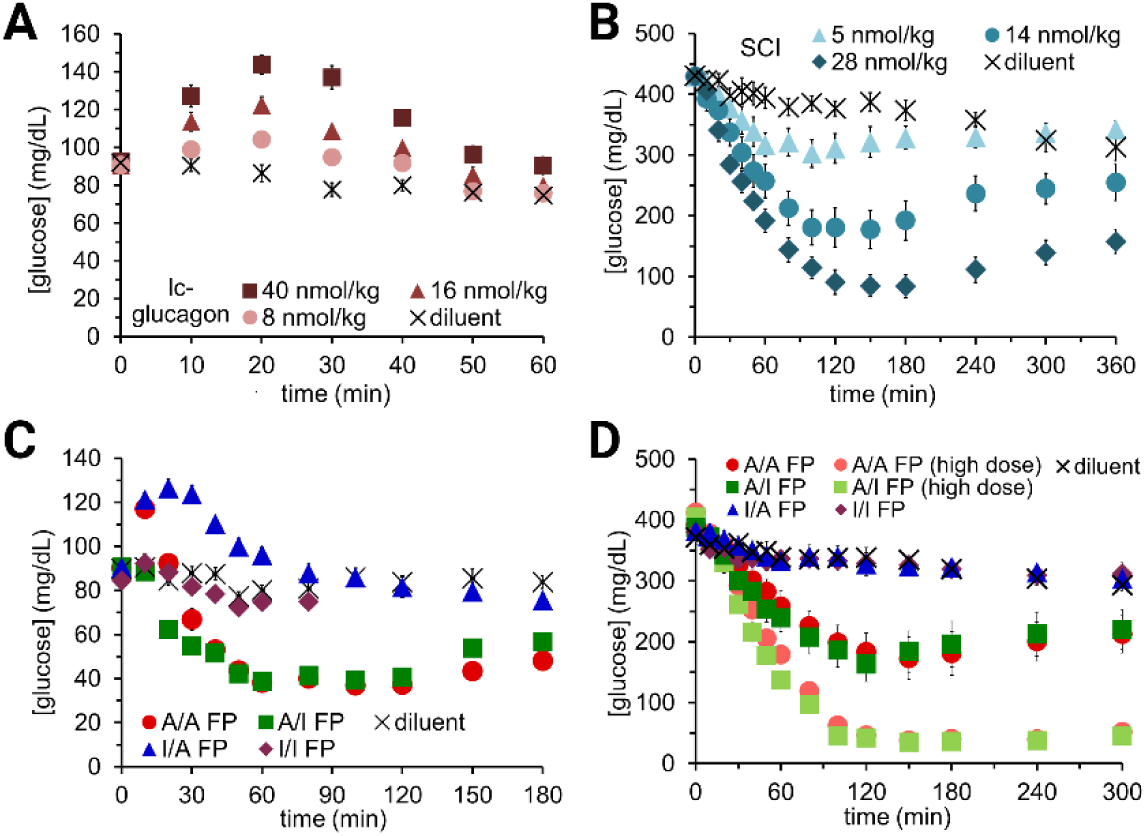
In vivo activities of isolated hormone analogs and FPs. (A) Dose-response relationship of isolated lc-glucagon as measured in non-diabetic rats following SQ injection (n=8 per group): 40 nmol/kg (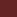), 16 nmol/kg (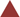), 8 nmol/kg (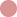) and diluent control (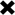). (B) Dose-response relationship of isolated SCI as measured in STZ-diabetic rats (STZ) following SQ injection (n=9 per group): 5 nmol/kg (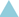), 14 nmol/kg (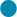), 28 nmol/kg (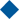) and diluent control (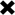). (C) Activities of FPs in non-diabetic rats. The dose in each case was 36 nmol/kg dose with n=9-10 rats per group. Initial increase in BGC was observed only on injection of A/A FP (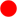; 0-20 min) or I/A FP (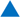; 0-45 min), demonstrating dependence on an active glucagon moiety. Effects of I/I FP (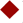) was indistinguishable from that of diluent alone (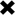). (D) Glucoregulatory activities of the FPs in STZ-diabetic rats (n=10 rats per group). The dose was 24 nmol/kg dose (color code as in panel C) or 36 nmol/kg (A/A FP [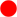] and A/I FP [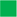]). Activities were measured under hyperglycemic conditions (mean initial BGC *ca*. 400 mg/dL); error bars represent SEM.

#### (i) Non-diabetic rats

In normal rats A/A FP exhibited both insulin- and glucagon activities (Fig. 6C). On SQ injection of A/A- and I/A FPs (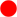 and 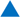 in Fig. 6C), but not A/I or I/I FPs (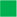 and 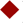), BGCs initially rose (time t=0-15 min) as expected, whereas A/A and A/I FPs (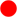 and 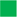), but not I/A or I/I FPs (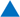 and 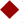) directed a subsequent reduction in BGC (time 15 min – 3 hr). Effects of the doubly-inactive analog I/I FP (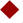) were similar to a those of control buffer injections.

#### (ii) Diabetic rats

STZ rats enabled measurement of insulin activities under hyperglycemic conditions (mean glycemia 400 mg/dL; Fig. 6D). As expected, A/A- and A/I FPs (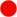 and 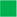 in Fig. 6D) exhibited similar potent insulin activities. This result is in accordance with co-injection of the independent analogs at a 1:1 molar ratio (Fig. S2), reflecting the more rapid absorption and briefer duration of glucagon signaling relative to insulin, as well as the very weak action of glucagon under hyperglycemic conditions (33, 34).

The contrasting PK/PD properties of insulin and glucagon on SQ injection of a rapid-acting formulation confounded observation of glucagon action subsequent to t=15 min post-injection and motivated the following use of continuous IV infusion to mimic a long-lived (basal) depot.

### Glucagon moiety activates glucose production under hyperinsulinemic conditions

A basal depot (whether SQ or as bound to a carrier protein in the circulation) would in principle provide continuous co-availability of the two hormonal activities. To emulate such behavior, we undertook a steady rate of A/A FP IV infusion. Hyperinsulinemic clamps were established in rats at two set points, euglycemia (BGC 110 mg/dL) and hypoglycemia (60 mg/dL). The former was after 30 minutes (min; Fig. 7B), and the latter at t=0 (Fig. 7A). Glucose infusion rates (GIR), a measure of exogenous glucose needed to maintain the clamped BGC, were recorded. The studies focused on A/A- and A/I FPs to test whether an active glucagon moiety might confer protection from hypoglycemia during the course of basal IRT.

**Figure 7.**
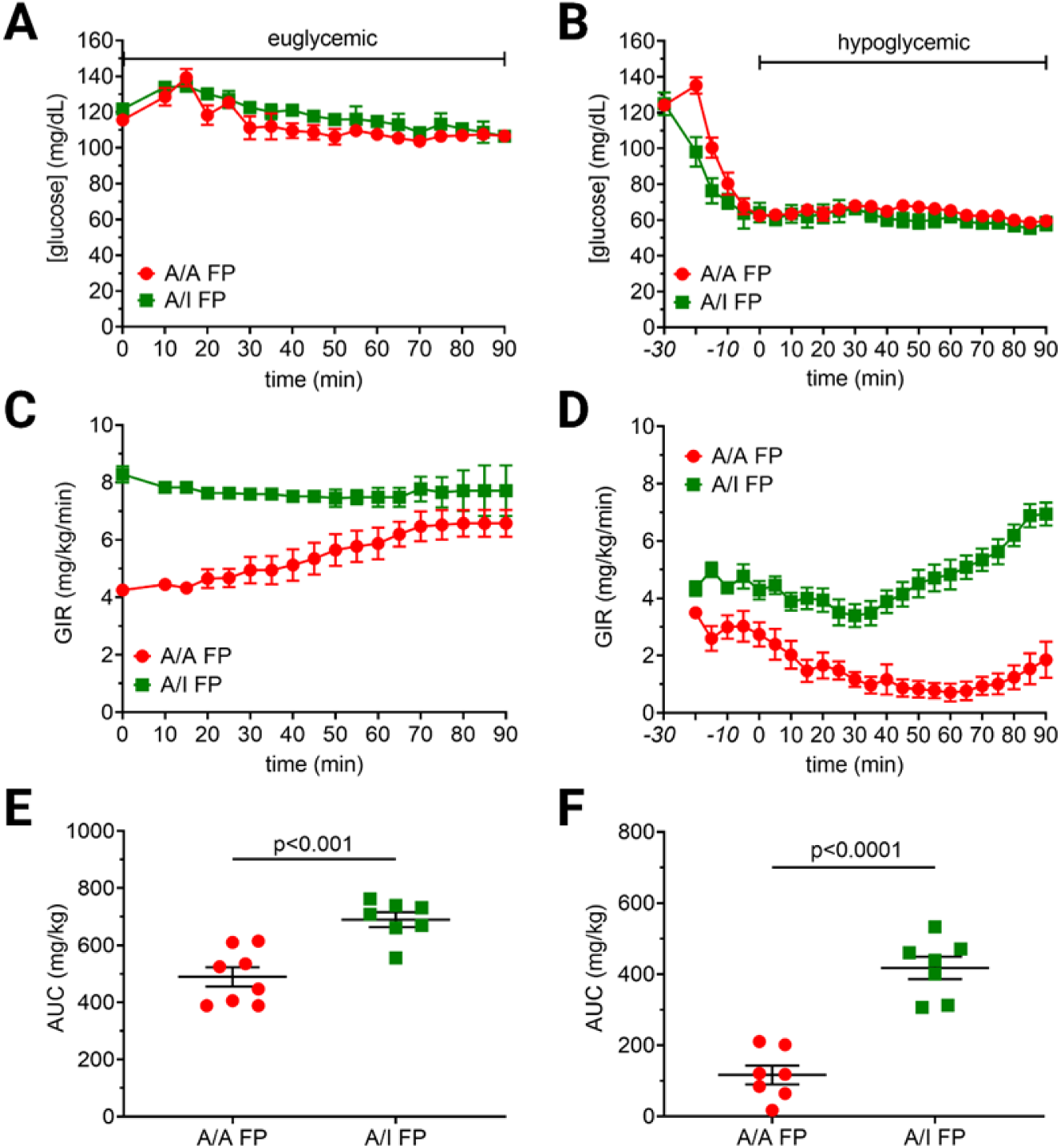
Hyperinsulinemic clamp studies of A/A and A/I FPs in rats. (A) Hyperinsulinemic-euglycemic clamp at 110 mg/dL. (B) Hyperinsulinemic-hypoglycemic clamp at 60 mg/dL. Establishing the clamp required 30 min to stabilize the hypoglycemia. (C, D) GIR of clamps at 110 and 60 mg/dL, respectively, indicating how much glucose is needed to be infused to maintain the clamp. The infusion rate was modified and recorded every 5 min. (E, F) Areas under the curve of the GIR data for the euglycemic and hypoglycemic clamp, respectively. Student’s t-test was used to determine statistical differences; error bars represent SEM. Seven rats were used per group, except for A/A FP in the euglycemic clamp (n=8). Color code: A/A FP (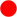) *and A/I FP* (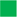).

Under both glycemic set points A/A FP (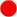 in Fig. 7C, D) exhibited a reduced GIR relative to A/I FP (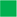). Strikingly, this effect was more pronounced under hypoglycemic conditions (Fig. 7C, D). The resulting GIR plots displayed a significant difference in the proteins’ ability to stimulate endogenous glucose production. To get a numerical approximation of the total glucose delivered to the animals during the glucose clamp, glucose infusion rates were recorded at regular time intervals. The trapezoidal rule was used to estimate the area under the curves (AUCs) of the glucose infusion rates over the course of the glucose clamps (0 to 90 min): 489 vs 690 mg/kg, p<0.005 under euglycemic clamp conditions; 117 *vs*. 418 mg/kg, p<0.0001 under hypoglycemic clamp conditions, demonstrating that A/A FP markedly effected glucagon signaling despite the hyperinsulinemic conditions (Fig. 7E, F). Because rates of glucose disposal in muscle are not thought to be regulated by glucagon, we ascribe the GIR findings to endogenous glucose production. Further, excursions in BGCs associated with A/A FP infusion were independent of baseline hypoglycemic counter-regulation, as evidenced by the similar epinephrine concentrations induced by A/A- and A/I FPs (Fig. S10-E, F).

### Biophysical studies of the fusion protein

Comparative far-ultraviolet CD spectra support the structural independence of the glucagon- and insulin-derived moieties as the calculated sum of the spectra of the isolated lc-glucagon and SCI closely correspond to the observed spectrum of A/A FP spectra as normalized by molar ellipticity [θ] per molecule. This correspondence was maintained at 4, 25, 37 and 45 °C (Fig. 8A, B, D and E, respectively) and recapitulated when [θ]_222_ was measured as a function of temperature in the range 4-88 °C (Fig. 8G). Artificial-intelligence (AI)-based structural modeling (53–55) predicted such independent domain folding (Fig. 8H; see Discussion).

**Figure 8.**
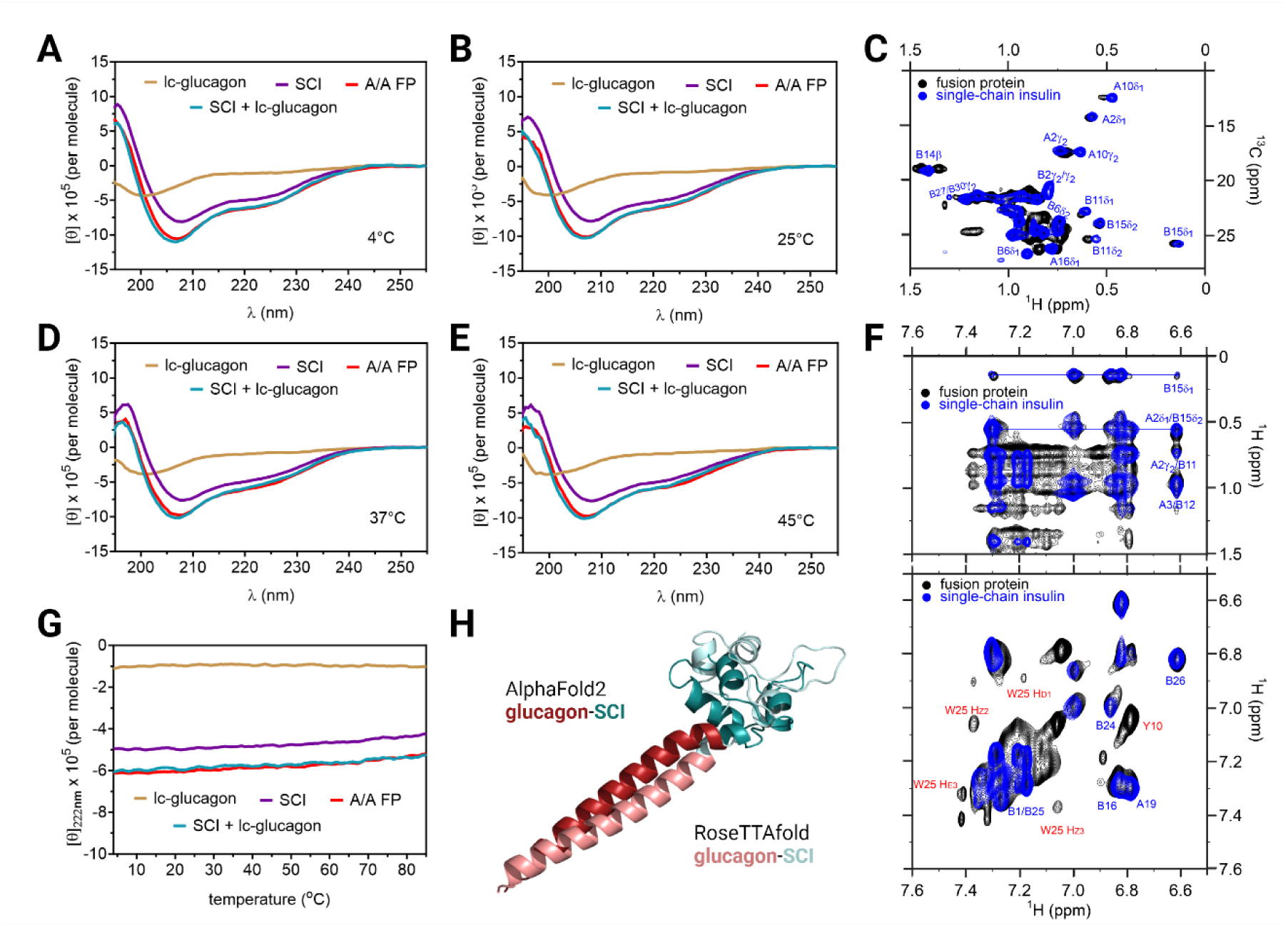
Structural independence of individual moieties in the FP. (A, B, D and E) CD spectra (190-255 nm) at 4, 25, 37 and 45 °C show that the addition of the lc-glucagon spectra to the SCI spectra matches the spectra of A/A FP at all temperatures (color code inset; 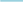 indicates calculated sum of observed spectra). Random-coil pattern of lc-glucagon spectrum (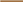) is in accordance with CD studies of WT glucagon (102). The α-helical signature of the isolated SCI (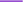) is in accordance with structures of homologous domains (47, 49). (G) CD-derived molar ellipticity [θ], measured at helix-sensitive wavelength 222 nm from 4-88 °C, exhibits alignment of observed A/A FP signal (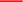) with the sum of the ellipticities its isolated components (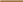 [lc-glucagon] and 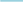 [SCI]). (H) AI-based structure prediction of lactam-free FP by AlphaFold2 and RoseTTAfold (53–55) (color code inset). (C) Overlay of methyl ^1^H-^13^C NMR HSQC spectra of A/A FP (black) and isolated SCI domain (blue). (F) Long-range signature NOEs between methyl protons and aromatic protons (top panel) and aromatic chemical shifts in ^1^H-^1^H TOCSY spectra (bottom panel) are notable for corresponding NMR features: A/A FP (black) and isolated SCI domain (blue). These data provide evidence that the insulin motif in the FP maintains native-like structure. Spectra were acquired at a ^1^H frequency of 700 MHz in D_2_O at pD 7.4 (direct meter reading) and at 25 °C.

To further study the structural independence of the individual moieties in the A/A FP, we performed high-resolution NMR spectroscopy (Fig. S6-9 and Tables S1-3). One-dimensional spectra are provided in Figure S6. Although overall ^1^H linewidths are sharp, as expected for a small protein, a subset of broad amide resonances in the SCI spectrum (as in prior NMR studies of insulin and SCIs (46)). The latter are further broadened in the FP (expanded Fig. S7-A, B, C), suggesting a dynamic influence of the tethered lc-glucagon segment; these resonances were nonetheless detectable in a 2D NOESY spectrum (Fig. S7-D). The ^1^H-^15^N main-chain “fingerprint” region of the heteronuclear single-quantum coherence (HSQC) spectrum of A/A FP is shown in Figure S8-A, B in relation to the spectra of the isolated hormone moieties. Acquired at natural abundance, these spectra are consistent with preservation of independent domain folding. This conclusion was corroborated with greater residue-specific resolution by comparison between ^1^H-^13^C HSQC spectra of A/A FP and the isolated SCI analog (Fig. S8-D, E). In addition, corresponding ^1^H-^1^H NOESY spectra exhibited overlapping long-range “signature” NOEs between methyl protons and aromatic protons in association with characteristic patterns of secondary chemical shift (47, 49) (Fig. 8C). Examples of such signature NOEs are provided by core contacts between the side chains of Val^B12^- Tyr^B26^, Leu^B15^-Phe^B24^, and Ile^A2^-Tyr^A19^ (Fig. S9-B). Further, similar patterns of aromatic ^1^H-^1^H TOCSY cross peaks, assigned to respective SCI core residues in the FP and isolated domain, were also observed (47, 49) (Fig. 8F and Fig. S9-F). Dispersion of these and related ^13^C aromatic chemical shifts, well-characterized in an isolated SCI (47, 49) and preserved in the FP (Fig. S8-D, S8-F, and S9-C), provided additional evidence for a correspondence of domain structure. An analogous correspondence of NMR features was observed on comparing spectra of A/A FP with matched spectra of the isolated lc-glucagon (Fig. S8-A, C, E and Fig. S9-A).

Although the isolated lc-glucagon peptide (as a monomer in solution) does not exhibit a stable α-helical conformation, comparative CD studies of the constrained analog and its unconstrained parent peptide were undertaken in the presence of a helicogenic co-solvent (trifluoro-ethanol [TFE]). Spectra were obtained as a function of the percentage of TFE. At low concentrations, TFE was more effective in inducing helical CD features in lc-glucagon than in its unconstrained parent peptide, suggesting that closure of the K^13^-E^17^ lactam bridge biases the otherwise-disordered nascent peptide conformation toward α-helix (Fig. S3).

## Discussion

Insulin therapies responsive to blood glucose concentration have been pursued for decades to enhance the safety and efficacy of IRT, especially in T1D (14). A leading approach in current clinical use exploits CGM feedback to enable algorithm-guided insulin-pump delivery (16, 17) (Fig. 1A). Other technologies envision glucose-dependent release of insulin from the SQ depot (18–20) (Fig 1B) or glucose-responsive hormone binding to modulate insulin clearance or bioavailability (21–23) (Fig. 1C). A key frontier of pharmacology envisions the engineering of glucose-regulated “switches” that modulate the release or activity of insulin (24) (Fig. 1D). Because CGM-mediated feedback in current closed-loop systems is limited by a time lag (*ca*. 15-20 min) between changes in interstitial glucose concentration relative to changes in BGC (56), interest has recently focused on unimolecular GRIs containing a chemical glucose sensor for real-time modulation of insulin activity in the blood stream (24). Such embodiments envision a coupling between insulin conformation and binding of glucose to the chemical sensor. To our knowledge, the safety or efficacy of such “smart” analogs has not been evaluated in clinical trials.

The present bihormonal approach dispenses with the molecular requirement for a chemical glucose sensor by exploiting an endogenous glycemia-associated switch in hepatic hormone responsiveness (25, 28, 29). Because the duration of native glucagon signaling is shorter than that of insulin, a glucagon-insulin FP would be more compatible with a basal (long-acting) formulation than with a prandial (short-acting) formulation. The former embodiment was herein modeled via continuous IV infusion. Although appropriate for basal-bolus therapy of patients with T1D (with the prandial component provided by a current rapid-acting formulation), it is likely that this biotechnology would also find translational application in basal-only treatment of patients with Type 2 DM (T2D), especially in a once-a-week form (for review, see (57)). The molecular simplicity of a bihormonal FP and its ultra-stability also promise real-world cost advantages (in manufacture, delivery and storage) relative to more complex glucose-responsive technologies (Fig. 1). The FP’s marked resistance to fibrillation above room temperature may also enhance global access.

### Multiple Agonism

Motivated by preclinical successes of glucagon/GLP-1 dual-agonists in reducing weight in obese rat models (39), multi-agonist peptides are emerging as promising therapeutic approaches to the management of human diabetes and obesity (58, 59). The similar peptide sequences of GLP-1, glucagon, and to a lesser extent, glucose-dependent insulinotropic polypeptide (GIP)—and molecular homologies among their respective receptors—have enabled design of dual- and triple-agonists as exemplified by tirzepatide (Mounjaro^®^), a dual GLP-1/GIP agonist recently approved by the U.S. Food & Drug Administration for the treatment of T2D (60) and by the current development of Retatrutide, a GLP-1, GIP, and glucagon receptor agonist (61). Although not appropriate for management of T1D, these and related peptides are also under regulatory review for the treatment of obesity in the absence of diabetes (40, 62–64). Hypoglycemic risk is minimal except when used in combination with insulin or other insulin secretagogues (such as sulfonylureas).

The present bihormonal FP is different in concept from incretin multi-agonists because insulin and glucagon are unrelated in sequence and bind to unrelated classes of membrane receptors. Our approach exploits differential hormonal response in the liver as a function of glycemia (25–29), and its molecular design considered a salient feature of respective hormone-receptor complexes: the free C-terminal residues of glucagon and free N-terminal B-chain residues of insulin (41–43). Fortuitously, glucagon tolerates C-terminal extensions (65, 66). Conversely, whereas N-terminal B-chain residues are important for the foldability of proinsulin, they are dispensable for receptor binding; deletions or modifications are allowed among active analogs (42, 67, 68). Indeed, Huang et al. recently described an insulin/glucagon/GLP-1 tri-agonist that uses “click” chemistry to bind the B-chain N-terminus of insulin to a C-terminal residue of a dual agonist (63). To our knowledge, the potential glucose responsiveness of this construct and its physical stability have not been evaluated.

### Basal glucose-responsive insulin

Once insulin or glucagon engage their respective receptors, ensuing cellular signaling pathways and associated biological responses exhibit intrinsic differences in duration. Whereas glucagon action attenuates after 20-30 min, insulin action continues for *ca*. three hours despite rapid receptor-mediated clearance (33, 34). On SQ or IV bolus injection of a rapid-acting A/A FP, the initial phase of glucagon action dissipated prior to the delayed phase of insulin action (Fig. 6). This mismatch would undermine protection from late hypoglycemia (2-4 hr post-injection). Clinical translation thus envisions an initial embodiment of the FP as a long-acting agent. The continuous presence of the FP in a long-term SQ depot or in the bloodstream (*e.g*., as bound to albumin) would in principle enable prolonged protection from hypoglycemia. Current basal insulin analogs use either of two strategies to achieve protracted activity (69): (i) an SQ depot of insulin hexamers that slowly release dimers and monomers into circulation (Fig. 9A; exemplified by the isoelectric precipitation of insulin *glargine* (70)); and (ii) a “circulating depot” provided by insulin-albumin complexes, and slowly released to target cells (Fig. 9B; exemplified by acylated insulin analogs [*e.g*., *detemir*, *degludec* or *icodec*] (71, 72)). Long-lived SCI-immunoglobulin fusion proteins have also been described (for review, see (69)). These molecular tactics may in principle be generalized to a glucagon-insulin FP. Additionally, continuous SQ infusion (pump therapy) would be an alternative way to ensure glucose-responsive basal IRT.

**Figure 9.**
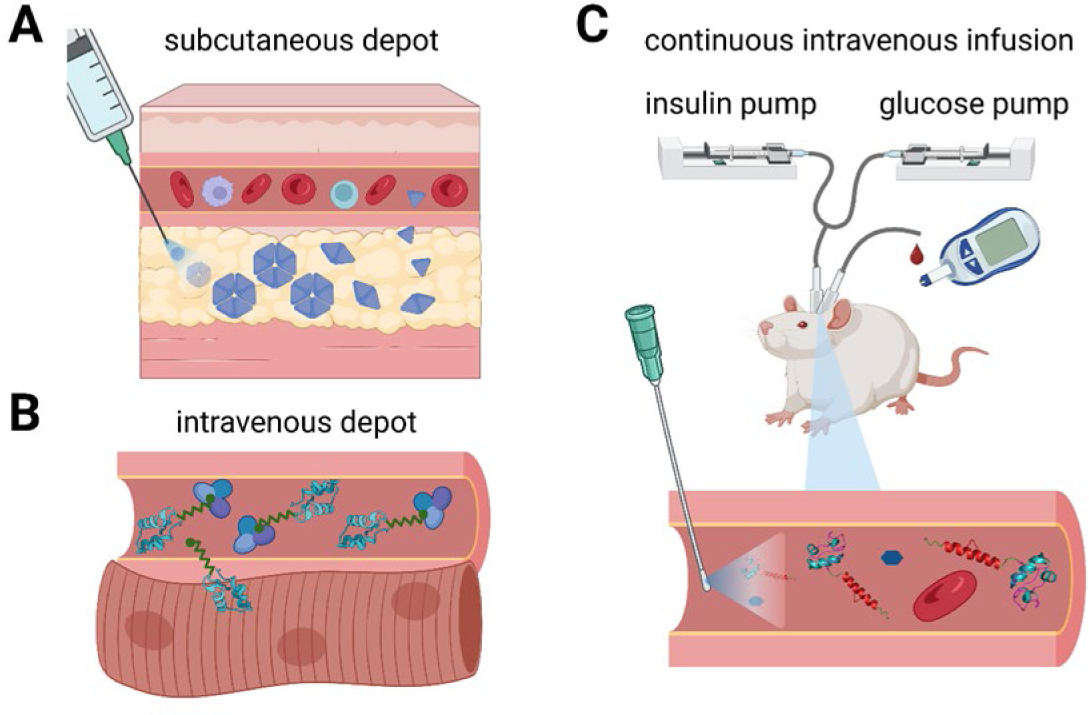
Continuous IV infusion mimics basal insulin action. (A) Classical basal insulin analogs, as exemplified by pI-shifted insulin *glargine* (70), provide a long-lived SQ depot due to self-assembly of zinc insulin hexamers (69). Slow dissociation into dimers and monomers allows absorption into the bloodstream. (B) Next-generation basal insulin analogs, such as once-a-week insulin icodec (71), exploit strong but reversible binding to albumin as a form of an “IV depot”: slow and reversible release precedes binding to target cells. (C) Hyperinsulinemic clamps in rats enable study of the A/A FP with its continuous IV infusion, emulating slow, incremental basal insulin analogs from a SQ or circulating depot. Here, this approach overcame briefer duration of glucagon activity relative to insulin action following SQ injection of a rapid-acting depot. Glucose was concurrently administered IV at a variable rate to maintain a pre-established glycemic set point; GIRs and BGCs were recorded. Extension of the FP strategy to a rapid-acting formulation would require novel re-engineering of the glucagon moiety to achieve a signaling duration (once its receptor was engaged) commensurate to that of insulin action. Images were created with Biorender.com.

To emulate a long-acting bihormonal FP *in vivo*, we employed continuous IV infusion in rats in conjunction with hyperinsulinemic clamps; glucose was co-infused at variable rates to maintain glycemia at fixed target levels (Fig. 9C). The markedly different GIR between A/A and A/I FPs under hypoglycemic conditions (Fig. 7D) provided evidence that the glucagon moiety can stimulate endogenous glucose production despite pharmacologic hyperinsulinemia. This is in accordance with prior clamp studies in dogs in which the two hormones were independently administered: the findings indicated increased sensitivity of the liver to glucagon during insulin-induced hypoglycemia (25, 28, 29). Conversely, glucagon signaling in the liver is down-regulated under hyperglycemic conditions. Such intrinsic hepatic physiology provides an endogenous switch in the absence of a chemical glucose sensor in the therapeutic entity.

A subtle difference in the physiology of glucagon signaling was observed on comparison of the A/A- and A/I FPs in respective hyperinsulinemic clamps under euglycemic or hypoglycemic conditions (Fig. 7C *vs*. Fig. 7D). Whereas time-dependent convergence of the A/A and A/I GIR curves under euglycemic conditions suggests progressive attenuation of glucagon signaling in the course of the clamp (Fig. 7C), such convergence did not occur under hypoglycemic conditions: the protective effect of the A/A FP persisted throughout the clamp period (Fig. 7D). Physiological attenuation of glucagon signaling in the liver under euglycemic or hyperglycemic conditions is homeostatic and has been proposed to reflect multiple molecular mechanisms, including decreasing intracellular cAMP concentrations, changes in gluconeogenic and glycogenolytic fluxes, hormone-dependent programs of gene expression related to mitochondrial metabolism, and allosteric effects of glucose-6-phosphate accumulation on the enzymatic activities of glycogen synthase, glycogen phosphorylase and glucokinase. The contribution of the latter allosteric mechanisms to the waning of glucagon signaling (despite sustained glucagon concentrations in the blood) was highlighted in a recent dog study (73). From a translational perspective, the maintenance of glucagon signaling by the A/A FP under hypoglycemic conditions and its waning under euglycemic conditions would in principle enhance therapeutic efficacy. We anticipate that a deeper basic understanding of glucagon signaling—in particular physiologic determinants of duration and molecular mechanisms of down-regulation—would enable extension of the present FP strategy from basal to prandial formulations.

### Prevention of protein fibrillation

Glucagon’s intrinsic propensity to form fibrils has previously been addressed through improved formulations (74, 75) or via incorporation of natural and/or non-natural amino-acid modifications (51, 76). Our approach is based on divergence between the hormone’s active α-helical structure and its inactive β-sheet-rich fibrillar structure (41, 77): a side-chain “bridge” with appropriate spacing in the peptide can bias a local α-helical conformation and preclude a regular β-sheet (78). Lactam bonds between the side chains of Lys and Glu, with [i, i+4] spacing, have been utilized to constrain the active conformation of glucagon by Hruby and coworkers (44, 45); our study in part built on these findings and extended the characterization to physical stability. We found that lactam-based protection from fibrillation is position-specific: whereas a K^17^-E^21^ lactam bond did not fully prevent fibrillation (Fig. 4A; lag times ca. 1 week at 37 °C), the K^13^-E^17^ lactam bond in the present FP effectively prevented fibrillation (Fig. 4A; lag times > 2 weeks). Although the underlying molecular mechanisms are not fully understood, we envision that the K^13^-E^17^ lactam bond either delays nucleation or disrupts the mature fibrillar structure; these possibilities are in accordance with prior studies describing the key contribution of native residue Tyr^13^ to fibrillation (77).

The economy of a single lactam bond circumvents the need for multiple amino-acid substitutions as exemplified by dasiglucagon, which carries seven modifications, including one non-natural amino acid (51). Such economy may reduce potential immunogenicity on long-term clinical use. Lactam bonds have been utilized to stabilize and constrain the structures of diverse peptides, for example, DiMarchi and colleagues introduced [i,i+4] lactam bonds to enhance glucagon receptor agonism in GLP-1/glucagon dual-agonists (39). Such tactics have also been applied to glucagon-related analogs of oxyntomodulin, GLP-1, and GLP-2 (79–81). In addition, a similar approach conferred fibrillation resistance to Aβ peptide 1-28 (82, 83). As a seeming paradox, at unfavorable positions in the peptide, lactam bonds can *stimulate* fibril formation, as was also observed in an Aβ analog wherein a specific lactam bridge (D^23^-K^28^) “locks-in” a naturally occurring salt bridge that contributes to nucleation of protofibrils (84).

Insulin also exhibits an intrinsic propensity to form amyloid (85, 86), mitigated in current pharmaceutical formulations by stabilizing additives (such as zinc ions and phenolic ligands) and multimeric assembly (87). The present FP exploits a foreshortened C domain in a single-chain polypeptide (88). This topology is compatible with biological activity but presumably restricts the “splaying” of the A- and B chains needed to form a cross-β assembly (46, 47, 49). Such fibrillation-resistant SCIs have been investigated to circumvent the present cold chain of global insulin distribution (89, 90).

### Challenges to clinical translation

Clinical translation would require a scalable method of manufacture. Although scalable protocols for SPPS have been developed in the preparation of incretin analogs (38–40), it is possible that, as a class, glucagon-insulin FPs may be manufacturable, in part or in whole, via recombinant expression. SCIs themselves are amenable to over-expression in *Pichia pastoris* (46, 47), and so presumably would be a lactam-free FP. Although insertion of a lactam constraint would presently not be possible in a recombinant system, such a bridge might be replaceable by paired-Cys hydrocarbon “stapling” as a post-fermentation step (91). In such an embodiment Orn would not be required. Development of such alternative embodiments would require recharacterization of structure-activity relationships with attention to relative hormonal activities. Further, because the present enzymatic ligation scheme employed a pre-modified glucagon peptide and pre-folded SCI, our results do not address the efficiency of nascent folding of a biosynthetic FP, either *in vivo* or *in vitro* from inclusion bodies. It is nonetheless encouraging that *de novo* folding algorithms (AlphaFold2 and RoseTTAfold; see Methods (53–55)) predict that an unconstrained glucagon analog fused to an SCI moiety would exhibit independent foldability into their respective active structures (Fig. 8H).

Clinical application of a glucagon-insulin FP as a basal formulation would circumvent a PD mismatch between the duration of glucagon signaling relative to insulin signaling (92, 93). To explore further the utility of such chimeric polypeptides, it would be necessary to examine their efficacy over a range of glucagon:insulin activity ratios in larger mammals, *i.e*., models with higher translational value to the development of human therapeutics. Metabolic benefits of glucagon, especially in combination with incretins (*e.g*., dual agonists (39)), have been investigated at high doses of glucagon, and so it is not clear whether synergistic actions of glucagon and insulin would manifest at lower glucagon concentrations (for review, see (94, 95). In addition, both insulin and glucagon alter protein metabolism (96), and so it would be important to assess the effect of glucagon-insulin FPs on that readout. Effects of such FPs on fat metabolism would likewise require evaluation. Despite such complexity of potential endocrine actions (whether favorable or adverse), this work has demonstrated that combining insulin and glucagon within a single polypeptide is feasible and can control glucose metabolism in a predictable way.

### Concluding remarks

There are currently two major barriers limiting the efficacy and safety of IRT in DM management. First, insulin-induced hypoglycemia is associated with acute and chronic sequalae (7), including cognitive decline (8); this threat often discourages aggressive glycemic targets in regimens otherwise intended to prevent or delay microvascular complications (2). Second, the physical instability of insulin (*i.e*., susceptibility to fibrillation above room temperature (85)), making necessary strict regulatory guidelines regarding storage and shelf life (88), in turn impose a complex and costly global cold chain of distribution (89). The latter considerations are even more severe in the case of glucagon (36). Fibrillation of these hormones leads in each case to pro-inflammatory, inactive aggregates.

In the present study we describe the design and characterization of an ultra-stable glucagon-insulin FP in an effort to circumvent both these barriers. FP design reflects the three-dimensional structures of the respective hormone-receptor complexes (41–43, 48) and incorporates elements inhibiting fibrillation of either domain. Glucose-responsive protection from hypoglycemia, as demonstrated in rats, exploits a physiological hepatic switch in respective hormonal responsiveness in the absence of a molecular glucose sensor in the therapeutic entity. We envision that the molecular simplicity of this approach and its ultra-stability may enhance global access to IRT, a key real-world challenge given the growing prevalence of both T1D and T2D in many regions of the world (90). In both affluent and developing societies molecular technologies to mitigate hypoglycemic risk promise to promote tight glycemic control, especially in the context of hypoglycemic unawareness (12), in turn enhancing safe and effective diabetes management.

## Materials and Methods

### Peptide synthesis

Glucagon and insulin analogs were chemically synthesized through SPPS as described (97); protocols are provided in the Supplement. The sequence of A/A FP is: HSQGTFTSDYSO<u>K</u>LD-S<u>E</u>OAQDFVQWLMNT*EEK*FVNQHLCGSHLVEALYLVCGEOGFFYTPETEEGPOOGIVEQCCQSI CSLEQLENYCN. Substitutions are shown in red; underlined are residues K^13^ and E^17^, sites of lactam-bond formation; and linking tripeptide segment (E^30^-E^31^-K^32^) is italicized. In Table S1 are given a complete set of sequences and amino-acid substitutions.

### Preparation of insulin analogs

The crude peptides in reduced form were folded by air oxidation to induce disulfide pairing (97). Insulin *lispro* was prepared as a synthetic control through a semi-synthesis of *des*-octapeptide insulin derived from a “DesDi” precursor and an octapeptide (sequence GFFTKPT) (98). Insulin analogs were purified by preparative rp-HPLC.

### Preparation of lactam glucagon analogs

On-resin selective deprotection of Lys(Alloc) and Glu(Allyl) residues with subsequent lactam bond formation on the desired positions followed a protocol previously described by Ahn et al. (2001) (44). Analogs were purified by preparative rp-HPLC.

### Trypsin-mediated ligation

A peptide bond was introduced between the C-terminal Lys of glucagon analogs and the α-amino-group of Phe^B1^ of the SCI through trypsin-mediated ligation in an organic co-solvent (99). Excepting the C-terminal Lys of glucagon analogs, all other basic residues in the FP were replaced with Orn to avoid internal tryptic cleavage sites. Following enzymatic ligation, FPs were purified by preparative rp-HPLC.

### Fibrillation Assays

Susceptibility to fibrillation was assessed through measurement of the lag time prior to the exponential phase of fibril formation via a thioflavin T (ThT) fluorescence assay (99, 100). The measurements were performed at 37° C with agitation in a 96-well plate (Corning, Corning, NY) read on a Synergy H1 automated microplate reader (BioTek, Winooski, VT).

### CD Spectroscopy

Far-UV CD spectra were acquired in 10 mM potassium phosphate (pH 7.4) and 50 mM KCl using a Jasco J-1500 spectropolarimeter (Jasco, Tokyo, Japan); peptide- or protein concentrations were 25 mM. Normalized ellipticities [θ] was independently calculated *per molecule* (to allow addition of spectra) and *per residue* (Fig. S5).

### Structure prediction

Initial modeling of the FP exploited deep learning-based methods as implemented in AlphaFold2 (53), as implemented in ColabFold (54) and RoseTTAfold (55) (https://robetta.bakerlab.org/). The protein sequence did not contain a lactam bond or Orn substitutions.

### NMR spectroscopy

NMR data were recorded using a BRUKER 700 MHz spectrometer equipped with ^1^H, ^19^F, ^13^C, ^15^N quadruple-resonance cryoprobe. ^1^H-NMR spectra were acquired at a proton frequency of 700 MHz at pH or pD 7.4 (direct meter reading) at 25 °C. The protein concentration was *ca*. 0.5 mM. ^1^H-^13^C and ^1^H-^15^N heteronuclear single-quantum coherence (HSQC) spectra were acquired at respective natural abundance. Data were processed with Topspin 4.0.6 (Bruker Biospin) and analyzed with Sparky software (101).

### Glucagon cell-based assay

A HEK-293 cell line stably overexpressing GCGR was generated (Fig. S12). These cells enabled calibration of a glucagon-dependent cAMP production assay using the LANCE™ Ultra cAMP Kit (PerkinElmer, Waltham, MA) following manufacturer instructions.

### Insulin cell-based assay

Insulin activities were measured using an in-cell pIR immunoblotting as described (24, 100). This 96-well plate assay, exploiting a Chinese hamster ovarian (CHO) cell line overexpressing the IR (isoform B), enabled parallel measurement of insulin-dependent IR autophosphorylation (pIR/IR ratio).

### SQ rat assays

Insulin activities were measured *in vivo* as described (97). The assays were done by SQ injection in male diabetic Lewis rats, rendered diabetic by STZ (100). Glucagon activity was evaluated following SQ injection in normal male Lewis rats fasted for 4 h before the assay. All experiments were performed in accordance with protocols approved by the CWRU Institutional Care and Use Committee (IACUC).

### Hyperinsulinemic clamps

Sprague Dawley rats of 300-315 g body weight were anesthetized 2-5 days after arrival and underwent aseptic surgery for vascular catheter implantations into the left carotid artery (for blood glucose sampling) and right jugular vein (for insulin- and glucose infusions). BGCs, measured every 5-10 min, were maintained at 100-120 mg/dL for 90 min during the euglycemic clamp and at 60 mg/dL for 90 min during the hypoglycemic clamp. GIRs required to maintain glycemic targets were recorded. All experiments were performed in accordance with protocols approved by the Yale Institutional Care and Use Committee (IACUC).

### Statistical analyses

Statistical analyses were performed using GraphPad Prism (version 9.4). Data are presented as means ± standard error of the mean (SEM); values between different groups were compared with unpaired Student t-test or two-way ANOVA with repeated measures. P<0.05 was considered significant.

## Acknowledgments

N.V. is a Fulbright Scholar and acknowledges funding issued by Fulbright Chile and ANID. We thank S. Sullivan (Helmsley Charitable Trust), N.B. Phillips (Case Western Reserve University), S.M. Strano (MIT) and R. DiMarchi (Indiana University) for helpful discussions. We thank Y.-C. Chen (Indiana University) for advice regarding insulin cell-based assays, B. Dhayalan for advice regarding SPPS and fibrillation assays, and A. Ehnbom for MD simulations. We also thank S. Poordian (CWRU) and Y. Ding (Yale University) for their assistance with rat studies. M.A.W. acknowledges adjunct/courtesy appointments in the Department of Chemistry at Indiana University (Bloomington) and the Weldon School of Biomedical Engineering at Purdue University. We thank the IUSM Chemical Genomics Core for the use of instrumentation and the IUSM Center for Diabetes and Metabolic Diseases. This work was supported in part by The Indiana University Precision Health Initiative to M.A.J. and M.A.W., by grants to R.I.H. from the National Institutes of Health (R01DK101984, R01DK020495 and P30DK045735), and by grants to M.A.W. from the Leona M. and Harry B. Helmsley Charitable Trust (1902-03727) and National Institutes of Health (R01 DK040949 and DK127761).

## Author Contributions

N.V., F.I.-B., M.A.J., R.I.H. and M.A.W. conceptualized the program of research and designed experiments. N.V. performed chemical protein syntheses and purification under the guidance of M.A.J. and M.A.W., performed cell-based, CD and fibrillation assays, and undertook data analyses. NMR studies were designed and interpreted by N.V., Y.Y. and M.A.W. A.D.C., F.I.B. and M.A.W. provided insight into pathophysiologic aspects of diabetes mellitus in patients and animal models. SQ rat studies with respective data analyses were performed by R.G. under the guidance of F.I.-B. IV clamp studies in rats, with respective data analysis, were performed by N.T under the guidance of R.I.H., N.V. and M.A.W drafted the original manuscript; and N.V., F.I.-B., M.A.J., R.I.H., A.D.C. and M.A.W. reviewed and edited manuscript. The overall program of research was directed by M.A.W.

