## Supplemental material for "Ultra-stable insulin-glucagon fusion protein exploits an endogenous hepatic switch to mitigate hypoglycemic risk"

**This PDF file includes:**

Supporting text
Figures S1 to S27
Tables S1 to S6
SI References

### SI Appendix

#### Supplementary Methods

*Peptide synthesis*

Glucagon- and insulin analogs were chemically synthesized through solid-phase peptide synthesis (SPPS) using infrared or induction heating in traditional Fmoc/tBu (fluorenylmethyloxycarbonyl/tert-butyl) chemistry and standard DIC/6-Cl-HOBt (diisopropylcarbodiimide/6-Cl-hydroxybenzotriazole) activation/coupling cycles. Tribute™ or Chorus™ automated peptide synthesizers (Gyros Protein Technology, Tucson, AZ) were used to run 0.1-mmol scale syntheses using N,N'-dimethylformamide (DMF) as main solvent. Preprogrammed protocols were created to deliver 10 equivalents of a protected amino acid, DIC and 6-Cl-HOBt in couplings heated to 60 °C for 10 minutes (min). Fmoc deprotection was done in piperidine (20% in DMF) at 50 °C in two repetitions of 3 min. Pre-loaded resins were used: Fmoc-Asn(Trt)-ChemMatrix® resin was used for insulin analogs, Fmoc-Lys(Boc)-Wang for glucagon analogs carrying a C-terminal lysine and Fmoc-Thr(tBu)-Wang for synthesis of native glucagon. Peptides were cleaved from the resin and deprotected by treatment with 90% trifluoroacetic acid (TFA), 2.5% triisopropylsilane (TIPS), 2.5% water, 2.5% (ethylenedioxy)-diethenethiol) (DODT), and 2.5% anisole. After cleavage, peptides were precipitated with cold (4 °C) ethyl ether. Natural Fmoc-amino acids, preloaded ChemMatrix® resins and 6-Cl-HOBt were purchased from Gyros Protein Technology (Tucson, AZ). Piperidine, DIC, Fmoc-Or(Boc) and preloaded Wang resins were purchased from Chem-Impex International, Inc. (Wood Dale, IL). TFA, TIPS, DODT and anisole were purchased from Sigma-Aldrich (St. Louis, MO). DMF, dichloromethane (DCM), methanol and ethyl ether were purchased from Fisher Scientific (Waltham, MA).

*Folding and purification of insulin analogs*

Crude single-chain insulin polypeptides were dissolved (in reduced form) to a final concentration of 0.1 mM in a buffer containing 20 mM glycine, 2 mM cysteine hydrochloride, and 2 mM cystine dihydrochloride with final pH adjusted to 10.5. This solution was stirred open to air at 4 °C overnight. The reaction was monitored by analytical reversed-phase (rp) HPLC; when completed, it was quenched with HCl (5 N) at a final pH of ~2.0. The solution was then filtered (0.22 µm); the folded peptide was purified by preparative rp-HPLC using Waters 2545 Quaternary pumping system (Waters, Milford, MA) with detection by UV absorption. Purifications were performed on a C4 PROTO 300 Å (20x250 mm, 10 µm) column (Higgins Analytical Inc, Mountain View, CA) using 0.1% TFA in $H_2O$ as solvent A (aqueous solvent) and 0.1% TFA in acetonitrile (Fisher Scientific, Waltham, MA) as solvent B (organic modifier).

*Lactam-bond formation and glucagon purification*

Before cleavage from the resin, selective deprotection of residues with differential orthogonal protection was done to prepare for the amide-bond (lactam) formation. In the chosen positions, Lys(alloc) and Glu(allyl) amino acids were used, and the deprotection reaction followed a protocol previously reported by Ahn et al. (2001) (1). In brief, 24 equivalents of $PhSi_3$ (phenylsilane) and 0.24 equivalents of $Pd(PPh_3)_4$ (Tetrakis(triphenylphosphine)palladium(0)), purchased from Sigma-Aldrich (St. Louis, MO), were used in the presence of constant bubbling of nitrogen gas ($N_2$) or argon gas (Ar). This reaction was done twice for 30 min with thorough intervening washes using DMF and DCM. Lactam-bond formation was induced by addition of 12 equivalents of DIEA (N,N-diisopropylethylamine), 6 equivalents of HBTU (Hexafluorophosphate benzotriazole tetramethyl uranium) and 6 equivalents of 6-Cl-HOBt. The reaction was run for 2 hours, and the completion was verified by the ninhydrin test. Finally, Fmoc group was removed by a 20 min reaction with 20% piperidine in DMF.The peptide resin was again washed with DMF and DCM prior to cleavage from resin and deprotection of the remaining groups. DIEA and HBTU were purchased from Sigma-Aldrich (St. Louis, MO). Glucagon analogs were purified similarly to insulin analogs, except that a CLIPEUS C8 (20 × 250 mm, 5 µm) column (Higgins Analytical Inc, Mountain View, CA) was used.

*Trypsin-mediated ligation*

A peptide bond was introduced between the C-terminal Lys of glucagon analogs and the α-amino-group of the folded single-chain insulin (SCI; $Phe^{B1}$) as catalyzed in an aqueous-organic co-solvent by trypsin. In all glucagon analogs and SCIs, native Lys- and Arg residues were substituted by ornithine (Orn) to avoid internal tryptic cleavage sites. An approximately 1:1 molecular ratio was used for trypsin-mediated ligation, typically ~ 9 mg of SCI were dissolved along with ~3 mg of a glucagon analog in 200 µl of a mixed-solvent system containing N'N'-dimethylacetamide/1,4-butanediol/0.2 M Tris-acetate in a ratio of 35:35:30 v/v/v. The pH was adjusted to neutral with 2 µl of 4-methylmorpholine; the reaction was carried out for 48 hours. Finally, the fusion protein was purified by preparative rp-HPLC on a CLIPEUS C8 (20 × 250 mm, 5 µm) column (Higgins Analytical Inc, Mountain View, CA).

*Characterization of purity and identity*

Purity and identity of all peptides and fusion proteins was confirmed by LC/MS using a Finnigan LCQ Advantage Mass Spectrometer System (ThermoFisher Scientific, Waltham, MA) coupled to an Agilent 1100 Series HPLC system (Agilent, Santa Clara, CA) equipped with a TARGA C8 (4.6 × 250 mm, 5 µm) column (Higgins Analytical Inc, Mountain View, CA). Purifications were repeated, if necessary, until achieving >95% purity was achieved. Pooled fractions were lyophilized and stored at -20 °C until use. Protein concentrations were assessed based on UV absorption at λ = 280 nm as measured on a NanoDrop 1000 spectrophotometer (ThermoFisher Scientific, Waltham, MA). Extinction coefficients at λ = 280 nm were calculated using the Protein Calculator (Innovagen, PepCalc.com).

*Fibrillation Assays*

Resistance to peptide- or protein fibrillation was tested by measuring the lag time prior to initiation of the exponential phase of fibril formation through a thioflavin-T (ThT) binding assay. This assay is based on ThT fluorescence emission at 480 nm, upon binding to mature fibrils, when excited at 440 nm. This assay was done as previously reported (2, 3). In brief, peptides were dissolved in phosphate-buffered saline (PBS at pH 7.4) to a final concentration of 100 µM and mixed with ThT (Sigma-Aldrich, St. Louis, MO) to a final concentration of 16 µM. Aliquots of 250 µL per well were placed in a 96-well plate under agitation at 37 °C and 15-min interval measurements using a Synergy H1 automated microplate reader (BioTek, Winooski, VT). Since nucleation of fibrillation is a stochastic process, lag times are typically variable among "identical" starting samples. We thus evaluated at least 8-10 technical replicates from at least 2 independent experiments ("biologic replicates"). The lag time was considered as the point at which fluorescence was 5 times the background signal.

*CD Spectroscopy*

Far-ultraviolet CD spectra (255-190 nm) were obtained using a Jasco J-1500 CD spectropolarimeter equipped with temperature control and an automated titration unit (Jasco, Tokyo, Japan). Samples were prepared at a concentration of 25 mM protein in 10 mM $KH_2PO_4/K_2HPO_4$ and 50 mM KCl at pH 7.4. A 1-mm pathlength quartz cuvette was used to obtain spectra at 4, 25, 37 and 45 °C with a wavelength resolution of 0.5 nm and averaging time 30 sec per point. Baseline buffer-only CD spectra were subtracted from protein-containing spectra acquired at the same temperature. Estimates of secondary-structure content were calculated from normalized spectra using the SELCON-3 algorithm packaged with the CDPro spectral analysis software (4-6). Using an automatic titration unit, thermodynamic stability was measured by monitoring guanidine-induced unfolding at helix-sensitive wavelength 222 nm. Free unfolding energies ($\Delta G_u$) were inferred at 25 °C from fitting to a two-state modeling using non-linear least-squares regression as described (7, 8).

*NMR spectroscopy*

NMR data were recorded using a BRUKER 700 MHz spectrometer equipped with $^1H$, $^{19}F$, $^{13}C$, $^{15}N$ quadruple-resonance cryoprobe. $^1H$-NMR spectra were acquired at a proton frequency of 700 MHz at pH or pD 7.4 (direct meter reading) at 25 °C. The protein concentration was *ca*. 0.5 mM. $^1H$-$^{13}C$ and $^1H$-$^{15}N$ heteronuclear single-quantum coherence (HSQC) spectra were acquired at respective natural abundance. Data were processed with Topspin 4.0.6 (Bruker Biospin) and analyzed with Sparky software (9).

*Molecular dynamics simulations*

Molecular dynamics (MD) simulations were implemented based on a glucagon crystal structure (protomer obtained from Protein Databank [PDB] entry 1GCN) and run at different simulated temperatures (25, 37 and 50 °C) using the CHARMM General Force Field (CGenFF) (10) and CHARMM code (build c47a2) (11). The sequence was modified in PyMol (12) to resemble the $K^{13}$-$E^{17}$-lactam, except for the side-chain to side-chain bond. First, MolProbity (13) was used to remove all hydrogen atoms, and these were replaced with hydrogen atoms positioned using the electron-cloud method. To the 32-residue protein a water box was added using the CHARMM-GUI (14) with subsequent charge neutralization using 0.1 M NaCl. A cubic water-box size of 78 Å was obtained. For terminal groups, patches "ACP" and "CT2" were used: these do not induce additional unphysical charge at either flanking end. Titratable residues were inspected using the PROPKA tool (15). The structures were energy-minimized to achieve a relaxed structure using the steepest-descent algorithm in 250 steps with a printing frequency of the energy set to 10 steps. The initial step size for the minimization algorithms was set to 0.005 ps.

Constant pressure (1 atm) MD simulations were run for 200 ns. Two different modes were employed, one set with restraints and the other without. In each case timesteps were set to 0.002 ps, and the number of MD steps per 1 ns was set to 500,000. The restrained simulations had two residues ($Lys^{13}$, $Glu^{17}$) mimicking a lactam ring fully restrained in space ("*cons harm abso force 10.0 mass select*"). Each simulation ran using the same starting structure and water box but with different initial seeds. Simulations were run in three replicas. Analyses were performed using VMD (16), PyMol and USCF Chimera (17).

*Cell-based glucagon-activity assay*

To measure glucagon activity, HEK-293 cells were engineered to stably overexpress the glucagon receptor (GCGR). Using this cell line (designated HEK-293-GCGR cells), we standardized a cAMP-production assay using the LANCE™ Ultra cAMP Kit (PerkingElmer, Waltham, MA) (Fig S1-A). In this assay a plate of HEK-293-GCGR cells was grown to 70% confluence and then incubated in a Low-Glucose medium for 24 hours. The cells were then transferred to a 384-well plate (1000 cells/well) and incubated with the analogs for 30 min. Production of cAMP was measured following instructions of the LANCE™ Ultra cAMP Kit.

*Cell-based insulin-activity assay*

To measure insulin activity, we used an In-cell pIR immunoblotting based on an assay previously reported (3, 18), with modifications. In brief, a Chinese hamster ovary-derived (CHO) cell line that overexpresses insulin receptor (IR) was seeded in a 96-well block with clear bottom plate at density ~15000 cells per well and left to grow for 48 hours. Before running the assay, cells were incubated in Dulbecco's Modified Eagle Medium (DMEM) medium without glucose, pyruvate (Sigma-Aldrich, St. Louis, MO) or serum for 3-4 hours. Next, the analogs were applied in serial dilution to each well and incubated for 20 min at 37 °C. The reaction was stopped by aspiration and incubation with 150 µl of 3.7% formaldehyde (Fisher Scientific, Waltham, MA) in PBS for 20 min. The cells were then permeabilized using 200 µl of 0.1 % Triton-X-100 (Sigma-Aldrich, St. Louis, MO) in PBS for 20 min. Next, 200 µl of the Odyssey Blocking Buffer (LICOR, Lincoln, NE) was used as a non-specific blocking buffer with incubation for 1 hour with gentle agitation at room temperature. After such blocking, cells were incubated with 1:1000 dilution of the antibody anti-pTyr 4G10 (Sigma-Aldrich, St. Louis, MO) overnight at 4 °C. After washing, the secondary antibody anti-mouse IgG-R-Phycoerythrin (Sigma-Aldrich, St. Louis, MO) in a 1:100 dilution was added and incubated for 1 hour at room temperature. Fluorescence was measured at 578 nm after 496-nm excitation. DRAQ5 (ThermoFisher Scientific, Waltham, MA) was also used to measure 700-nm emission as a control to estimate cell number. Fluorescence was measured on a Synergy Neo2 plate reader (BioTek, Winooski, VT) (Fig S1-B).

*Rat subcutaneous injection assays*

Animals were maintained in an accredited facility of Case Western Reserve University School of Medicine. All procedures were approved by the Institutional Animal Care and Use Committee

(IACUC) office of the University. Animal care and use were monitored by the University's Veterinary Services.

*In vivo* insulin activity was measured as described (19). Measurements of the glucose-lowering effect of insulin analogs and fusion proteins were performed in male Lewis rats (average body mass of ~300 g) who were rendered diabetic by streptozotocin (STZ) as described (3); they were fasted for ~1 h prior to injection.. Analogs were dissolved in PBS (pH 7.4) and the doses intended were injected in ~100 µL/300 g rat. Rats were injected under the skin into the soft tissue in the posterior aspect of the neck. Following injection, blood-glucose concentrations were measured at set time points using a small drop of blood (~25 uL) obtained from the clipped tip of the rat's tail using a clinical glucometer (EasyMax® V Glucose Meter, Oak Tree Health, Las Vegas, NV).

The *in vivo* glycemic effects of glucagon analogs and fusion proteins were evaluated in normal male Lewis rats (250-350 g) that were fasted for 4 h before the injection. Glucagon analogs were prepared at 150 µg/ml in sterile 50mM Tris buffer (pH 8.0) and administered by subcutaneous (SQ) injections in a volume of ~200 uL/333g rat. Rats were grouped according to their $2^{nd}$ measured fasting blood-glucose concentrations to ensure similar starting values (time = 0). For thermal-stability assays, glucagon and analogs (115 µM in sterile 50 mM Tris buffer [pH 8.0]) were incubated at 45 °C with mild agitation; aliquots were taken at times 0, 1, 2 and 4 weeks for SQ testing in rats as above.

*Hyperinsulinemic clamps*

Male Sprague-Dawley rats weighing 300-315 g were purchased from Charles River Laboratories (Wilmington, MA) and housed in the Yale Animal Resource Center in temperature (22-23 °C) and humidity-controlled rooms. Animals, fed with standard rat chow and water *ad libitum*, were acclimatized to a 12-h light cycle. Procedures and protocols were approved by the Yale University Institutional Animal Care and Use Committee.

Animals were anesthetized 2-5 days after arrival and underwent aseptic surgery in which vascular catheters were implanted into the left carotid artery (for blood glucose sampling) and into the right jugular vein (for insulin and glucose infusions). The catheters of overnight-fasted rats were connected to infusion pumps in the morning of the study and then left undisturbed to minimize handling stress for at least 60 min before baseline sampling.

The euglycemic clamp was induced with 0.096 nmol/kg/min of the fusion protein plus variable infusion of 20% glucose. Fusion proteins were formulated in 15% dimethylsulfoxide (DMSO), 0.2% bovine serum albumin (BSA) in PBS (pH 7.4). Blood-glucose concentrations were measured every 5-10 min and maintained at the target range of 100-120 mg/dL for 90 min. The hypoglycemic clamp portion was initiated with 0.48 nmol/kg/min of the fusion proteins and co-infused with 20% glucose at variable rates. The plasma glucose concentrations reached 60 mg/dL 30 min after initiation, and they were maintained for the remaining 60-90 min. Serum collected at times -30, 0, 30, 60 and 90 min were used for glucagon- and epinephrine measurements by ELISA according to the manufacturer's protocol (Mercodia).

*Statistical analyses*

Statistical analyses were performed using GraphPad Prism software (version 9.4). Data are presented as the means ± SEM. Glucose infusion rate (GIR) changes during the clamp study were summed for each animal to provide an integrated *area under the curve* (AUC) relative to the baseline. Data values between different groups were compared with unpaired Student's t-test or two-way ANOVA with repeated measures. $P<0.05$ was considered significant.

#### Supplementary Results and Discussion

*SCI and lc-glucagon co-injection*

The fusion proteins described in this work inevitably have a 1:1 molar ratio (*i.e.*, glucagon:insulin analog). Because the respective analogs are less potent than are the native hormones, the effective activity ratio typically favored higher relative insulin activity. In control studies of the separate component analogs, an SQ-injection assay was performed in rats to test the glucodynamic response of the simultaneous injection, at different sites, of separate hormone analogs at a 1:1 ratio. Thus, the SCI was either tested alone or as a 1:1 ratio with lc-glucagon ($K^{13}$-$E^{17}$-lactam, see main text). As shown in Figure S2, there was no significant difference between SCI alone and when co-administrated with the glucagon analog, at the two doses tested, 14 nmol/kg (◆ for SCI, and ■ for co-administration) and 38 nmol/kg (◆ for SCI and ■ for co-administration) This was expected due to the acute activity of glucagon analogs as previously observed (Fig. 7A) and the functional "switch" to a more insulin-sensitive state of the liver under hyperglycemic conditions (20).

As an aside, we note that co-injection of distinct hormones would encounter potential drawbacks as a co-formulation, including different solubility properties in a formulation (21, 22) and, on SQ injection, respective durations of post-receptor signaling (23, 24). In addition, a fusion protein would in principle provide a single molecular entity for regulatory approval.

*Structure of glucagon analog*

To explore whether a lactam bond between glucagon side-chains 13-17 (with helix-compatible spacing [i, i+4]) might confer an increase in α-helix propensity, CD spectra (190-270 nm) were obtained as a function of trifluoroethanol (TFE) concentration; this organic co-solvent induces α-helical structure. Peptide analogs were studied in the presence or absence of the lactam bridge. In Figure S3-A are shown spectra of $K^{13}$-$E^{17}$-linear (—) and lc-glucagon (—) in 10% TFE. The analog carrying the lactam bond showed a deeper 222 nm signal, suggesting an increased α-helical content. Deconvolution to estimate helical content was done using CDPRO software with SELCON3 deconvolution algorithm (Fig. S3-B). This analysis provided evidence for an increased α-helix propensity for lc-glucagon (▲) compared with parent unconstrained peptide (■) at all TFE concentrations tested.

The CD experiments were extended by MD simulations. To this end, 200 ns MD simulations were performed, beginning with the crystal structure (1GCN) and modified with $K^{13}$, $E^{17}$ substitutions and with -EEK C-terminal extension. The simulations were run at 50 °C to allow for structural reorganization and denaturation. In one run the $K^{13}$ and $E^{17}$ side chains were positionally restrained to emulate a side-chain lactam bond, and this simulation showed a resilient α-helical structure with disordered structure at both ends of the molecule (Fig. S4). In another run the analog was simulated without such restraints leading to different structures at the end of the run, all with random coil structure in the middle of the molecule, breaking the α-helix near residues 13 and 17. These simulations suggest that the lactam bond can nucleate and stabilize local helical structure, "locking" an active local conformation and preventing non-native reorganization towards a β-sheet-rich fibrillar structure (Fig. S4). This is more evident when measuring the Phi and Psi angles of residues $Ser^{11}$, $Asp^{15}$, $Ala^{19}$ (2 residues before, in the middle and after the lactam restrain) as the MD simulations progress (Fig. S3-C to F). The Phi and Psi angles when the molecule is not restrained (Fig. S3-C and D), especially on $Ala^{19}$, change to values outside of the allowed for right-handed α-helix (-50° to -80° for Phi and -35° to -60° for Psi), which does not happen when the restraints are applied (Fig. S3-E and F).

We note that in the absence of TFE, as seen in Fig. S3-B, lactam-bridged and parent analogs exhibited similar spectra suggestive of a predominance of random coil. This finding was as expected based on the solution NMR structure of native glucagon (25) (Fig. S4). However, glucagon's active conformation is α-helical (26), which is also the structure it adopts when crystalized (27) and as a trimer (28) (Fig. S4). In addition to these conformations, glucagon can form amyloid-like fibrils with two co-existing full-length β-sheet formations (29) (Fig. S4).

*Structure of fusion protein*

The structure of the fusion protein in relation to the structures of its individual components was studied by CD and NMR spectroscopy. CD spectra (190-255 nm) are presented as molar ellipticity (θ) *per residue* (Fig. S5) and additionally, as molar ellipticity (θ) *per molecule* (Fig. 8). The latter are included to enable meaningful addition or subtraction of spectra: comparison of the spectra of the separate SCI (—) and lc-glucagon (—) with the spectrum of to the fusion protein (—) suggests that the moieties do not alter or perturb each other's structure when fused in a single polypeptide.

As a complementary CD-derived probe, respective stabilities were measured by using molar ellipticity (θ) at helix-sensitive wavelength 222 nm (a) as a function of temperature in the range 4-88 °C (Fig. S5-E) and (b) as a function of guanidine-HCl concentration (Fig. S5-F; see above). Trends were similar to those observed in studies of a related SCI (8). The SCI and the A/A FP each exhibited increased resistance to guanidine denaturation with respective free energies $\Delta G_u$ 4.2 ± 0.1 kcal/mol and 4.5 ± 0.1 kcal/mol (Table S5). These values are markedly higher than that of insulin *lispro* ($\Delta G_u$ 3.0 ± 0.1 kcal/mol). The glucagon analog does not exhibit a strong $[\theta]_{222\text{-nm}}$ signal, and so these data support independence of the SCI moiety's structure in the fusion protein.

An additional method to evaluate the structural independence of the component domains of the fusion protein was provided by high-resolution NMR spectroscopy. 1D $^1$H-NMR spectra of lc-glucagon and SCI are presented in relation to the fusion protein in Figure S6. All three proteins at the concentration of 0.4 mM showed good NMR behavior at PH 7.4 and at 25 ºC, as shown in the bottom panel of Figures S6-A to C. Well-resolved resonances of $Cys^{A11}$, $Leu^{B6}$ and $Gly^{B8}$ amide proton (magenta arrows) of SCI were not observed in the $^1$H-NMR spectrum of the fusion protein due to exchange line broadening ("conformational broadening"), even at the low concentration of 80 µM (5 times dilution, Fig. 7, top panel). Due to such broadening, only weak NOE cross-peaks were observed involving the amide protons of residues $Cys^{A11}$, $Leu^{B6}$ and $Gly^{B8}$ in the 2D NOESY spectrum of the fusion protein (Fig. S7). Such trends have previously been noted in NMR studies of insulin or insulin analogs (30, 31).

To further characterize the structure of A/A FP, 2D $^1$H-$^{15}$N HSQC NMR spectra were obtained at natural abundance. Overlay of respective $^1$H-$^{15}$N HSQC spectra of the individual moieties and the fusion protein are provided in Figures S8-A and B. The main-chain amide resonances of free lc-glucagon (*red* peaks in Fig. S8-A) are in the $^1$HN chemical shift region of 7.5-8.8 ppm, while the spectrum of free SCI shows typical NMR resonances of native-like insulin (31-33). $^1$H-$^{13}$C HSQC spectra (also acquired at natural abundance) provide correlations between a $^{13}$C atom and an attached proton ($^1$H) or protons via one-bond J-couplings. The aliphatic region is shown in Figures S8-C-D, and the aromatic region in Figures S8-E-F. Observation of corresponding cross-peaks provides evidence for the retention of the domain structures in the fusion protein, presumably due to their dynamic independence. The combined use of 2D homonuclear and nature-abundance heteronuclear NMR spectra allowed for near-complete heteronuclear NMR resonance assignments ($^1$H, $^{13}$C and $^{15}$N) of the separate lc-glucagon and SCI, enabling mapping of secondary structure based on pattern of secondary chemical shifts (36-38). (We note in passing that whereas the isolated glucagon analog exhibited random-coil-like $^1H_N$/$^{15}$N chemical shifts, trends in $^{13}C_\alpha$ and $^{13}C_\beta$ chemical shifts are considered to be more sensitive to protein secondary structure (37, 39)). In lc-glucagon, the α-helix-like segment comprises residues Y10-1D5, and other residues reveal random coil structure. Its $^{13}C_\alpha$ and $^{13}C_\beta$ chemical shift, and secondary chemical shift are summarized in Table S2. In comparison to the $^1$H-$^{13}$C HSQC spectra of the fusion protein, the resonances of the glucagon moiety revealed no changes or only slight shift, particularly in $^{13}$C dimension, implying that glucagon moiety in the fusion protein maintains similar conformation to the free glucagon.

Of particular interest, the aromatic resonances of glucagon moiety's Trp side chain in the fusion protein displayed large changes in $^1$H chemical shifts (Fig. S8-E), likely due to repacking of the indole ring near other aromatic residues when linked to the SCI. That the insulin moiety in the fusion protein maintained a native-like structure that was evidenced by characteristic patterns of (a) secondary chemical shifts in the fusion protein's $^1$H-$^{13}$C HSQC spectrum (Fig. S8-D, S8-F and Fig. S9-A) and (b) the long-range NOEs between methyl groups and aromatic rings in the hydrophobic core (Fig. S9-B).

*Alternative lactam bridges*

Insertion of alternative (i, i+4) side-chain lactam bond was investigated at positions previously described by Day et. al., (2009) (40); these alternative schemes focused on modifications that maintain or enhance glucagon activity in glucagon/GLP-1 dual agonists. To this end, a lactam bridge was inserted at residue positions 12-16, 16-20, 20-24 or 24-28 with respective pairs of Lys-Glu substitutions. Each bridge conferred protection from fibrillation for up to a month (Table S6); $K^{17}$-$E^{21}$ was the only lactam-bridged analog with less marked resistance to fibrillation (Fig. 4-A). In terms of *in vitro* potency, all of them also showed an increase when compared to the linear counterpart, suggesting that the lactam might be stabilizing the active conformation of glucagon (helical). The $K^{16}$-$E^{20}$-lactam, $K^{20}$-$E^{24}$-lactam and $K^{24}$-$E^{28}$-lactam analogs exhibited a potency similar to glucagon-EEK and twofold greater than $K^{13}$-$E^{17}$-lactam (lc-glucagon). To implement an insulin/glucagon activity ratio in the fusion protein that matched previous molar ratios biased towards less glucagon activity, the $K^{13}$-$E^{17}$-lactam analog was selected due to its 5-fold less potency compared to native glucagon (Table 1).

*Hypoglycemic response*

Hypoglycemic responses during the hypoglycemic clamp (Fig. S10-A) were studied by measuring glucagon (Fig. S10-B), C-peptide (Fig. S10-C), endogenous insulin (Fig. S10-D) and epinephrine (Fig. S10-E and F) levels. Results showed no significant difference between A/I FP (●), A/I FP (■) and the parent SCI (○) in epinephrine levels (abbreviations as in the main text), indicating that the difference in glucose infusion rates (GIR) observed (Fig. S10-A) is independent of hypoglycemic counter-regulation, and can be attributed to the endogenous glucose production caused by the glucagon moiety. As expected, A/I fusion proteins showed a GIR pattern similar to SCI-only, corroborating the lack of glucagon activity in the A/I fusion protein. Glucagon was, unfortunately, only possible to measure in the SCI-only group due to the cross-reactivity of the ELISA kit with our analog which caused levels higher than the detection limit (627 pg/ml). Endogenous insulin production was monitored by measuring serum insulin and C-peptide concentrations, which were low as expected, especially during the hypoglycemic phase.

*Inactive insulin and glucagon analogs*

To prepare the four fusion proteins described in Figure 4 (main text), we designed an inactive insulin and an inactive glucagon analog. To render the SCI inactive, a $Val^{A3} \rightarrow Leu$ substitution was incorporated. This modification corresponds to an inactive clinical mutation first described as insulin *Wakayama* (41). Apart from $Leu^{A3}$, the active and inactive SCIs were identical in sequence. To render the glucagon analog inactive, a side-chain lactam bridge was placed between positions 9 and 13, creating a $K^{9}$-$E^{13}$-lactam analog, previously reported as inactive by Ahn et al. (2001) (1). Observed *in vitro* and *in vivo* activities confirmed the predicted inactivities of these analogs (Fig. S11). In addition to these control analogs, an unrelated protein (hen egg white lysozyme) was tested *in vitro* for IR phosphorylation to measure the background signal of the assay and used to normalize AUC of the analogs (Table 1).

*Development of HEK293-GCGR cell line*

Unsuccessful attempts to standardize the cAMP assay using HepG2 cells (a hepatocarcinoma-derived cell line) led us to develop a stable cell line overexpressing the human glucagon receptor in HEK293 cells. The final cell line was monoclonal, and its GCGR expression was corroborated via Western blot. The GCGR gene carried a C-terminal Flag tag for easier identification and eventual flow cytometry selection (if needed). Western blotting showed a high expression of GCGR in the stable cell line relative to parent HEK293 cells when probed with anti-GCGR and anti-Flag antisera (Fig. S12)

*Analog purification and identification*

All analogs were purified by rp-HPLC with >95% purity; masses were characterized by LC-MS. The different sequence modifications, molecular masses, expected LC-MS ions and identified ions for all analogs are provided in Table S1. Respective HPLC chromatograms attesting to the purity and ESI-MS ions corroborating identities are presented in Fig. S13 to Fig. S27.

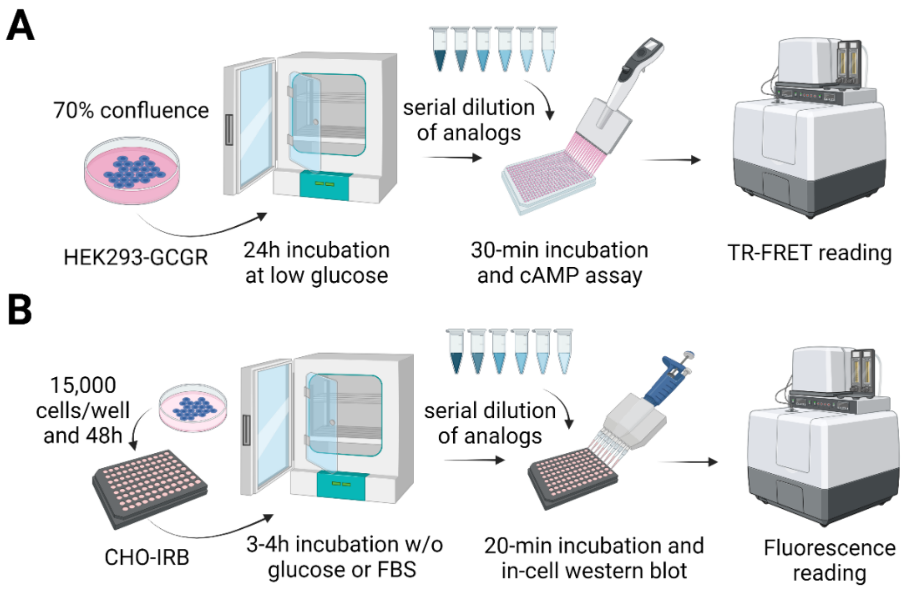

**Figure S1.** *Workflow for cell-based assays*. (A) Workflow for glucagon activity assay. A stable cell line that overexpresses glucagon receptor (HEK293-GCGR) was made. Cells are incubated in low glucose medium (1g/L) for 24h before assay. Serial dilutions of analogs are made and used for 30-min incubation with 1000 cells/well on a 384-well plate. cAMP is measured following the instructions of the LANCE™ Ultra cAMP Kit which requires TR-FRET reading. (B) Workflow for insulin activity assay. A stable cell line that overexpresses insulin receptor (CHO-IRB) previously developed was used (3, 18). 15,000 cells/well are placed on a 96-well plate and incubated for 48h. 3-4h before the assay the culture media is replaced with DMEM without glucose and without FBS. Serial dilutions of analogs are made and used for 20-min incubation followed by fixation, permeabilization, BSA blocking, and antibody incubation for an in-cell western blot assay. Total Tyr phosphorylation is assessed through Phycoerythrin fluorescence as a measure of IR phosphorylation. Figure created with BioRender.com.

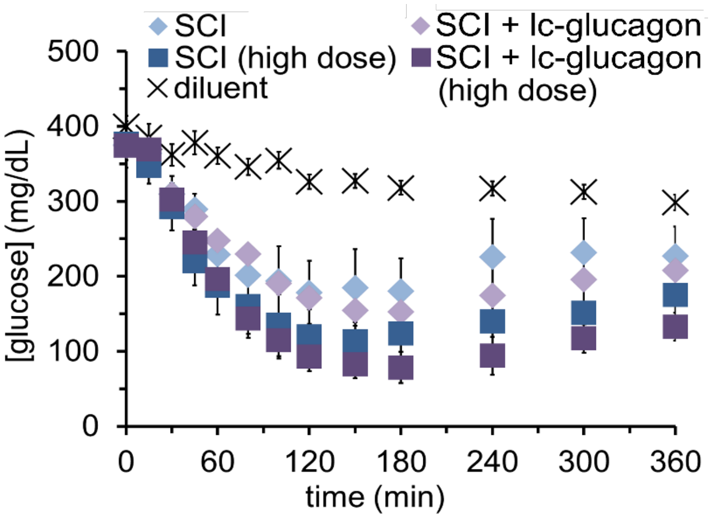

**Figure S2.** *In vivo evaluation of simultaneous injection of SCI and lc-glucagon*. The co-injection of both SCI and lc-glucagon on a 1:1 ratio was evaluated through SQ injection on diabetic rats (STZ) at two different doses, 14 nmol/kg (◆) and 38 nmol/kg (■). SCI alone at the same doses was used as a comparison, 14 nmol/kg (◆) and 38 nmol/kg (■). An n=6 was used per group. Error bars represent SEM.

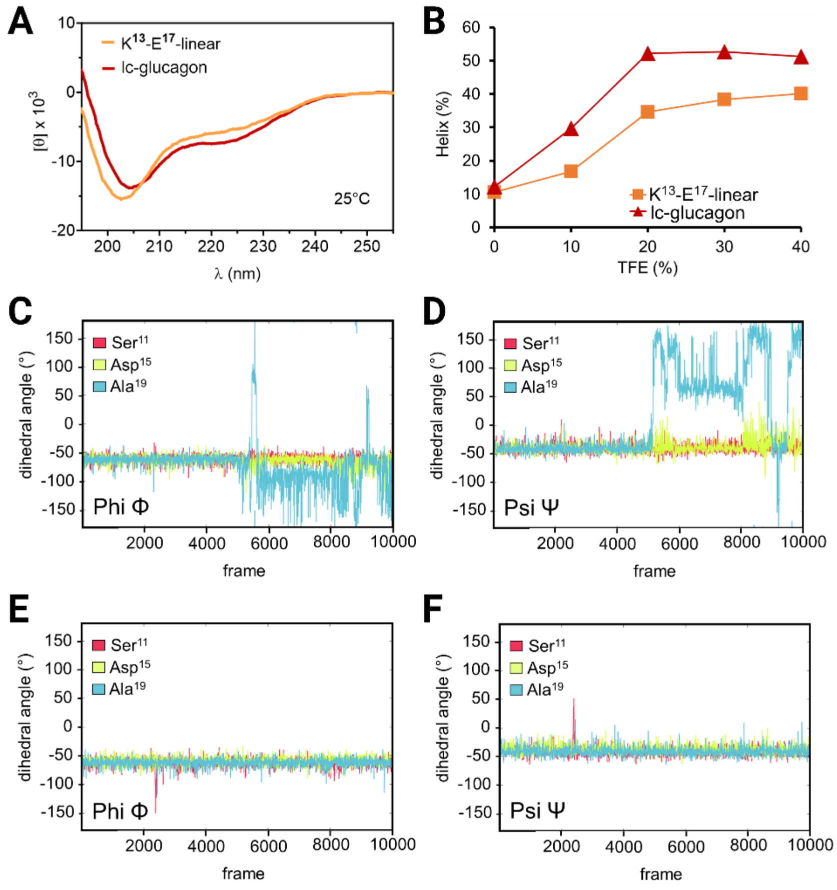

**Figure S3.** *$K^{13}$-$E^{17}$ lactam bond effect on helical structure.* (A) Circular dichroism wavelength scans (190 to 270 nm) of $K^{13}$-$E^{17}$-linear (—) versus lc-glucagon (—) in 10% TFE solution. (B) The helix percentage at different concentrations of TFE was calculated using CDPRO software with SELCON3 deconvolution algorithm. The data shows an increased α-helix propensity for the lc-analog (▲) versus $K^{13}$-$E^{17}$-linear (■) (C to F) MD simulations of 200ns were run from the crystal structure (1GCN) with $K^{13}$, $E^{17}$ modifications and -EEK c-terminal extension, at 50°C (to allow for structure reorganization). Two conditions were tested: with the $K^{13}$ and $E^{17}$ side chains positionally restrained to emulate lactam bond or without the restrain to emulate the linear analog. (C) Phi angles of residues $Ser^{11}$, $Asp^{15}$ and $Ala^{19}$ during the 200ns MD run when the restrain is not applied. (D) Psi angles of residues $Ser^{11}$, $Asp^{15}$ and $Ala^{19}$ during the non-restrained 200ns MD run. (C) Phi angles of residues $Ser^{11}$, $Asp^{15}$ and $Ala^{19}$ during the 200ns MD run when the restrain is applied. (D) Psi angles of residues $Ser^{11}$, $Asp^{15}$ and $Ala^{19}$ during the restrained 200ns MD run. Color codes on panels C-F is $Ser^{11}$ (■), $Asp^{15}$ (■) and $Ala^{19}$ (■).

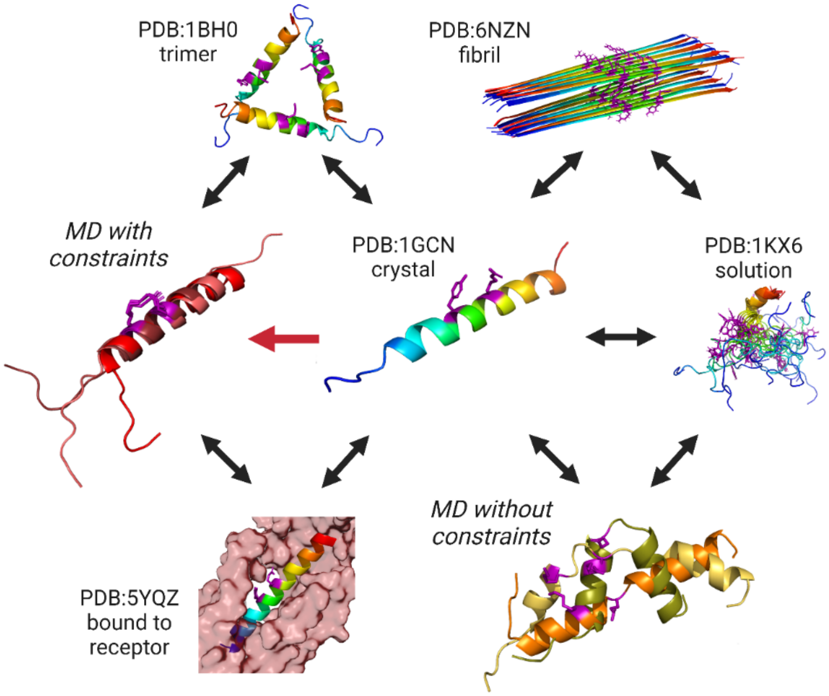

**Figure S4.** *Possible structures of glucagon.* Different reported structures of native glucagon or analogs plus the two 200ns MD simulations. Crystal structure (PDB ID: 1GCN), trimer structure (PDB ID: 1BH0) and receptor bound structure (PDB ID: 5YQZ) are mostly α-helical, while the solution NMR structure (PDB ID: 1KX6) is mostly random coil and the fibril structure (PDB ID: 6NZN), is completely β-sheet. All reported structures are shown with rainbow coloring from red (N-terminus) to blue (C-terminus) and with side-chains of residues 13 and 17 highlighted in purple. MDs were run from the crystal structure (1GCN) with $K^{13}$, $E^{17}$ modifications and -EEK c-terminal extension, at 50°C (to allow for structure reorganization). Two conditions were tested, with $K^{13}$ and $E^{17}$ side chains positionally restrained to emulate lactam bond (left) or without restraint (bottom right). Three representative frames from the last 20ns are shown structurally aligned. MD with restrain showed resilient α-helical structure.

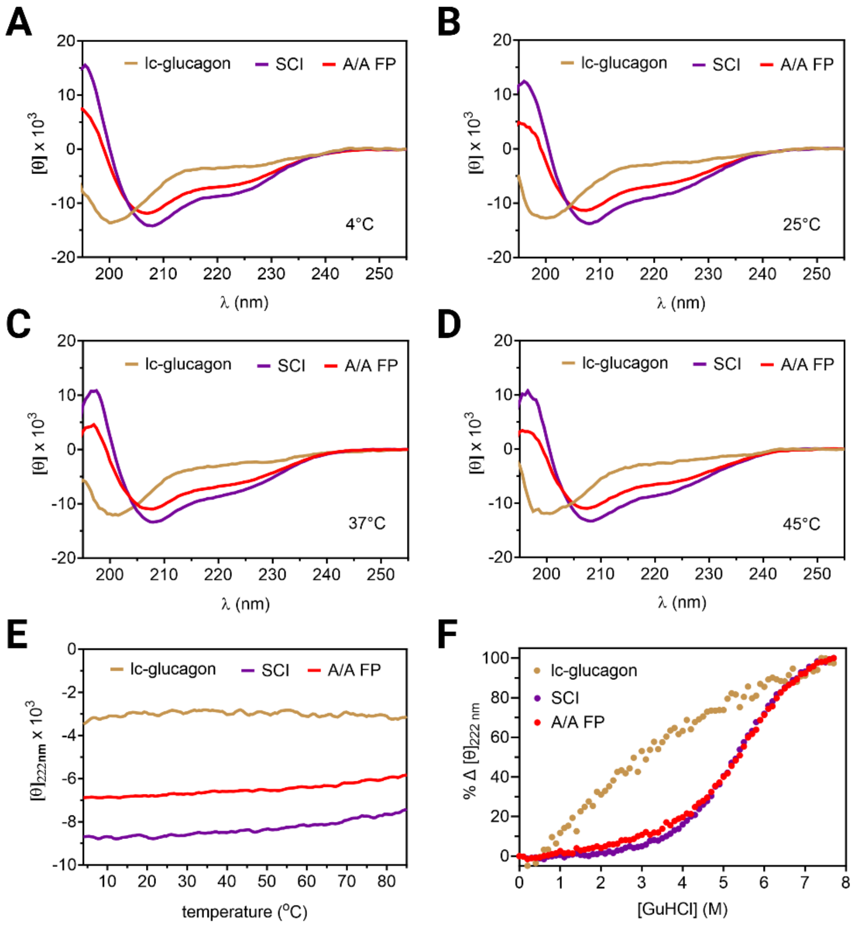

**Figure S5.** *CD studies on lc-glucagon, SCI and A/A FP.* (A-D) Circular dichroism wavelength scans (190 to 255 nm, molar ellipticity [θ] per residue) at 4°C, 25°C, 37°C and 45°C show a mostly random coil pattern for the glucagon analog and a more α-helical structure for the SCI analog. (E) The molar ellipticity [θ] at 222 nm, as a probe for helicity, was measured from 4°C to 88°C, showing a thermostable structure for SCI and A/A FP without major signal reduction below 45°C. (F) Guanidine denaturation studies monitored by molar ellipticity [θ] at 222 nm show a highly stable structure comparable to reported SCIs (8, 42). Color code for all panels is: lc-glucagon (—), SCI (—) and A/A FP (—).

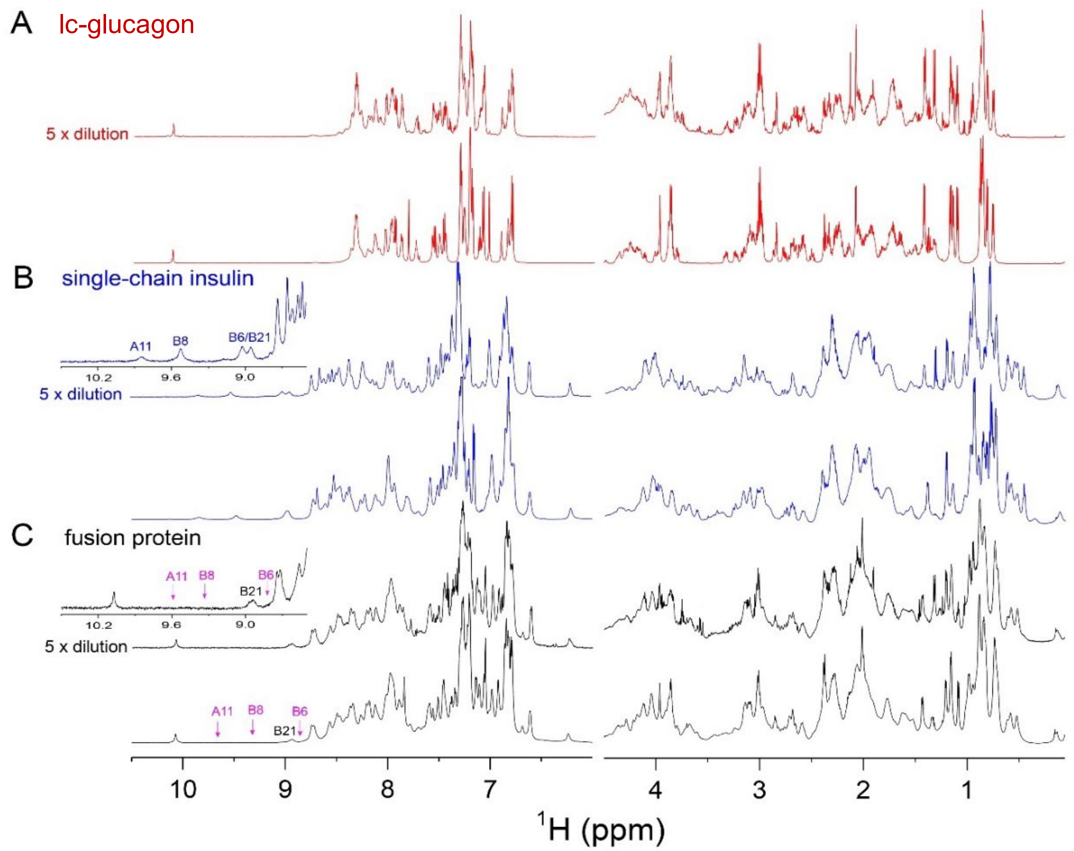

**Figure S6.** Stack plot of 1D proton spectra. (A) 1D $^1$H-NMR spectra of $K^{13}$-$E^{17}$-lactam glucagon (red); spectra were acquired at 0.5 mM (bottom panel) and 0.1 mM (top panel, 5 times dilution) of concentration. (B) 1D $^1$H-NMR spectra of the analog SCI (blue) acquired at 0.4 mM (bottom panel) and 80 $\mu$M (top panel, 5 times dilution) of concentration. (C) 1D $^1$H-NMR spectra of A/A FP (black) acquired at 0.4 mM (bottom panel) and 80 $\mu$M (top panel, 5 times dilution) of concentration. Left panel: aromatic and amide proton region; right panel: aliphatic proton region. Expanded spectra of single-chain insulin and A/A FP acquired with 80 $\mu$M were inserted into left of the spectra. Amide proton resonances of $Cys^{A11}$, $Gly^{B8}$ and $Leu^{B6}$ residues in down field in A/A FP were not observed due to spectral line broadening. Magenta arrows indicated resonances of $Cys^{A11}$, $Gly^{B8}$ and $Leu^{B6}$ that were observed in the NOESY spectrum of fusion protein. Spectra were acquired at a $^1$H frequency of 700 MHz in 90/10 ($H_2O$/$D_2O$) ratio at pH 7.4 (direct meter reading) and at 25 °C.

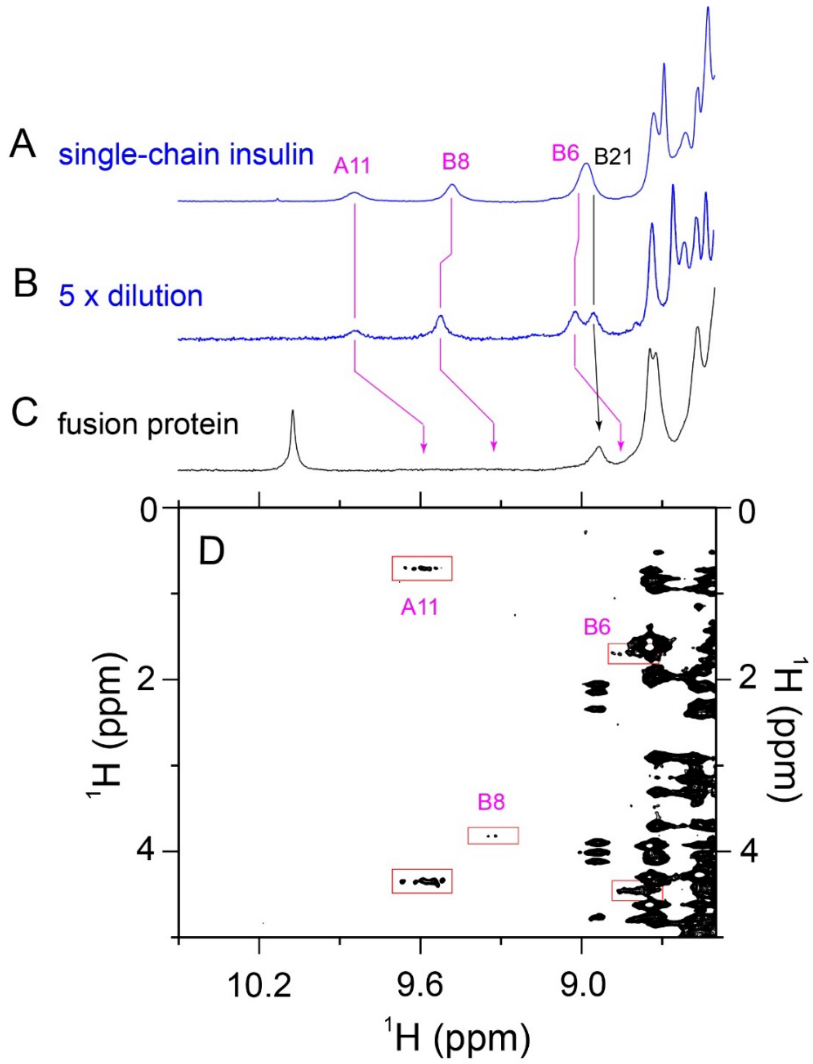

**Figure S7.** Down field region of 1D $^1$H-NMR spectra of SCI at protein concentration of 0.4 mM (blue, panel A) and 80 μM (blue, panel B), and that of fusion protein (A/A FP) at 0.4 mM (black, panel C). Amide proton resonances of $Cys^{A11}$, $Gly^{B8}$ and $Leu^{B6}$ residues in down field in the fusion protein were not observed due to spectral line broadening that showed very weak NOE cross peaks in the NOESY spectrum. Spectra were acquired at a $^1$H frequency of 700 MHz in 90/10 ($H_2O$/$D_2O$) ratio at pH 7.4 (direct meter reading) and at 25 °C.

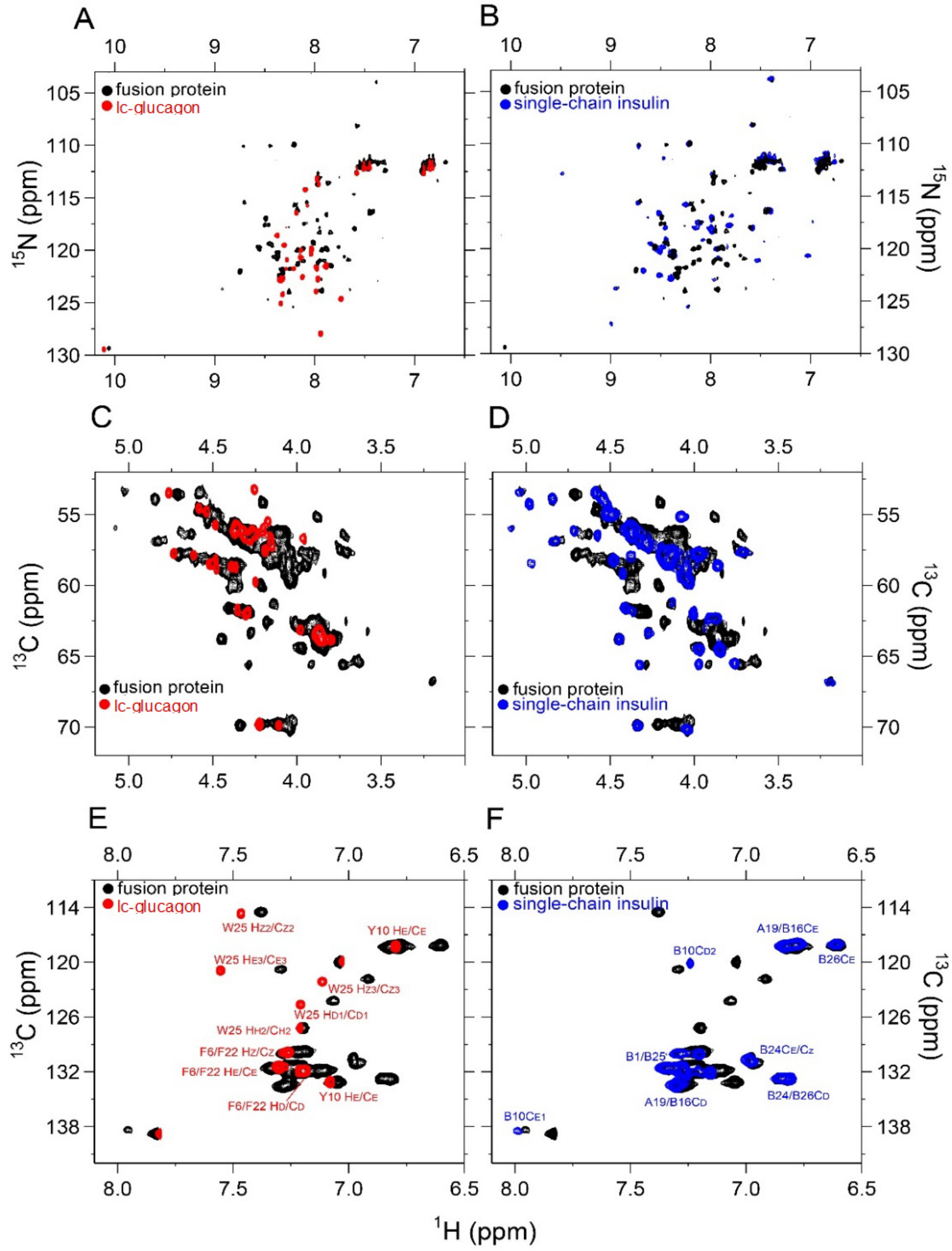

**Figure S8.** Comparison of nature abundance HSQC spectra of lc-glucagon, SCI analog and fusion protein (A/A FP). (A) Overlay of $^{1}$H-$^{15}$N HSQC spectra of A/A FP (black) and lc-glucagon (red). (B) Overlay of $^{1}$H-$^{15}$N HSQC spectra of A/A FP (black) and SCI analog (blue). (C) $^{1}$H$\alpha$-$^{13}$C$\alpha$ HSQC spectral overlay of A/A FP (black) and lc-glucagon (red). (D) $^{1}$H$\alpha$-$^{13}$C$\alpha$ HSQC spectral overlay A/A FP (black) and SCI (blue). Aromatic $^{1}$H-$^{13}$C HSQC spectral overlay of (E) A/A FP (black) and lc-glucagon (red) and (F) A/A FP (black) and single-chain insulin (blue). The spectral assignments were labeled in red for glucagon and in blue for single-chain insulin. Spectra were acquired at a $^{1}$H frequency of 700 MHz at pH 7.4 (direct meter reading) and at 25 °C.

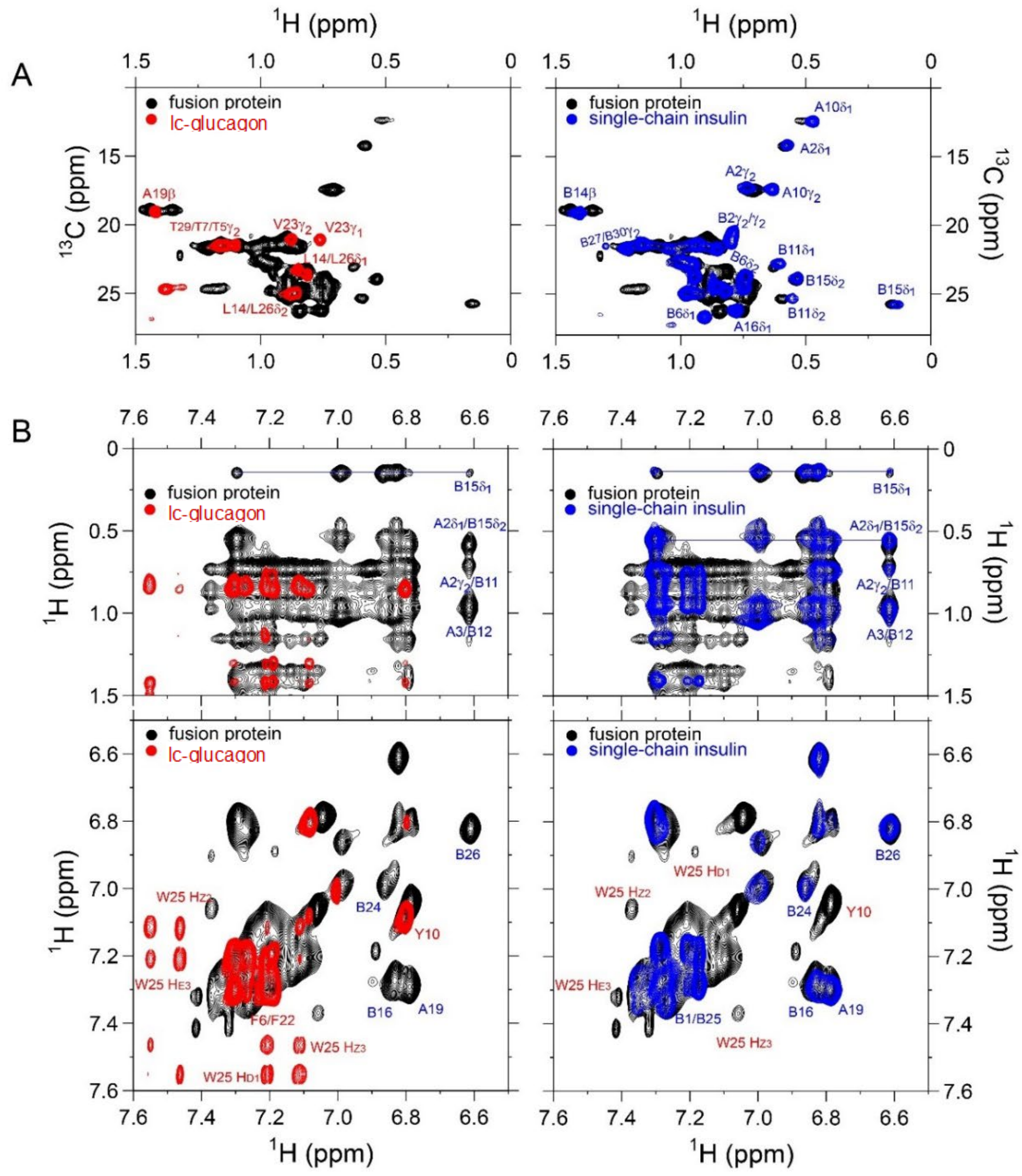

**Figure S9.** (A) Overlay of methyl $^1$H-$^{13}$C HSQC spectra: left panel: fusion protein (A/A FP) (black) and lc-glucagon (red); right panel: fusion protein (A/A FP) (black) and SCI analog (blue). (B) long-range signature NOEs between methyl protons and aromatic protons (top panel) and aromatic TOCSY spectra (bottom panel); left panel: overlay of fusion protein (A/A FP) (black) and lc-glucagon (red); right panel: overlay of fusion protein (A/A FP) (black) and single-chain insulin analog (blue). The spectral assignments were labeled in red for the glucagon analog and in blue for the SCI analog. Spectra were acquired at a $^1$H frequency of 700 MHz in $D_2O$ at pH 7.4 (direct meter reading) and at 25 °C.

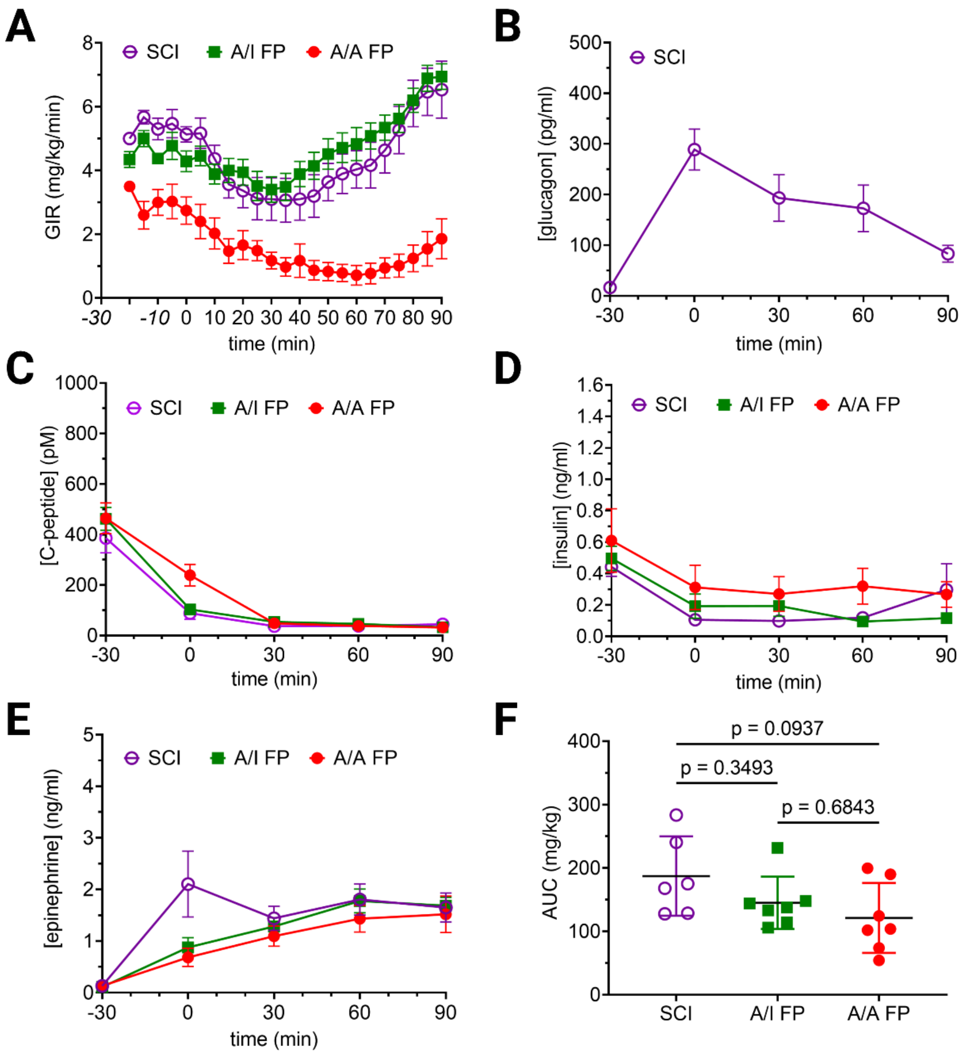

**Figure S10.** *Hypoglycemic response*. (A) GIR of the hyperinsulinemic hypoglycemic clamp (from Fig.8-D) on A/A FP (●), A/I FP (■) and SCI (○) (as control). (B) Glucagon levels in the SCI group were measured every 30 min. Glucagon levels were not possible to measure in A/A FP and A/I FP due to cross-reaction with the lc-glucagon leading to oversaturation of signal (above detection limit). (C) C-peptide levels measured on blood samples from the hypoglycemic clamp on all groups showed an expected decrease under hypoglycemic conditions. (D) Endogenous insulin levels measured on blood samples from the hypoglycemic clamp on all groups showed an expected decrease under hypoglycemic conditions. (E) Epinephrine levels on blood samples showed similar hypoglycemic counter-regulation in all groups. (F) AUC (0-90 min) calculated from (E).

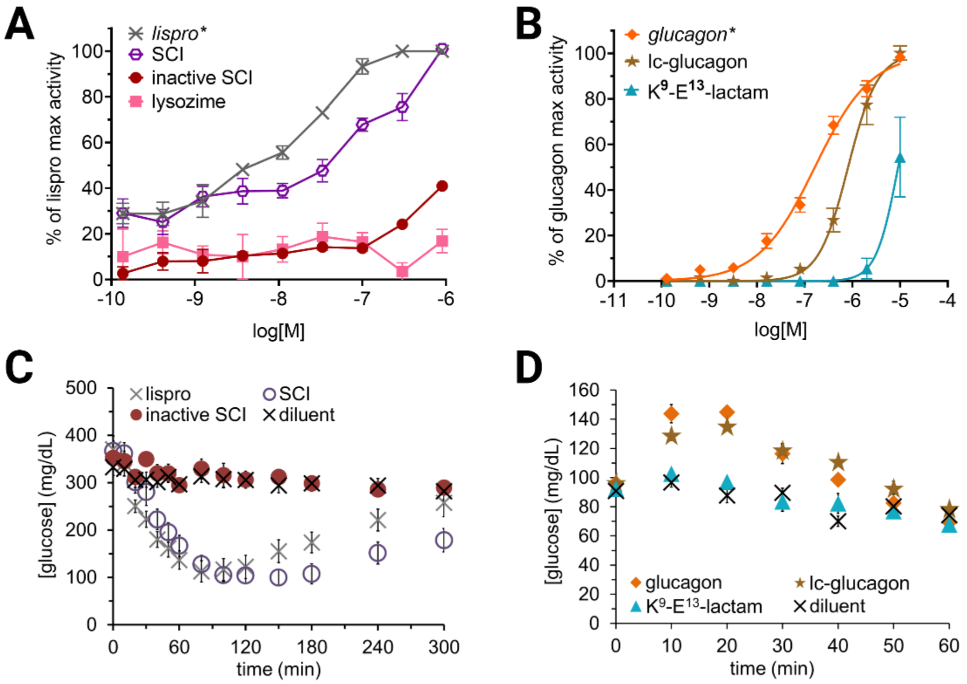

**Figure S11.** *Biological evaluation of inactive analogs*. (A) In vitro insulin activity assay comparing SCI (○) vs. inactive SCI (Leu$^{A3}$-SCI) (●) where total Tyr phosphorylation was measured through an in-cell western blot, as an indirect way of measuring insulin receptor phosphorylation. A CHO-IRB cell line was used. Lysozyme (■) was added as a control to quantify background signal. (B) In vitro glucagon activity assay comparing lc-glucagon (active) (★) vs. K$^9$-E$^{13}$-lactam (inactive) (▲) as measured by cAMP production using the LANCE™ Ultra cAMP Kit. (C) In vivo insulin activity (lowering of glucose levels) comparing *lispro* (×), SCI (○) vs. inactive SCI (Leu$^{A3}$-SCI) (●) as measured in diabetic rats (STZ) for each analog through SQ injection of lispro (8.6 nmol/kg) or SCI analogs (15.6 nmol/kg). (D) In vivo glucagon activity (increase in glucose levels) comparing glucagon (◆), lc-glucagon (active) (★) vs. K$^9$-E$^{13}$-lactam (▲) as measured in normal rats for each analog through SQ injection and comparison to native glucagon (12 nmol/kg), lc-glucagon (23 nmol/kg), or K$^9$-E$^{13}$-lactam (40 nmol/kg). Error bars represent SEM; n = 5 per group in panels C and D, respectively.

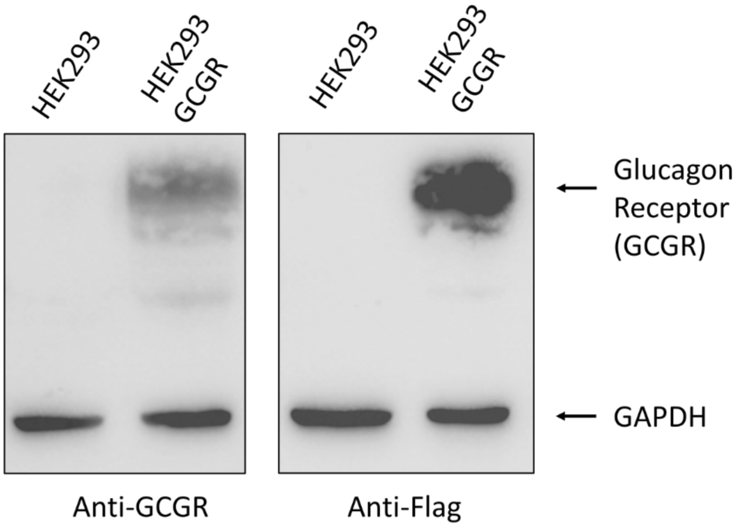

**Figure S12.** *Western blot evaluation of HEK-GCGR*. A stable, monoclonal, cell line that overexpresses glucagon receptor (GCGR) was developed from HEK293 cells, and it was evaluated by Western Blot. GCGR gene carried a Flag Tag at the C-terminus allowing for the evaluation through Anti-GCGR and Anti-Flag antibodies. Anti-GAPDH was used as a loading control and HEK293-GCGR was compared to original HEK293.

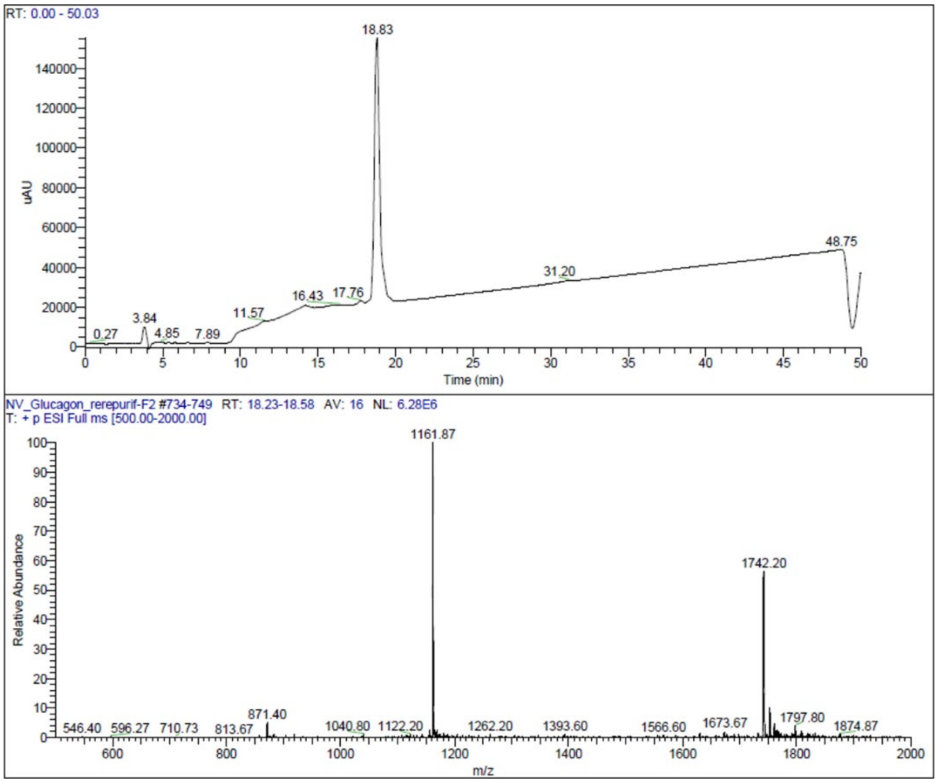

**Figure S13.** *LC/MS analysis of glucagon*. Top panel is the HPLC chromatogram monitored by UV absorbance at 215nm. Bottom panel shows the ESI-MS spectra of positive ions.

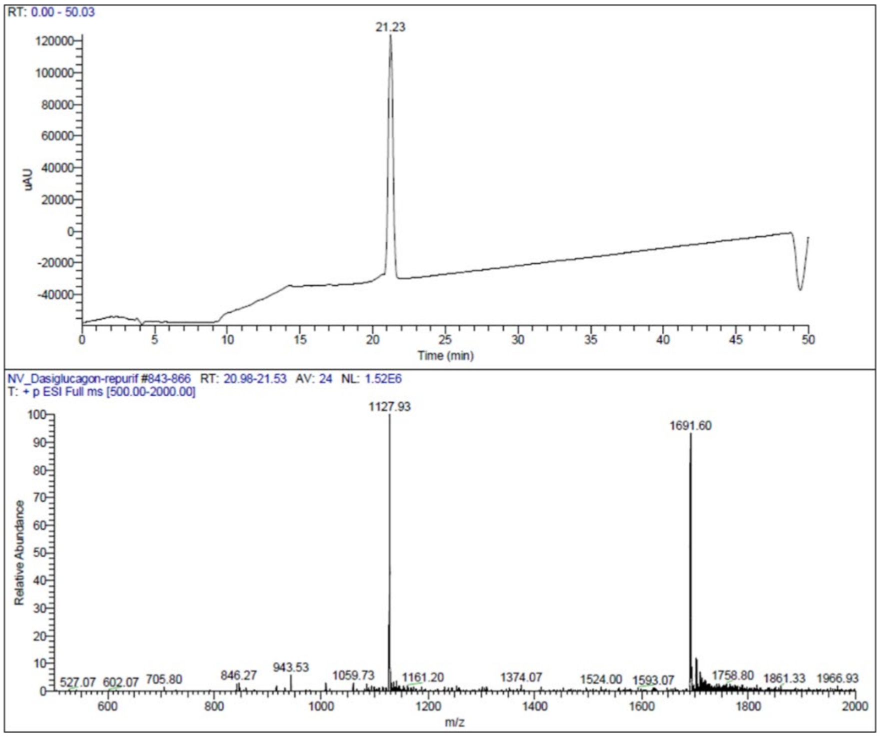

**Figure S13.** *LC/MS analysis of dasiglucagon*. Top panel is the HPLC chromatogram monitored by UV absorbance at 215nm. Bottom panel shows the ESI-MS spectra of positive ions.

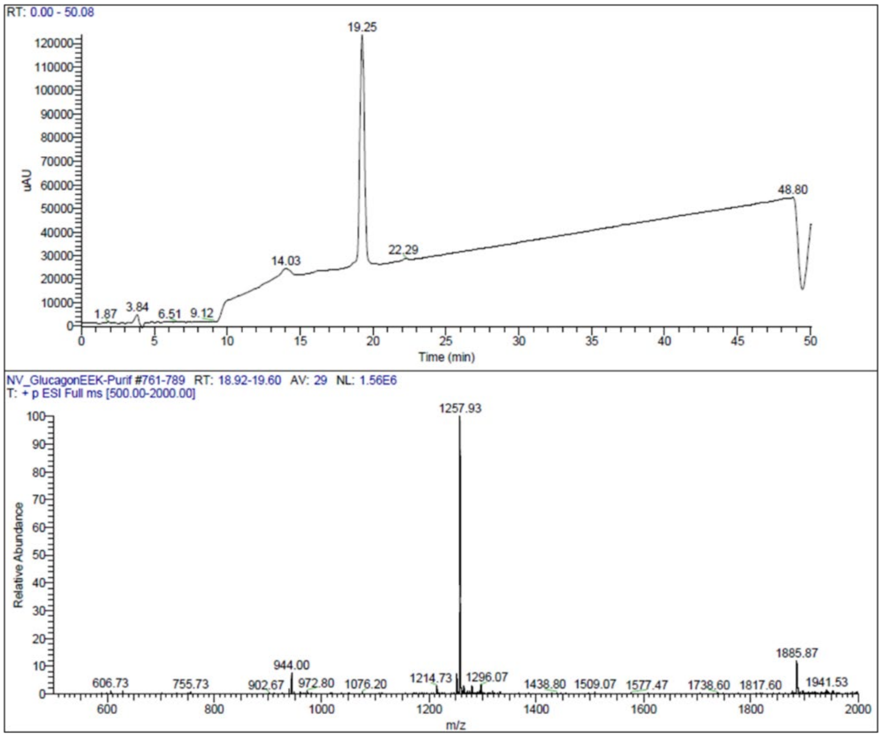

**Figure S15.** *LC/MS analysis of glucagon-EEK analog*. Top panel is the HPLC chromatogram monitored by UV absorbance at 215nm. Bottom panel shows the ESI-MS spectra of positive ions.

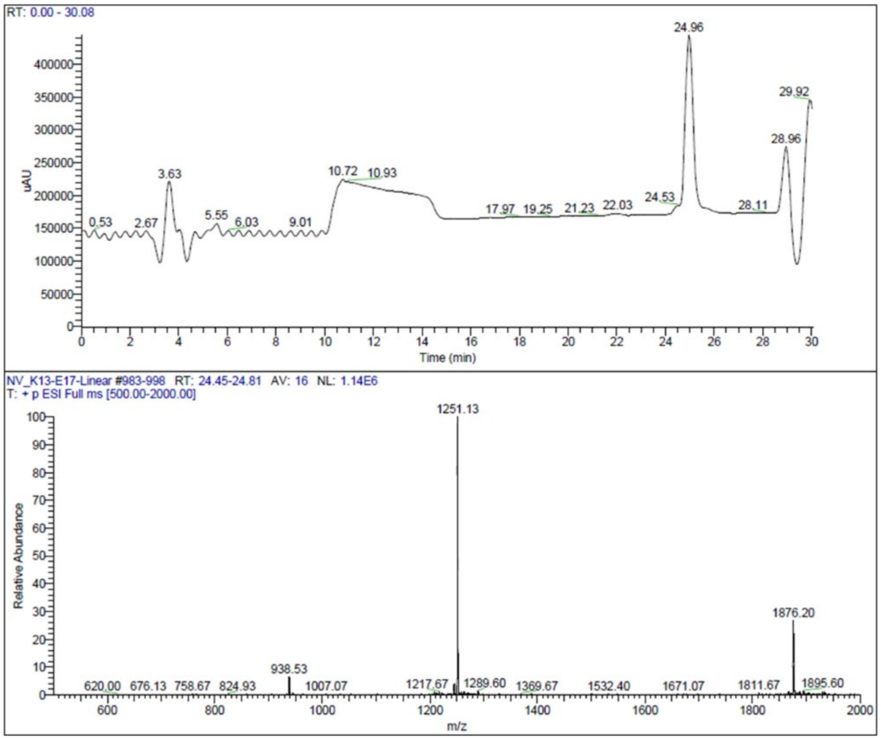

**Figure S16.** *LC/MS analysis of $K^{13}$-$E^{17}$-linear glucagon analog*. Top panel is the HPLC chromatogram monitored by UV absorbance at 215nm. Bottom panel shows the ESI-MS spectra of positive ions.

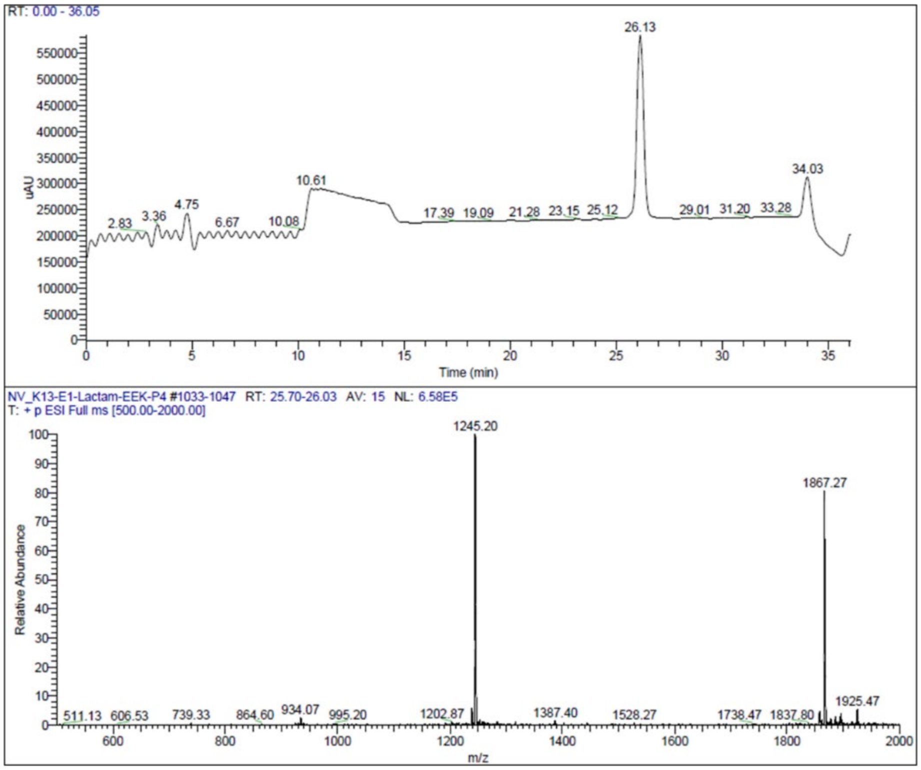

**Figure S17.** *LC/MS analysis of lc-glucagon ($K^{13}$-$E^{17}$-lactam glucagon analog)*. Top panel is the HPLC chromatogram monitored by UV absorbance at 215nm. Bottom panel shows the ESI-MS spectra of positive ions.

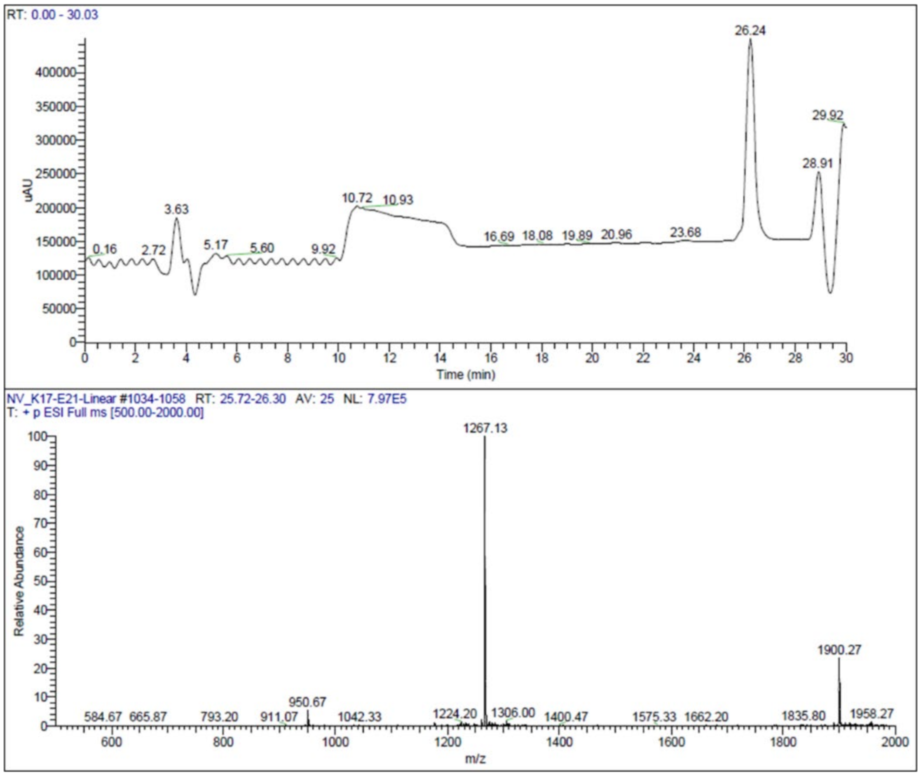

**Figure S18.** *LC/MS analysis of* $K^{17}$*-*$E^{21}$*-linear glucagon analog*. Top panel is the HPLC chromatogram monitored by UV absorbance at 215nm. Bottom panel shows the ESI-MS spectra of positive ions.

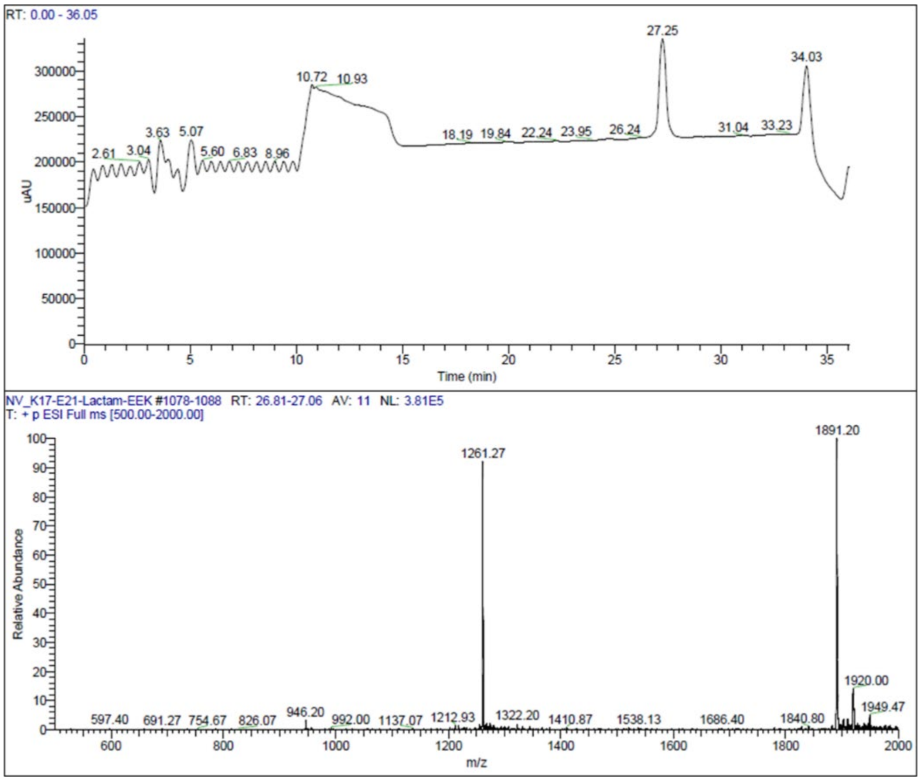

**Figure S19.** *LC/MS analysis of $K^{17}$-$E^{21}$-lactam glucagon analog*. Top panel is the HPLC chromatogram monitored by UV absorbance at 215nm. Bottom panel shows the ESI-MS spectra of positive ions.

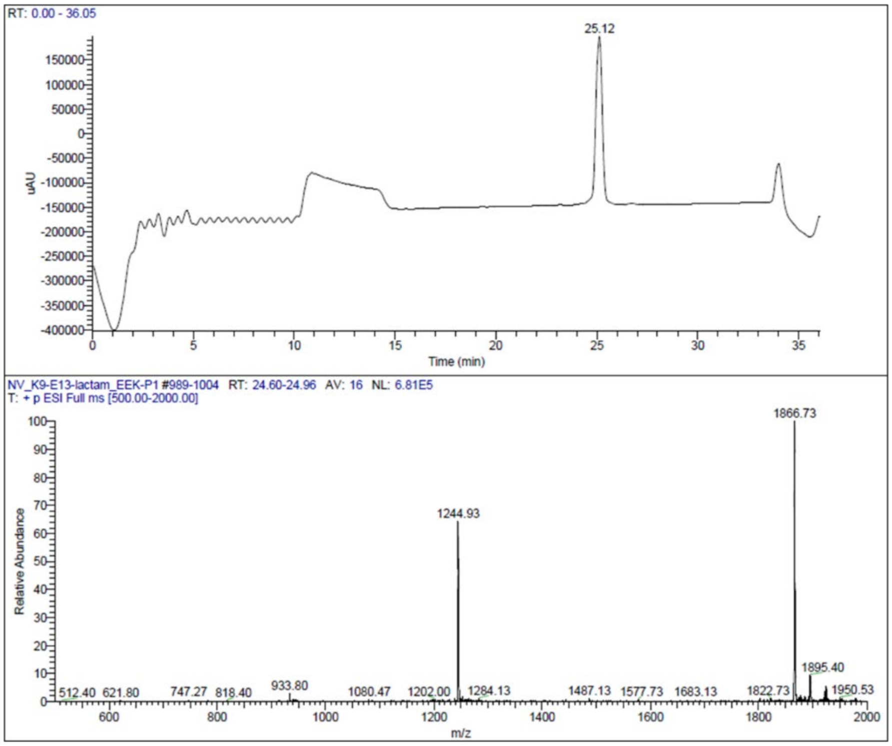

**Figure S20.** *LC/MS analysis of $K^9$-$E^{13}$-lactam glucagon analog*. Top panel is the HPLC chromatogram monitored by UV absorbance at 215nm. Bottom panel shows the ESI-MS spectra of positive ions.

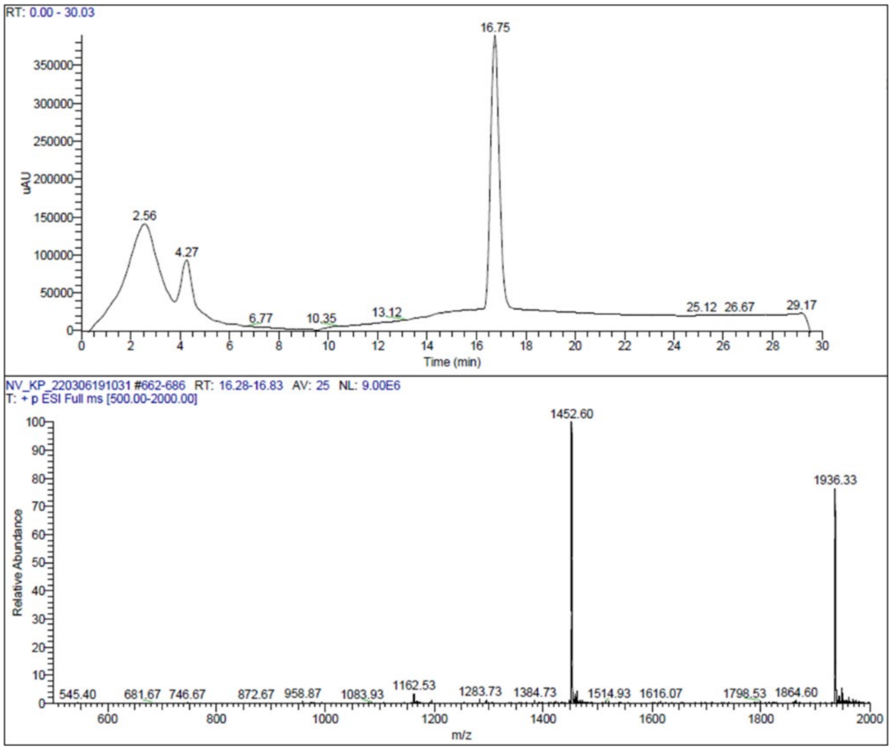

**Figure S21.** *LC/MS analysis of lispro insulin analog*. Top panel is the HPLC chromatogram monitored by UV absorbance at 215nm. Bottom panel shows the ESI-MS spectra of positive ions.

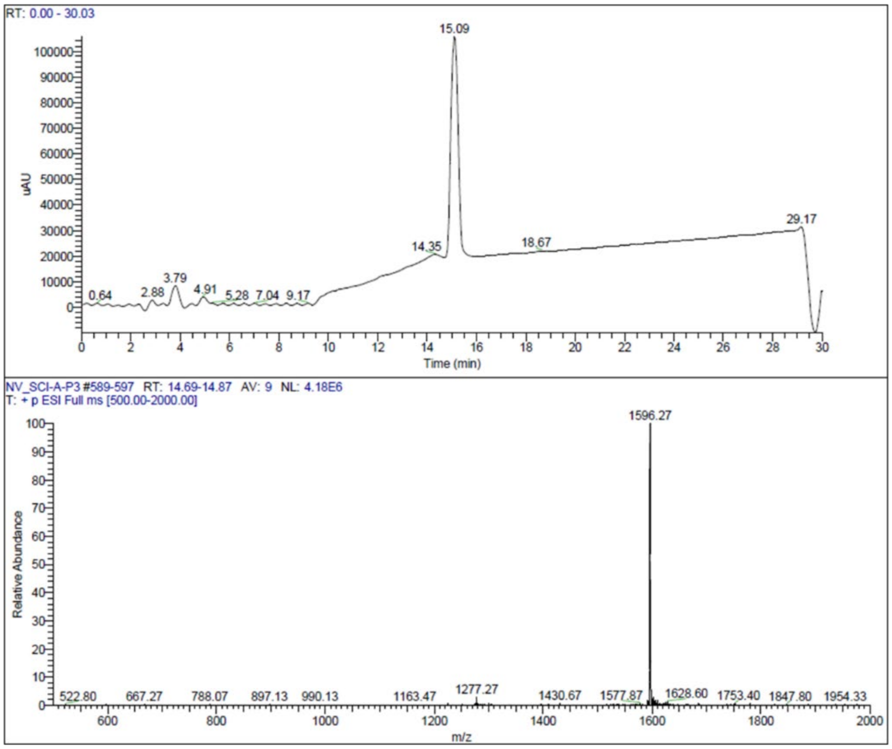

**Figure S22.** *LC/MS analysis of SCI analog*. Top panel is the HPLC chromatogram monitored by UV absorbance at 215nm. Bottom panel shows the ESI-MS spectra of positive ions.

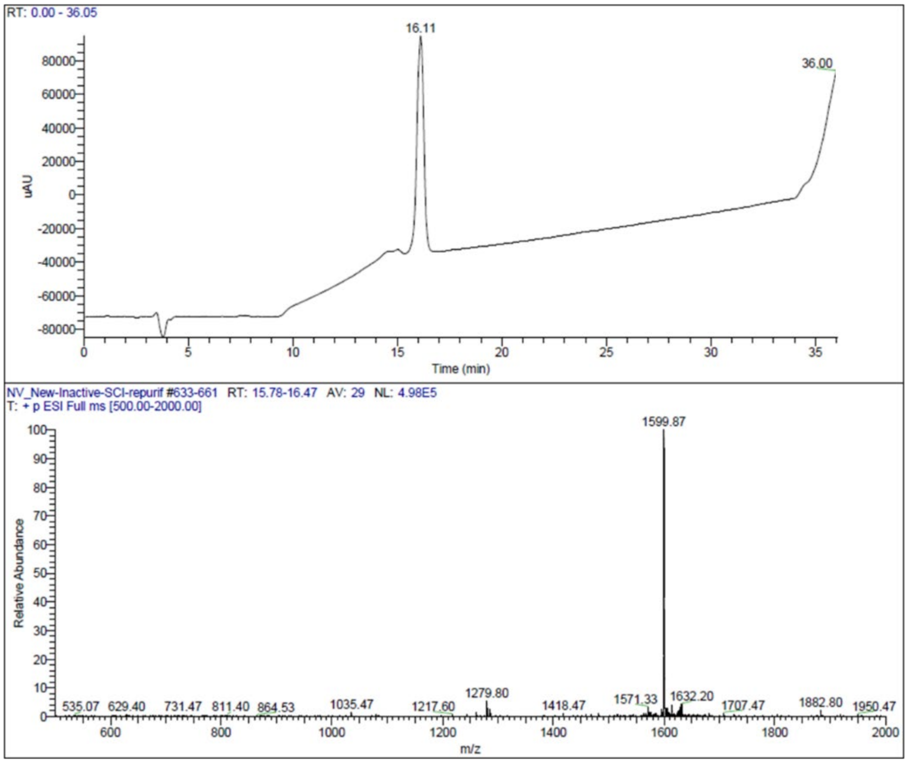

**Figure S23.** *LC/MS analysis of Inactive SCI analog*. Top panel is the HPLC chromatogram monitored by UV absorbance at 215nm. Bottom panel shows the ESI-MS spectra of positive ions.

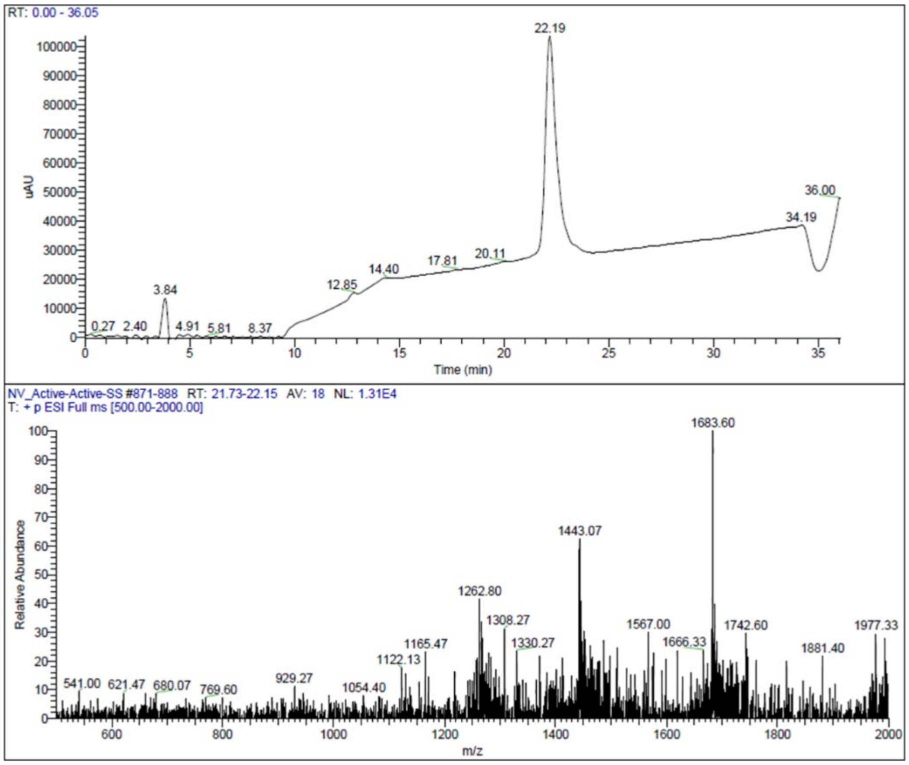

**Figure S24.** *LC/MS analysis of Active/Active FP*. Top panel is the HPLC chromatogram monitored by UV absorbance at 215nm. Bottom panel shows the ESI-MS spectra of positive ions. Due to the molecular weight and the m/z limit on the instrument (2000), the ions observed are are 1683.6 (M+6/6), 1443.07 (M+7/7) and 1262.8 (M+8/8),, which are less abundant, causing a lower signal-to-noise ratio.

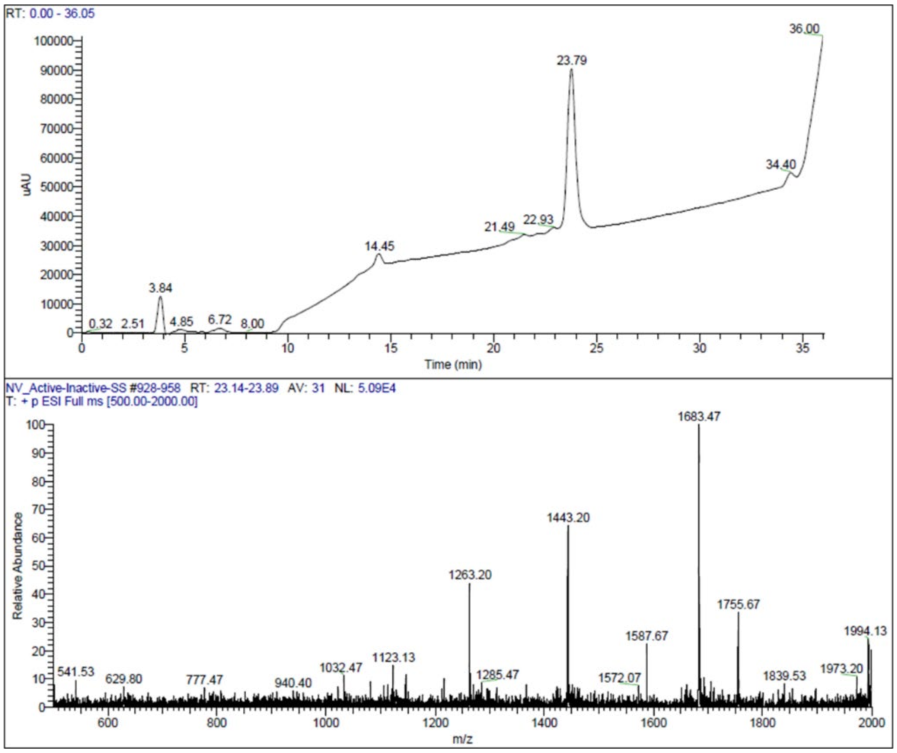

**Figure S25.** *LC/MS analysis of Active/Inactive FP*. Top panel is the HPLC chromatogram monitored by UV absorbance at 215nm. Bottom panel shows the ESI-MS spectra of positive ions. Due to the molecular weight and the m/z limit on the instrument (2000), the ions observed are 1683.47 (M+6/6), 1443.20 (M+7/7) and 1263.2 (M+8/8), which are less abundant, causing a lower signal-to-noise ratio.

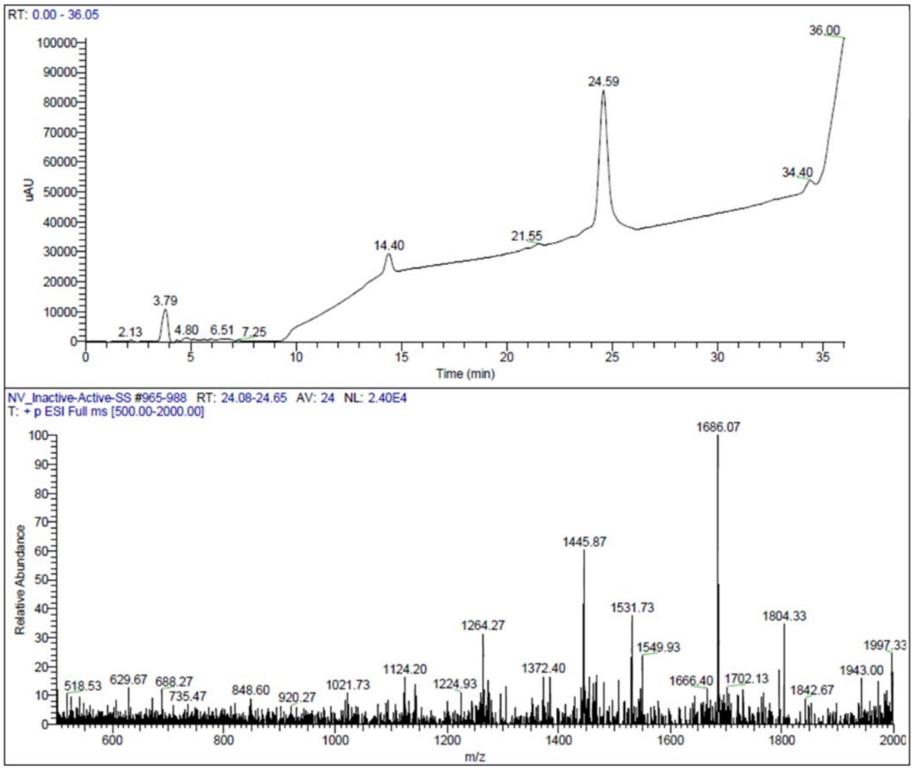

**Figure S26.** *LC/MS analysis of Inactive/Active FP*. Top panel is the HPLC chromatogram monitored by UV absorbance at 215nm. Bottom panel shows the ESI-MS spectra of positive ions. Due to the molecular weight and the m/z limit on the instrument (2000), the ions observed are 1686.07 (M+6/6), 1445.87 (M+7/7) and 1264.27 (M+8/8), which are less abundant, causing a lower signal-to-noise ratio

.

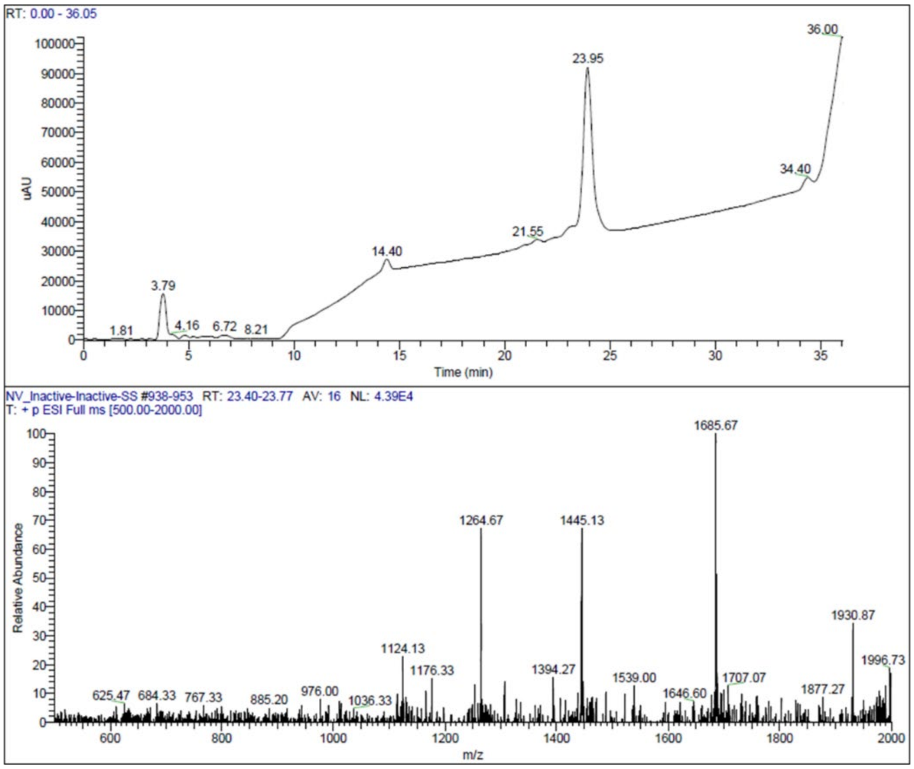

**Figure S27.** *LC/MS analysis of Inactive/Inactive FP*. Top panel is the HPLC chromatogram monitored by UV absorbance at 215nm. Bottom panel shows the ESI-MS spectra of positive ions. Due to the molecular weight and the m/z limit on the instrument (2000), the ions observed are 1685.67 (M+6), 1445.13 (M+7) and 1264.67 (M+8/8), which are less abundant, causing a lower signal-to-noise ratio.

**Table S1.** Identification of analogs

| Analog | Sequence | Modifications | Molecular Weight (g/mol) | Expected ions | Observed ions |
|---|---|---|---|---|---|
| glucagon | HSQGTFTSDYSKYLDSRRAQDFV QWLMNT | N/A | 3482.79 | 1742.39, 1161.93 | 1742.20, 1161.87 |
| dasiglucagon | HSQGTFTSDYSKYLDXARAEEFV KWLEST | See (44), X = α-aminoisobutyric acid | 3381.61 | 1691.80, 1128.20 | 1691.60. 1127.93 |
| glucagon-EEK | HSQGTFTSDYSOYLDSOOAQDFV QWLMNTEEK | K and R replaced to Orn EEK C-term extension | 3771.04 | 1886.52, 1258.01 | 1885.87, 1257.93 |
| $K^{13}$-$E^{17}$-linear | HSQGTFTSDYSOKLDSEOAQDFV QWLMNTEEK | $K^{13}$ and $E^{17}$ K and R replaced to Orn EEK C-term extension | 3751.01 | 1876.51, 1251.34 | 1876.20, 1251.13 |
| $K^{13}$-$E^{17}$-lactam (lc-glucagon) | HSQGTFTSDYSOKLDSEOAQDFV QWLMNTEEK | $K^{13}$-$E^{17}$ lactam bond K and R replaced to Orn EEK C-term extension | 3733.01 | 1867.51, 1245.34 | 1867.27, 1245.20 |
| $K^{17}$-$E^{21}$-linear | HSQGTFTSDYSOYLDSKOAQEFV QWLMNTEEK | $K^{17}$ and $E^{21}$ K and R replaced to Orn EEK C-term extension | 3799.1 | 1900.55, 1267.37 | 1900.27, 1267.13 |
| $K^{17}$-$E^{21}$-lactam | HSQGTFTSDYSOYLDSKOAQEFV QWLMNTEEK | $K^{17}$-$E^{21}$ lactam bond K and R replaced to Orn EEK C-term extension | 3781.10 | 1891.55, 1261.37 | 1891.20, 1261.27 |
| $K^{9}$-$E^{13}$-lactam (inactive) | HSQGTFTSKYSOELDSOOAQDFV QWLMNTEEK | $K^{9}$-$E^{13}$ lactam bond K and R replaced to Orn EEK C-term extension | 3732.07 | 1867.04, 1245.02 | 1866.73, 1244.93 |
| lispro insulin | FVNQHLCGSHLVEALYLVCGERG FFYTKPT- GIVEQCCTSICSLYQLENYCN | $Lys^{B28}$ and $Pro^{B29}$ | 5807.67 | 1936.89, 1452.92 | 1936.33, 1452.60 |
| SCI | FVNQHLCGSHLVEALYLVCGEOG FFYTPETEEGPOOGIVEQCCQSI CSLEQLENYCN | $Glu^{B29}$, $Gln^{A8}$, $Glu^{A14}$ R replaced to Orn EEGPOO linker | 6382.17 | 1596.54, 1277.43 | 1596.27, 1277.27 |
| inactive SCI | FVNQHLCGSHLVEALYLVCGEOG FFYTPETEEGPOOGILEQCCQSI CSLEQLENYCN | $Leu^{A3}$, $Glu^{B29}$, $Gln^{A8}$, $Glu^{A14}$ R replaced to Orn EEGPOO linker | 6396.20 | 1600.05, 1280.24 | 1599.87, 1279.80 |
| Active/Active FP | HSQGTFTSDYSOKLDSEOAQDFV QWLMNTEEKFVNQHLCGSHLVEA LYLVCGEOGFFYTPETEEGPOOG IVEQCCQSICSLEQLENYCN | lc-glucagon fused to SCI | 10097.16 | 1683.86, 1443.45 | 1683.6, 1443.07 |
| Active/Inactive FP | HSQGTFTSKYSOELDSOOAQDFV QWLMNTEEKFVNQHLCGSHLVEA LYLVCGEOGFFYTPETEEGPOOG IVEQCCQSICSLEQLENYCN | $K^{9}$-$E^{13}$-lactam fused to SCI | 10096.22 | 1683.70, 1443.32 | 1683.47, 1443.20 |
| Inactive/Active FP | HSQGTFTSDYSOKLDSEOAQDFV QWLMNTEEKFVNQHLCGSHLVEA LYLVCGEOGFFYTPETEEGPOOG ILEQCCQSICSLEQLENYCN | lc-glucagon fused to Inactive SCI | 10111.19 | 1686.20, 1445.46 | 1686.07, 1445.87 |
| Inactive/Inactive FP | HSQGTFTSKYSOELDSOOAQDFV QWLMNTEEKFVNQHLCGSHLVEA LYLVCGEOGFFYTPETEEGPOOG ILEQCCQSICSLEQLENYCN | $K^{9}$-$E^{13}$-lactam fused to Inactive SCI | 10110.25 | 1686.04, 1445.32 | 1685.67, 1445.13 |

**Table S2**. The $^{13}C_\alpha$ and $^{13}C_\beta$ chemical shift of lc-glucagon at pH 7.4 and at 25 °C[a]

| residue | | His1 | | Ser2 | | Gln3 | | Gly4 | | Thr5 | | Phe6 | | Thr7 | | Ser8 | |
|---|---|---|---|---|---|---|---|---|---|---|---|---|---|---|---|---|---|
| | | $^{13}C_\alpha$ | $^{13}C_\beta$ | $^{13}C_\alpha$ | $^{13}C_\beta$ | $^{13}C_\alpha$ | $^{13}C_\beta$ | $^{13}C_\alpha$ | | $^{13}C_\alpha$ | $^{13}C_\beta$ | $^{13}C_\alpha$ | $^{13}C_\beta$ | $^{13}C_\alpha$ | $^{13}C_\beta$ | $^{13}C_\alpha$ | $^{13}C_\beta$ |
| random coil shift[b] | | 55.78 | 29.62 | 58.35 | 63.88 | 55.94 | 28.67 | 45.34 | | 61.59 | 69.75 | 56.94 | 39.43 | 61.59 | 69.75 | 58.35 | 63.88 |
| free form | chemical shift | 56.56 | 33.10 | 58.11 | 63.61 | 56.09 | 29.40 | 45.27 | | 61.84 | 69.79 | 57.66 | 39.75 | 61.64 | 69.68 | 58.58 | 63.78 |
| | 2nd shift[c] | 0.78 | 3.48 | -0.24 | 0.80 | 0.15 | 0.73 | -0.07 | | 0.25 | 0.04 | 0.72 | 0.84 | 0.05 | -0.06 | 0.23 | -0.10 |
| fusion protein | chemical shift | 56.32 | 32.32 | 58.19 | 63.61 | 56.23 | 29.35 | 45.30 | | 61.84 | 69.82 | 57.83 | 39.75 | 61.75 | 69.74 | 58.75 | 63.77 |
| | 2nd shift[c] | 0.54 | 2,70 | -0.16 | 0.80 | 0.29 | 0.68 | -0.04 | | 0.25 | 0.07 | 0.89 | 0.84 | 0.16 | -0.01 | 0.40 | -0.11 |
| **residue** | | **Asp9** | | **Tyr10** | | **Ser11** | | **Orn12[d]** | | **Lys13[e]** | | **Leu14** | | **Asp15** | | **Ser16** | |
| | | $^{13}C_\alpha$ | $^{13}C_\beta$ | $^{13}C_\alpha$ | $^{13}C_\beta$ | $^{13}C_\alpha$ | $^{13}C_\beta$ | $^{13}C_\alpha$ | $^{13}C_\beta$ | $^{13}C_\alpha$ | $^{13}C_\beta$ | $^{13}C_\alpha$ | $^{13}C_\beta$ | $^{13}C_\alpha$ | $^{13}C_\beta$ | $^{13}C_\alpha$ | $^{13}C_\beta$ |
| random coil shift[b] | | 54.09 | 40.76 | 57.72 | 38.71 | 58.35 | 63.88 | 56.11 | 30.52 | 56.40 | 32.57 | 54.85 | 41.87 | 54.09 | 40.76 | 58.35 | 63.88 |
| free form | chemical shift | 54.46 | 41.05 | 58.74 | 38.30 | 59.69 | 63.44 | 56.84 | 30.48 | 57.18 | 32.78 | 55.97 | 42.37 | 55.68 | 41.13 | 58.57 | 63.09 |
| | 2nd shift[c] | 0.37 | 0.29 | 1.02 | -0.41 | 1.34 | -0.44 | 0.73 | -0.04 | 0.78 | 0.21 | 1.12 | 0.50 | 1.59 | 0.37 | 0.22 | -0.79 |
| fusion protein | chemical shift | 54.62 | 41.07 | 58.96 | 38.40 | 59.93 | 63.44 | 57.14 | \ | 58.59 | 32.67 | 56.25 | 42.32 | 55.78 | 41.09 | 58.63 | 63.06 |
| | 2nd shift[c] | 0.53 | 0.31 | 1.24 | -0.31 | 1.58 | -0.44 | 1.03 | \ | 1.41 | 0.10 | 1.40 | 0.45 | 1.69 | 0.33 | 0.28 | -0.82 |
| **residue** | | **Glu17[e]** | | **Orn18[d]** | | **Ala19** | | **Gln20** | | **Asp21** | | **Phe22** | | **Val23** | | **Gln24** | |
| | | $^{13}C_\alpha$ | $^{13}C_\beta$ | $^{13}C_\alpha$ | $^{13}C_\beta$ | $^{13}C_\alpha$ | $^{13}C_\beta$ | $^{13}C_\alpha$ | $^{13}C_\beta$ | $^{13}C_\alpha$ | $^{13}C_\beta$ | $^{13}C_\alpha$ | $^{13}C_\beta$ | $^{13}C_\alpha$ | $^{13}C_\beta$ | $^{13}C_\alpha$ | $^{13}C_\beta$ |
| random coil shift[b] | | 56.39 | 30.02 | 56.11 | 30.52 | 52.67 | 19.03 | 55.94 | 28.67 | 54.09 | 40.76 | 56.94 | 39.43 | 61.80 | 32.68 | 55.94 | 28.67 |
| free form | chemical shift | 56.50 | 29.48 | 56.07 | 30.49 | 53.14 | 19.05 | 56.19 | 29.37 | 54.69 | 41.04 | 58.42 | 39.51 | \ | 32.58 | 56.57 | 29.22 |
| | 2nd shift[c] | 0.11 | -0.54 | -0.04 | -0.03 | 0.47 | 0.02 | 0.25 | 0.70 | 0.60 | 0.28 | 1.48 | 0.08 | \ | -0.10 | 0.63 | 0.55 |
| fusion protein | chemical shift | 56.70 | 29.49 | 56.27 | 30.42 | 53.10 | 18.93 | 56.22 | 29.26 | 55.05 | 41.04 | 58.33 | 39.51 | \ | 32.51 | 56.34 | 29.28 |
| | 2nd shift[c] | 0.31 | -0.53 | 0.16 | -0.10 | 0.43 | -0.10 | 0.28 | 0.59 | 0.96 | 0.28 | 1.39 | 0.08 | \ | -0.17 | 0.40 | 0.61 |
| **residue** | | **Trp25** | | **Leu26** | | **Met27** | | **Asn28** | | **Thr29** | | **Glu30** | | **Glu31** | | **Lys32** | |
| | | $^{13}C_\alpha$ | $^{13}C_\beta$ | $^{13}C_\alpha$ | $^{13}C_\beta$ | $^{13}C_\alpha$ | $^{13}C_\beta$ | $^{13}C_\alpha$ | $^{13}C_\beta$ | $^{13}C_\alpha$ | $^{13}C_\beta$ | $^{13}C_\alpha$ | $^{13}C_\beta$ | $^{13}C_\alpha$ | $^{13}C_\beta$ | $^{13}C_\alpha$ | $^{13}C_\beta$ |
| random coil shift[b] | | 57.62 | 29.27 | 54.85 | 41.87 | 55.12 | 32.93 | 52.94 | 38.22 | 61.59 | 69.75 | 56.39 | 30.02 | 56.39 | 30.02 | 56.40 | 32.57 |
| free form | chemical shift | 57.78 | 29.33 | 55.39 | 42.48 | 55.77 | 32.93 | 53.38 | 38.90 | 61.96 | 69.85 | 55.56 | 30.42 | 55.57 | 30.40 | 57.46 | 33.88 |
| | 2nd shift[c] | 0.16 | 0.06 | 0.54 | 0.61 | 0.65 | 0.00 | 0.44 | 0.68 | 0.36 | 0.10 | 0.18 | 0.40 | 0.19 | 0.38 | 1.06 | 1.31 |
| fusion protein | chemical shift | 57.69 | 28.77 | 56.12 | 42.42 | 55.92 | 32.75 | 53.54 | 38.96 | 61.89 | 69.82 | \ | 30.50 | \ | \ | \ | \ |
| | 2nd shift[c] | 0.07 | -0.50 | 1.27 | 0.55 | 0.80 | -0.18 | 0.60 | 0.74 | 0.30 | 0.07 | \ | 0.48 | \ | \ | \ | \ |

[a]All chemical shifts were calibrated in parts per million (ppm) relative to 4,4-dimethyl-4-silapentane-1-sulfonic acid (DSS) as an internal standard, which was set to 0 ppm.

[b]The $^{13}C_\alpha$/$^{13}C_\beta$ chemical shift of random coil (43)

[c]The $^{13}C_\alpha$/$^{13}C_\beta$ secondary chemical shift was defined the difference between observed chemical shifts and random coil chemical shifts which generally were used to predict secondary structure of proteins (36-38).

[d]The random coil chemical shifts of $^{13}C_\alpha$ and $^{13}C_\beta$ resonance for ornithine residue were obtained from the $^{1}H$-$^{13}C$ HSQC spectrum of peptide GGOGG acquired at pH 7.4 (direct meter reding) and at 25 °C.

[e] Amino acid position of sidechain to sidechain lactam bond formation.

\ Chemical shift was not possible to assign.

**Table S3**. The $^{13}C_\alpha$ and $^{13}C_\beta$ chemical shift of A-chain residues of SCI at pH 7.4 and at 25°C[a]

| residue | | GlyA1 | | IleA2 | | ValA3 | | GluA4 | | GlnA5 | | CysA6 | | CysA7 | | GlnA8 | |
|---|---|---|---|---|---|---|---|---|---|---|---|---|---|---|---|---|---|
| | | $^{13}C_\alpha$ | | $^{13}C_\alpha$ | $^{13}C_\beta$ | $^{13}C_\alpha$ | $^{13}C_\beta$ | $^{13}C_\alpha$ | $^{13}C_\beta$ | $^{13}C_\alpha$ | $^{13}C_\beta$ | $^{13}C_\alpha$ | $^{13}C_\beta$ | $^{13}C_\alpha$ | $^{13}C_\beta$ | $^{13}C_\alpha$ | $^{13}C_\beta$ |
| random coil shift[b] | | 45.34 | | 60.64 | 29.62 | 61.80 | 32.68 | 56.39 | 30.02 | 55.94 | 28.67 | \ | \ | \ | \ | 55.94 | 28.67 |
| free form | chemical shift | 45.80 | | 63.07 | \ | 65.35 | 31.71 | 59.41 | 29.22 | 58.86 | \ | 54.49 | \ | 53.72 | 39.20 | \ | \ |
| | 2nd shift[c] | 0.46 | | 2.03 | \ | 3.55 | -0.97 | 3.02 | -0.80 | 2.92 | \ | \ | \ | \ | \ | \ | \ |
| fusion protein | chemical shift | 45.91 | | \ | \ | 65.37 | 31.80 | 59.09 | 29.30 | 58.86 | \ | \ | \ | \ | 39.02 | \ | \ |
| | 2nd shift[c] | 0.57 | | \ | \ | 3.57 | -0.88 | 2.70 | -0.68 | 2.92 | \ | \ | \ | \ | \ | \ | \ |

| residue | | SerA9 | | IleA10 | | CysA11 | | SerA12[c] | | LeuA13 | | GluA14 | | GlnA15 | | LeuA16 | |
|---|---|---|---|---|---|---|---|---|---|---|---|---|---|---|---|---|---|
| | | $^{13}C_\alpha$ | $^{13}C_\beta$ | $^{13}C_\alpha$ | $^{13}C_\beta$ | $^{13}C_\alpha$ | $^{13}C_\beta$ | $^{13}C_\alpha$ | $^{13}C_\beta$ | $^{13}C_\alpha$ | $^{13}C_\beta$ | $^{13}C_\alpha$ | $^{13}C_\beta$ | $^{13}C_\alpha$ | $^{13}C_\beta$ | $^{13}C_\alpha$ | $^{13}C_\beta$ |
| random coil shift[b] | | 58.35 | 63.88 | 60.64 | 29.62 | \ | \ | 58.35 | 63.88 | 54.85 | 41.87 | 56.39 | 30.02 | 55.94 | 28.67 | 54.85 | 41.87 |
| free form | chemical shift | 56.09 | 64.56 | 60.36 | \ | \ | \ | 56.51 | 65.60 | 58.64 | 41.31 | 59.83 | \ | 58.75 | 28.90 | 58.20 | 42.11 |
| | 2nd shift[b] | -2.26 | 0.68 | -0.28 | \ | \ | \ | -1.84 | 1.72 | 3.79 | -0.56 | 3.44 | \ | 2.81 | 0.23 | 3.35 | 0.24 |
| fusion protein | chemical shift | 56.22 | 64.36 | 60.00 | \ | \ | \ | 56.39 | 65.62 | 58.57 | 41.33 | 59.55 | \ | 58.78 | 29.06 | 58.28 | 42.14 |
| | 2nd shift[c] | -2.11 | 0.48 | -0.64 | \ | \ | \ | -1.96 | 1.74 | 3.72 | -0.54 | 3.16 | \ | 2.84 | 0.39 | 3.43 | 0.27 |

| residue | | GluA17 | | AsnA18 | | TyrA19 | | CysA20 | | AsnA21 | |
|---|---|---|---|---|---|---|---|---|---|---|---|
| | | $^{13}C_\alpha$ | $^{13}C_\beta$ | $^{13}C_\alpha$ | $^{13}C_\beta$ | $^{13}C_\alpha$ | $^{13}C_\beta$ | $^{13}C_\alpha$ | $^{13}C_\beta$ | $^{13}C_\alpha$ | $^{13}C_\beta$ |
| random coil shift[b] | | 56.09 | 30.02 | 52.94 | 38.22 | 57.72 | 38.71 | \ | \ | 52.94 | 38.22 |
| free form | chemical shift | 58.15 | 29.75 | 55.18 | 38.64 | 59.19 | 38.63 | 53.34 | 35.74 | 55.09 | 38.96 |
| | 2nd shift[b] | 2.06 | -0.27 | 2.24 | 0.42 | 1.47 | -0.08 | \ | \ | 2.15 | 0.74 |
| fusion protein | | 58.24 | 29.73 | 55.20 | 38.71 | 59.05 | 38.48 | 53.36 | 35.97 | 55.00 | 38.81 |
| | | 2.15 | -0.29 | 2.26 | 0.49 | 1.33 | -0.23 | \ | \ | 2.06 | 0.59 |

[a]All chemical shifts were calibrated in parts per million (ppm) relative to 4,4-dimethyl-4-silapentane-1-sulfonic acid (DSS) as an internal standard, which was set to 0 ppm.

[b]The $^{13}C_\alpha/^{13}C_\beta$ chemical shift of random coil was average value (43)

[c]The $^{13}C_\alpha/^{13}C_\beta$ secondary chemical shift was defined the difference between observed chemical shifts and random coil chemical shifts which generally were used to predict secondary structure of proteins (36-38).

[d]The random coil chemical shifts of $^{13}C_\alpha$ and $^{13}C_\beta$ resonance for ornithine residue were obtained from the $^{1}H$-$^{13}C$ HSQC spectrum of peptide GGOGG acquired at pH 7.4 (direct meter reding) and at 25 °C.

\ Chemical shift was not possible to assign.

**Table S4**. The $^{13}C_\alpha$ and $^{13}C_\beta$ chemical shift of B-chain residues of SCI at pH 7.4 and at 25°C[a]

| residue | | PheB1 | | ValB2 | | AsnB3 | | GlnB4 | | HisB5 | | LeuB6 | | CysB7 | | GlyB8 |
|---|---|---|---|---|---|---|---|---|---|---|---|---|---|---|---|---|
| | | $^{13}C_\alpha$ | $^{13}C_\beta$ | $^{13}C_\alpha$ | $^{13}C_\beta$ | $^{13}C_\alpha$ | $^{13}C_\beta$ | $^{13}C_\alpha$ | $^{13}C_\beta$ | $^{13}C_\alpha$ | $^{13}C_\beta$ | $^{13}C_\alpha$ | $^{13}C_\beta$ | $^{13}C_\alpha$ | $^{13}C_\beta$ | $^{13}C_\alpha$ |
| random coil shift[b] | | 56.94 | 39.43 | 61.80 | 32.68 | 52.94 | 38.22 | 55.94 | 28.67 | 55.78 | 29.62 | 54.85 | 41.87 | \ | \ | 45.34 |
| free form | chemical shift | 58.46 | 41.72 | 62.00 | 32.86 | 53.43 | 38.22 | 54.80 | 28.93 | 57.89 | 29.12 | 53.78 | 44.76 | 54.25 | 47.56 | 46.75 |
| free form | 2nd shift[c] | 1.52 | 1.19 | 1.20 | 0.18 | 0.49 | 0.00 | -1.14 | 0.26 | 2.11 | -0.50 | -1.07 | 2.89 | \ | \ | 1.41 |
| fusion protein | chemical shift | \ | 41.60 | \ | \ | 53.47 | 38.60 | 54.88 | 28.95 | \ | 29.20 | \ | 44.31 | \ | 47.35 | 46.73 |
| fusion protein | 2nd shift[c] | \ | 1.07 | \ | \ | 0.53 | 0.38 | -1.06 | 0.28 | \ | -0.42 | \ | 2.44 | \ | \ | 1.39 |

| residue | | SerB9 | | HisB10 | | LeuB11 | | ValB12 | | GluB13 | | AlaB14 | | LeuB15 | | TyrB16 | |
|---|---|---|---|---|---|---|---|---|---|---|---|---|---|---|---|---|---|
| | | $^{13}C_\alpha$ | $^{13}C_\beta$ | $^{13}C_\alpha$ | $^{13}C_\beta$ | $^{13}C_\alpha$ | $^{13}C_\beta$ | $^{13}C_\alpha$ | $^{13}C_\beta$ | $^{13}C_\alpha$ | $^{13}C_\beta$ | $^{13}C_\alpha$ | $^{13}C_\beta$ | $^{13}C_\alpha$ | $^{13}C_\beta$ | $^{13}C_\alpha$ | $^{13}C_\beta$ |
| random coil shift[b] | | 58.35 | 63.88 | 55.78 | 29.62 | 54.85 | 41.87 | 61.80 | 32.68 | 56.39 | 30.02 | 52.67 | 19.03 | 54.85 | 41.87 | 57.72 | 38.71 |
| free form | chemical shift | 61.18 | 62.30 | \ | 29.70 | 57.76 | 40.39 | 66.81 | 31.69 | 59.33 | 29.69 | 55.12 | 19.11 | 57.63 | 40.14 | 61.83 | 38.03 |
| free form | 2nd shift[c] | 2.83 | -1.58 | \ | 0.08 | 2.91 | -1.48 | 5.01 | -1.01 | 2.94 | -0.33 | 2.55 | -0.08 | 2.78 | -1.73 | 4.09 | -0.68 |
| fusion protein | chemical shift | 61.27 | 62.78 | \ | 30.00 | 57.70 | 40.54 | 66.79 | 31.64 | 59.37 | 29.42 | 55.15 | 18.93 | 57.69 | 40.22 | 61.69 | 38.08 |
| fusion protein | 2nd shift[c] | 2.92 | -1.10 | \ | 0.38 | 2.85 | -1.33 | 4.99 | -0.04 | 2.98 | -0.60 | 2.58 | -0.10 | 2.84 | -1.65 | 3.97 | -0.63 |

| residue | | LeuB17 | | ValB18 | | CysB19 | | GlyB20 | | GluB21 | | OrnB22[d] | | GlyB23 | | PheB24 | |
|---|---|---|---|---|---|---|---|---|---|---|---|---|---|---|---|---|---|
| | | $^{13}C_\alpha$ | $^{13}C_\beta$ | $^{13}C_\alpha$ | $^{13}C_\beta$ | $^{13}C_\alpha$ | $^{13}C_\beta$ | $^{13}C_\alpha$ | | $^{13}C_\alpha$ | $^{13}C_\beta$ | $^{13}C_\alpha$ | $^{13}C_\beta$ | $^{13}C_\alpha$ | | $^{13}C_\alpha$ | $^{13}C_\beta$ |
| random coil shift[b] | | 54.85 | 41.87 | 61.80 | 32.68 | \ | \ | 45.34 | | 56.39 | 30.02 | 56.11 | 30.52 | 45.34 | | 56.94 | 39.43 |
| free form | chemical shift | 58.04 | 42.22 | 65.52 | 32.81 | 53.83 | 36.74 | 46.63 | | 57.72 | 29.45 | \ | \ | 44.80 | | 55.97 | \ |
| free form | 2nd shift[c] | 3.19 | 0.35 | 3.72 | 0.13 | \ | \ | 1.29 | | 1.33 | -0.57 | \ | \ | -0.54 | | -0.97 | \ |
| fusion protein | chemical shift | 58.11 | 42.26 | 65.59 | 32.62 | 53.91 | 36.61 | 46.55 | | 57.79 | 29.54 | \ | \ | 44.83 | | 55.91 | \ |
| fusion protein | 2nd shift[c] | 3.26 | 0.39 | 3.77 | -0.06 | \ | \ | 1.21 | | 1.40 | -0.48 | \ | \ | -0.51 | | -1.03 | \ |

| residue | | PheB25 | | TyrB26 | | ThrB27 | | ProB28 | | GluB29 | | ThrB30 | |
|---|---|---|---|---|---|---|---|---|---|---|---|---|---|
| | | $^{13}C_\alpha$ | $^{13}C_\beta$ | $^{13}C_\alpha$ | $^{13}C_\beta$ | $^{13}C_\alpha$ | $^{13}C_\beta$ | $^{13}C_\alpha$ | $^{13}C_\beta$ | $^{13}C_\alpha$ | $^{13}C_\beta$ | $^{13}C_\alpha$ | $^{13}C_\beta$ |
| random coil shift[b] | | 56.94 | 39.43 | 57.72 | 38.71 | 61.59 | 69.75 | 63.53 | 31.87 | 56.39 | 30.02 | 61.59 | 69.75 |
| free form | chemical shift | 56.91 | \ | 58.41 | 40.14 | \ | 70.20 | 63.33 | 32.22 | 58.25 | 30.50 | 61.57 | 69.92 |
| free form | 2nd shift[c] | -0.03 | \ | 0.69 | 1.43 | \ | 0.45 | 0.00 | 0.35 | 1.86 | 0.38 | -0.02 | 0.17 |
| fusion protein | chemical shift | 56.89 | \ | 58.21 | 40.18 | \ | 70.14 | 63.34 | 32.24 | \ | \ | 61.59 | 69.91 |
| fusion protein | 2nd shift[c] | -0.05 | \ | 0.49 | 1.47 | \ | 0.39 | 0.01 | 0.37 | \ | \ | 0.00 | 0.16 |

[a]All chemical shifts were calibrated in parts per million (ppm) relative to 4,4-dimethyl-4-silapentane-1-sulfonic acid (DSS) as an internal standard, which was set to 0 ppm.

[b]The $^{13}C_\alpha$/$^{13}C_\beta$ chemical shift of random coil was average value (43)

[c]The $^{13}C_\alpha$/$^{13}C_\beta$ secondary chemical shift was defined the difference between observed chemical shifts and random coil chemical shifts which generally were used to predict secondary structure of proteins (36-38).

[d]The random coil chemical shifts of $^{13}C_\alpha$ and $^{13}C_\beta$ resonance for ornithine residue were obtained from the $^{1}H$-$^{13}C$ HSQC spectrum of peptide GGOGG acquired at pH 7.4 (direct meter reding) and at 25 °C.

\ Chemical shift was not possible to assign.

**Table S5.** CD characterization of fusion protein and individual moieties

| Analog | α-helix content (%)* | $\Delta G_U$† (kcal/mol) | m-value‡ (kcal/mol*M) | $C_{mid}$ (M) |
|---|---|---|---|---|
| lispro | 16.6 | 3.0 ± 0.1 | 0.6 ± 0.01 | 4.7 ± 0.1 |
| glucagon | 14.1 | N/A§ | N/A | N/A |
| lc-glucagon | 16.2 | N/A§ | N/A | N/A |
| SCI | 19.9 | 4.2 ± 0.1 | 0.8 ± 0.02 | 5.4 ± 0.1 |
| A/A FP | 13.8 | 4.5 ± 0.1 | 0.8 ± 0.03 | 5.7 ± 0.2 |

* α-helix content was calculated from normalized CD spectra using the SELCON-3 algorithm packaged with the CDPro spectral analysis software (4-6). Spectra was acquired with samples in in 50 mM KCl and 10 mM KPi (pH 7.4) at 25 °C.

† *Parameters were inferred from CD-detected guanidine denaturation data by application of a two-state model to the molar ellipticity (θ) at 222 nm data; uncertainties represent fitting errors for a given dataset. Assays were run in 50 mM KCl and 10 mM KPi (pH 7.4) at 25 °C.

‡ The m-value [slope Δ(G)/Δ(M)] correlates with extent of hydrophobic surfaces exposed on denaturation.

§ Glucagon and lc-glucagon did not display a denaturation that followed a two-state model and therefore the fit was not possible.

**Table S6.** In vitro characterization of additional lactam (cyclic) and linear glucagon analogs

| Analog | lag time* ± SEM (hours) | cAMP $EC_{50}$† (nM) [95% CI] |
|---|---|---|
| glucagon | 0 | 163 [134, 198] |
| glucagon-EEK | 32 ± 2 | 389 [342, 442] |
| lc-glucagon | > 672 | 813 [671, 988] |
| $K^{12}$-$E^{16}$-linear | 478 ± 67 | 860 [672, 1102] |
| $K^{12}$-$E^{16}$-lactam | 558 ± 76 | 1750 [1191, 2368] |
| $K^{16}$-$E^{20}$-linear | 310 ± 33 | 463 [387, 577] |
| $K^{16}$-$E^{20}$-lactam | > 672 | 258 [135, 514] |
| $K^{20}$-$E^{24}$-linear | 570 ± 65 | 911 [707, 1172] |
| $K^{20}$-$E^{24}$-lactam | > 672 | 437 [358, 546] |
| $K^{24}$-$E^{28}$-linear | 514 ± 77 | 1020 [672, 1573] |
| $K^{24}$-$E^{28}$-lactam | > 672 | 403 [248, 666] |

* Fibrillation lag times were measured at 100μM in phosphate-buffered saline (pH 7.4). Samples were agitated via continuous shaking at 37°C. Time of initial fluorescence, defined as a 5-fold increase over baseline in ThT fluorescence, provided a criterion for onset of fibrillation.

† Increase in cAMP production on a HEK293 cell line stably overexpressing glucagon receptor was used to measure in vitro glucagon activity. Dose-response curve was fitted to a nonlinear regression equation to obtain $EC_{50}$ values.
